# Low-invasive sampling method for taxonomic for the identification of archaeological and paleontological bones by proteomics of their collagens

**DOI:** 10.1101/2023.10.18.562897

**Authors:** Isabelle Fabrizi, Stéphanie Flament, Claire Delhon, Lionel Gourichon, Manon Vuillien, Tarek Oueslati, Patrick Auguste, Christian Rolando, Fabrice Bray

## Abstract

Collagen from paleontological bones is an important organic material for isotopic measurement, radiocarbon and paleoproteomic analyzes, to provide information on diet, dating and taxonomy. Current paleoproteomics methods are destructive and require from a few milligrams to several tenths of milligrams of bone for analysis. In many cultures, bones are raw materials for artefact which are conserved in museum which hampers to damage these precious objects during sampling. Here, we describe a low-invasive sampling method that identifies collagen, taxonomy and post-translational modifications from Holocene and Upper Pleistocene bones dated to 130,000 and 150 BC using dermatological skin tape-discs for sampling. The sampled bone micro-powders were digested following our highly optimized eFASP protocol, then analyzed by MALDI FTICR MS and LC-MS/MS for identifying the genus taxa of the bones. We show that this low-invasive sampling does not deteriorate the bones and achieves results similar to those obtained by more destructive sampling. Moreover, this sampling method can be performed at archaeological sites or in museums.

## INTRODUCTION

Collagen is an extremely important material for revealing the past of bones.^1^ But sample preparation remains one of the most difficult aspects of paleoproteomic experiments, as to access the proteome removing mineral part of the bone made from hydroxyapatite requires special steps while preserving proteins, which is not necessary for other tissue types. Collagen isolation methods are destructive and consumes from a few milligrams to several tenths of milligrams of bone. The bone mineral must be eliminated by acid or basic demineralization and the residual collagen gelatinized, before enzymatic digestion and peptide analysis. In paleoproteomics, several methods of preparation have been introduced recently such as in solution digestion after proteins extraction, FASP method (Filter Aided Samples Preparation), eFASP (enhanced Filter Aided Samples Preparation) method or an approach based on paramagnetic beads with surfaces modified by resin bearing carboxylate group called SP3 (<u>S</u>ingle-pot, <u>S</u>olid-phase-enhanced <u>S</u>ample-Preparation) and “Species by Proteome INvestigation” (SPIN), a shotgun proteomics workflow for analysing archaeological bone.^2–8^ However, the destructive nature of these methods is undesirable when analyzing archaeological material, such as bone remains or rare artifacts made from bone. Several important considerations must be taken into account before destructive sampling of an artefact is carried out, such as the probability of successful analysis, the choice of sampling technique to minimize traces on the object, the amount of material to be sampled, and the effects of current sampling on future research and artifacts conservation.^9^

The development of a method of species identification that does not damage the object or leaves very little visible trace on the object and which can be used for the analysis of museum collections is still a challenge despite the facts that several methods has been described recently. One of the first developed method is ammonium bicarbonate buffer extraction without demineralization which was performed on bones to identifying taxa, even if the cold acid demineralization allows a better yield during the extraction of proteins.^10^ A second method which allows non-destructive analysis of samples is based on the triboelectric effect of a (PVC) eraser.^11^ This method was originally set up for the analysis of parchment and on other archaeological materials such as bone and ivory.^12–15^ The triboelectric effect has been applied to bones contained in plastic storage bags allowing identifying them.^16^ In 2019, Kirby *et al*. used polishing films to sample photographs.^17–18^ Then Zara Evans *et al.* applied this technique for the taxonomic identification of bone artefacts, with successful results on 5,000-year-old remains.^19^ More recently, proteins on skin and bones on the surfaces of cranial bone of a mummified Egyptian from the 26^th^ Dynasty (664–525 BC) has been identified using dermatology grade skin sampling strips.^20^ We describe here that low-invasive proteomics identification of Iron Age, Neolithic and Upper Pleistocene bones from 120,000-150 BC based on sampling with dermatological skin tape discs. Our new sensitive digestion method based on 96 well plate demineralization and digestion and MALDI FTICR MS analysis has a sensitivity below the milligram of bones. We first compare dermatological skin tape discs sampling with previously describe minimally invasive methods using our optimized protocol. Then we show that dermatological skin tape discs sampling method allows sufficient collagen to be obtained for correct taxonomic identification and does not affect the appearance of the bone and can be used in archaeological, paleontological or museum sites.

## EXPERIMENTAL SECTION

### Chemicals and biochemicals

All aqueous solutions were prepared from ultrapure grade water obtained by water filtration with a two stages Millipore system (Milli-Q® Academic with a cartouches Q-Gard 1 and Progard 2, Merck Millipore, Burlington, Massachusetts, United States). All chemicals, biochemicals and solvents were purchased from Merck (Merck KGaA, Darmstadt, Germany) and used without purification. All solvents were MS analytical grade.

### Samples

Neolithic samples came from Tremblay-en-France (France) (260 – 150 BC), Bouchain (Nord, France) (3200-2900 cal. BC) and the cave of Pertus II (Alpes-de-Haute-Provence, France) (3800-3300 cal. BC).^21–24^ Upper Pleistocene samples came from the Sarrasins cave (Loverval, Belgium) (63 ka BC) and the Waziers (Nord, France) (132 – 123 ± 8 ka BC).^25^ Modern samples came from the laboratories (EEP and HALMA) of the authors specialized in archaeology or paleontology. The samples Mod 1, 3 and 4 were cleaned after maceration with water at 35°C. The sample Mod 2 were cleaned after maceration with water and detergent at 35°C then were bleached with hydrogen peroxide. All information on bones are given in Supplementary Information 1, Table S1 and the photos of bones are in Supplementary Information 1, Figure S1, S2, S3

### Triboelectric rubbing in bag sampling

The sampling method is based on the protocol described by McGrath *et al*.^12^ First, entire bone artifacts were placed in a clean, labeled Minigrip 60 µm, 100 × 150 mm (PlanetArcheo, Marcilly-le-Châtel, France) sample bags made of neutral low density polyethylene. The bone artifacts were rubbed gently for 60 s in the sealed bag to create triboelectric friction between the bag and the bone artifact. Bone artifacts were removed from the bag and stored in new, sterile sample bags. 300 μL of warmed AmBic (65 ℃) was pipetted into the empty sample bag and massaged gently for two minutes. The AmBic solution was extracted from the bag and placed in a 1.5 mL Eppendorf™ tubes. In solution trypsin digestion was then applied following the protocol described below.

### Eraser sampling by rubbing

An area of 1 cm² from bone artifacts were rubbed with a small polyvinyl chloride (PVC) eraser (Staedtler, UGAP, France) on a flat section of the artifact’s surface as described in McGrath *et al.*^12^ If the outer surface of the artifact was dirty, the initial eraser shavings were discarded and the area was erased again with a clean section of the eraser. Approximately, 15 mg of eraser shavings were collected and placed into 200 μL Ambic solution. Samples were vortexed at a low speed for 5 min (Vortex genie® 2, Scientific Industries, Inc., USA) and the solution was placed in a new 1.5 mL Eppendorf™ tubes. Gelatinization and in solution trypsin digestion were then applied following the protocol described below.

### Swab mopping sampling

An area of 1 cm² from bone artifacts was rubbed with a small swab (Foamtec UltraSOLV™ 1700 Series Swabs, Merck Life Science S.A.S, France) on a flat section of the artifact’s surface. A different zone was used for each method involving rubbing. The swab was placed in an Eppendorf™ 5 mL tube and 300 µL of AmBic solution were added. Swab was vortexed at a low speed for 5 min and the solution was placed in a new 1.5 mL Eppendorf™ tubes. Gelatinization and in solution trypsin digestion were then applied following the protocol described below.

### Ammonium bicarbonate buffer etching sampling

This method was based on Van Doorm *et al.*^10^ A piece of bone weighing approximately 50 mg was placed in 1.5 mL Eppendorf™ tubes and covered with AmBic solution. Gelatinization and in solution trypsin digestion were then applied following the protocol described below with the piece of bone in the Eppendorf.

### Conventionnal destructive sampling

This method was based on Bray *et al.*^26^ Approximately 1 mg of bone was removed by scraping with a scalpel blade and demineralized in 100 μL of 5% TFA solution during 4 h at room temperature with vortexing. The demineralization solution was recovered, 8 µL NaOH 6 M was added to neutralize TFA and the 100 µL of 100 mM AmBic. In solution trypsin digestion was then applied. The bone powder was rinsed with 200 μL of 50 mM AmBic and this step was repeated twice. Then 200 μL AmBic was added on bone powder. Gelatinization and in solution trypsin digestion were then applied following the protocol described below.

### Dermatological skin tape-discs sampling on Iron Age, Neolithic and Upper Pleistocene bones

Sample preparation was conducted according to established guidelines for ancient protein work, to minimize exogenous laboratory contamination.^1^ The surface of the bones where the samples were going to be taken was first cleaned with a brush to remove post-depositional deposits. Then the area was cleaned with a wipe soaked with mQ water (Kimberly Clark Kimtech Science, Nanterre, France). The samples were taken after the surface had dried. Each tape-disk (D-Squame®, 22 mm diameter, 3.8 cm^2^, Monaderm, Principality of Monaco) was pressed manually against the bone for 3 s, before being stripped from the bone. For experiments involving several samplings, each tape-disk was placed at the same place on the bone. Tape-disks were cut into quarters and the four pieces were placed in individual 1.5 mL Eppendorf™ (Eppendorf, Hamburg, Germany). Extraction of bone particles was realized by adding 500 μL of extraction buffer (8 M urea, 100 mM ammonium bicarbonate pH 8.8), followed by 15 min sonification in iced water (0– 4°C) in an ultrasonic bath (Advantage Lab, Switzerland) followed by 1 h on vortex at 4 °C. The tape-disk quarters were scraped in the Eppendorf with a spatula to release all bone particles in the solution and the tape-disk quarters were discarded. The solutions containing the bone particles were combined in a 5 mL Eppendorf™, then concentrated with a 0.5 mL Amicon® ultra centrifugal filters with a cut-off of 10 kDa (EMD Millipore, Darmstadt, Germany) by centrifugation for 20 min at 10,000 g using Eppendorf™ centrifuge 5430R (Eppendorf, Hambourg, Germany). Before use, 0.5 mL Amicon® ultra centrifugal filters with a cut-off of 10 kDa were freshly incubated overnight with the passivation solution containing 5% (v/v) Tween®-20 were washed in a water bath during 20 min four times. After concentration, 100 µL of denaturation buffer (8 M urea, 50 mM DTT, 100 mM ammonium bicarbonate pH 8.8) were added and the Amicon® was incubated at 4 °C overnight.

### eFASP digestion optimized for skin tape-disc sampling

The proteomic method is based on the protocol described by Bray *et al.* which has been optimized for skin tape-disc sampling.^7^ The bone powder suspension in the Amicon® obtained in the previous step was centrifuged during 20 min at 10,000 g, then washed with 100 µL of exchange buffer (8 M urea, 100 mM ammonium bicarbonate pH 8.8) and then the exchange buffer was eliminated by centrifugation during 20 min at 10,000 g. The filtrates were discarded, then 200 µL of exchange buffer were added again into the Amicon® filter which was centrifuged. This step was repeated twice. Proteins were alkylated during 1 h at room temperature in the dark using 100 µL of alkylation buffer (8 M urea, 50 mM iodoacetamide, and 100 mM ammonium bicarbonate, pH 8.8). The Amicon® filter was centrifuged for 20 min at 10,000 g and the filtrate was discarded. After the alkylation step, 200 µl of exchange buffer was added to the Amicon® filter which was centrifuged for 20 min at 10,000 g and the filtrate was discarded. 200 µl of AmBic solution (50 mM ammonium bicarbonate pH 8.8) were added to the Amicon® filter and then centrifuged. This step was repeated twice, discarding the filtrate at each step. The Amicon® filter was transferred to new 2 mL tubes. 100 µL of AmBic solution and 40 μl of trypsin (0.05 µg/µl, Promega, Madison, USA) were added and incubated with shaking at 400 rpm in a heating block tube (MHR23, Hettich, Netherlands) overnight at 37 °C. After this step, the peptides present in the Amicon® filter were recovered in the lower tube by centrifugation during 15 min at 10,000 g. In order to obtain a maximum of peptides, the Amicon® filters was washed twice with 50 µl of AmBic solution. The filtrates containing all the peptides were transferred to 1.5 mL Eppendorf™ tubes and were evaporated to dryness at room temperature with a SpeedVac™ Concentrator (Eppendorf, Hamburg, Germany). Tryptic peptides were desalted on 96-well plates C18 (Affinisep, Petit-Couronne, France) following the protocol described below.

### Gelatinization and in solution trypsin digestion

Following individual sampling procedures, all samples were gelatinized, except the demineralization solution from the destructive sampling, in AmBic at 65 ℃ for one hour with shaking at 400 rpm in a heating block tube. Samples were incubated overnight (12–18 h) at 37 ℃ with 10 µL of trypsin solution (0.05 µg / µl in 50 mM AmBic solution) with shaking at 400 rpm on a heating stirrer MHR23 (Hettich, Tuttlingen, Germany). After digestion, 300 µL of 0.5% acetic acid were added and peptides were purified and desalted using 96-well plate C18.

### Purification of peptides

Briefly, the plate was washed once with 500 µL of acetonitrile (ACN) followed by a washing step with 80% ACN, H_2_O 0.5% acetic acid repeated 3 times and a second washing repeated 3 times with H_2_O alone 0.5% acetic acid. Tryptic peptides from bone powder were resuspended in 200 µL of a H_2_O, 0.5% acetic acid solution. Tryptic peptides form both solutions were transferred to C18 96-well plate and eluted with a vacuum manifold. The plate was washed 3 times with 200 µL of H_2_O, 0.5% acetic acid. Peptides were recovered in a V-bottom well collecting plate using 100 µL of a 80% ACN, 0.1% acetic acid solution followed by 100 µL of ACN. The plate was evaporated on TurboVap 96 Evaporator (Caliper LifeScience, Hopkinton, USA). For mass spectrometry analysis, the sample was dissolved again in 10 μl of solvent A of LC (see below). The concentration was then estimated by measuring the OD at 215 nm using 1 μl of the solution using a droplet UV spectrometer (DS-11+, Denovix, Wilmington, USA). Samples were diluted at a concentration of 1 µg/µL before LC-MS/MS analysis.

### Liquid chromatography-tandem mass spectrometry

LC-MS/MS analyses were performed on an Orbitrap Q Exactive plus mass spectrometer hyphenated to a U3000 RSLC Microfluidic HPLC System (ThermoFisher Scientific, Waltham, Massachusetts, USA). 1 μl of the peptide mixture at a concentration of 1 µg/µL was injected with solvent A (5% acetonitrile and 0.1% formic acid v/v) for 3 min at a flow rate of 10 μl.min-1 on an Acclaim PepMap100 C18 pre-column (5 μm, 300 μm i.d. × 5 mm) from ThermoFisher Scientific. The peptides were then separated on a C18 Acclaim PepMap100 C18 reversed phase column (3 μm, 75 mm i.d. × 500 mm), using a linear gradient (5-40%) of solution B (75% acetonitrile and 0.1% formic acid) at a rate of 250 nL.min-1. The column was washed with 100% of solution B during 5 minutes and then re-equilibrated with buffer A. The column and the pre-column were placed in an oven at a temperature of 45°C. The total duration of the analysis was 140 min. The LC runs were acquired in positive ion mode with MS scans from *m/z* 350 to 1,500 in the Orbitrap mass analyzer at 70,000 resolution at *m/z* 200. The automatic gain control was set at 1e10^6^. Sequentially MS/MS scans were acquired in the high-energy collision dissociation cell for the 15 most-intense ions detected in the full MS survey scan. Automatic gain control was set at 5e10^5^, and the normalized collision energy was set to 28 eV. Dynamic exclusion was set at 90 s and ions with 1 and more than 8 charges were excluded.

### MALDI FTICR analysis

Desalted peptides (1 µL) were deposited on 384 ground steel MALDI plates (Bruker Daltonics, Bremen, Germany), then 1 µL of HCCA matrix at 10 mg/mL in ACN/H_2_O 70:30 v/v 0.1% TFA was added for each sample spot and dried at room temperature. MALDI FTICR experiments were carried out on a Bruker 9.4 Tesla SolariX XR FTICR mass spectrometer controlled by FTMS Control software and equipped with a CombiSource and a ParaCell (Bruker Daltonics, Bremen, Germany). A Bruker Smartbeam-II Laser System was used for irradiation at a frequency of 1,000 Hz using the “Minimum” predefined shot pattern. MALDI FTICR spectra were generated from 500 laser shots in the *m/z* range from 693.01 to 5,000. 2M data points were used per spectrum which corresponds to a transient duration of 5.0332 s. Twenty spectra were averaged. The transfer time to the ICR cell was set to 1.2 ms and the quadrupole mass filter set at *m/z* 600 was operated in RF-only mode.

### Bioinformatics

MS raw data from MALDI FTICR were processed using DataAnalysis 5.0. For deisotoping and extracting the monoistopic peaks the SNAP algorithm was employed with the following parameters of S/N > 3 and quality 0.6. The procedure for the deamidation value calculation from MALDI FTICR was based on Bray *et a,l* 2023.^26^ The identification is supported by all peptide markers presented in previous reports.^27–32^

Proteomics data were processed with Mascot v 2.5.1 against NCBI database mammalian (NCBI_2022_01, 9,016,701 sequences) and Aves (NCBI_2022_01, 2,494,584,292 sequences). Three missed cleavages, 10 ppm mass error for MS and MS/MS were applied. Cysteine carbamidomethylation (+57.02 Da) was set as fixed modification. Methionine oxidation (+15.99 Da) and asparagine, glutamine deamidation (+0.98 Da) were selected as variable modifications.

The second bioinformatics analysis was performed with PEAKS X plus (Bioinformatics software, Waterloo, Canada) against a home-made database containing 1,765 collagen sequences extracted from NCBI database (All_Collagen, from Bray *et al* 2023) restricted to Mammalian. ^26^ Precursor’s mass tolerance was fixed to 10 ppm and fragment ion mass tolerance to 0.05 Da. Three missed cleavages were allowed. The same post-translational modifications (PTMs) above were allowed plus hydroxylation of amino acids (RYFPNKD) (+15.99 Da) as variable modifications. Five variable PTMs were allowed per peptides. PEAKS PTM and SPIDER ran with the same parameters. Results were filtered using the following criteria: protein score –10logP ≥ 20’, 1% peptide False Discovery Rate (FDR)’, PTM with Ascore = 20’, mutation ion intensity = 5% and Denovo ALC ≥ 50%. Peptides with amino acids substitutions was filtered with minimal intensity set as 1e107. Peptides identified on collagen I alpha 1 and I alpha 2 were aligned against the NCBI non redundant protein sequence (all non-redundant GenBank CDS translations+PDB+SwissProt+PIR+PRF excluding environmental samples from WGS projects containing 308’,570’,119 sequences) to find similarity with BLASTp (https://blast.ncbi.nlm.nih.gov/Blast.cgi?PAGE=Proteins). The scoring parameter alignment used BLOSUM62 matrix.^33^ The specific peptides were validated with a score of 100% of identity and full query coverage.

The mass spectrometry proteomics data have been deposited on the ProteomeXchange Consortium (http://proteomecentral.proteomexchange.org) via the PRIDE partner repository with the data set identifier PXD044039 (Private data, Username, Password: WzZDnHxu).^34^

### Calculation of deamidation and hydroxyproline level

MS raw data were also processed by MaxQuant (MQ) software 1.5.8.3 (https://www.maxquant.org/) to further investigate about deamidation level of the proteins identified.^35^ In this search, for each identified species specific COL1A1 and COL1A2 sequences from Uniprot and NCBi were used.^26^ Database search was carried out using the following parameters: (i) full tryptic peptides with a maximum of 3 missed cleavage sites; (ii) cysteine carbamidomethylation as a fixed modification and (iii) oxidation of methionine, the transformation of N-terminal glutamine and N-terminal glutamic acid residue to pyroglutamic acid, hydroxyproline oxidation, and the deamidation of asparagine and glutamine as variable modifications. Match type was “match from and to” and the decoy mode was set to “revert”. PSM (Peptide-Spectrum Matches) and Protein and Site decoy fraction False Discovery Rate (FDR) were set at 0.01 as threshold for peptide and protein identifications. Match between run was used with 2 min for match time windows and 20 min for alignment time windows. Dependent peptides FDR was set at 0.01 All the other parameters were set as default.

An estimation of the percentage of deamidation for N and Q residues for each sample was calculated using the freely available command-line script “deamidation” (https://github.com/dblyon/deamidation), which use the MaxQuant “evidence.txt” file. The calculations were done separately for potentially original peptides and potential contaminants peptides as previously reported in Mackie *et al.* 2018.^36^

For the determination of deamidation following the traditional ZooMS approach COL1A1 peptides named P1^m^ (position COL1A1-508-519) with the sequence GVQ^dem^GP^ox^PGPAGPR for mammals was used.^27, 37^ The peptide sequence GVQ^dem^GP^ox^PGPQGPR named P1^b^ with the same position on COL1A1 compared to mammals and wich appears conserved across all birds, Australian marsupials and some reptiles was used.^38^ In MALDI FTICR the percentage of deamidation of the peptide from COL1A1 is calculated simply by dividing the intensity of the monoisotopic peak of the deamidated peptide by the sum of the intensity of the monoisotopic peak of the non-deamidated and that of the deamidated peptide without any assumption. In LC-MS/MS analyses without database search the same percentage is calculated from the MS spectrum of the relevant peptides identified by their MS/MS spectra and retention times. The MS/MS spectra are given in Supplementary Information S3, Figure S1-S4. The *m/z* of the peptides with and without deamidation used for the calculation of deamidation percentage are presented in Supplementary Information 1, Table S3 which contains the masses expected in the MALDI FTICR and LC-MS/MS analyses for native and deamidated peptides An estimation of the percentage of hydoxyproline residues for each sample was calculated with PEAKS X+ PTM results. The calculation method is the same as for the calculation of deamidation on the COL1A1 peptide. The percentage of hydroxyproline residues is calculated for each peptide from COL1A1 and COL1A2 of the species identified then the average is calculated.

## RESULTS AND DISCUSSION

### Bone surface modification induced by the different samplings

Dermatological skin tape-disc sampling was first tested on a fragment of mandible from modern *Bos taurus* (Figure 1) to see the effect of the sampling on bone surface. The skin tape-disc measures 22 mm in diameter. It is placed on the surface of the cleaned bone and manually pressure is applied to the patch for 3 seconds. Then the skin tape-disc is gently removed using clamps. Figure 1, A shows a photograph of *Bos taurus* mandible with the dermatological skin tape-discs attached on the surface and Figure 1, B after removing it. When several patches are used the following patches are placed at the same place to avoid affecting a large area of bone.

**Figure 1.**
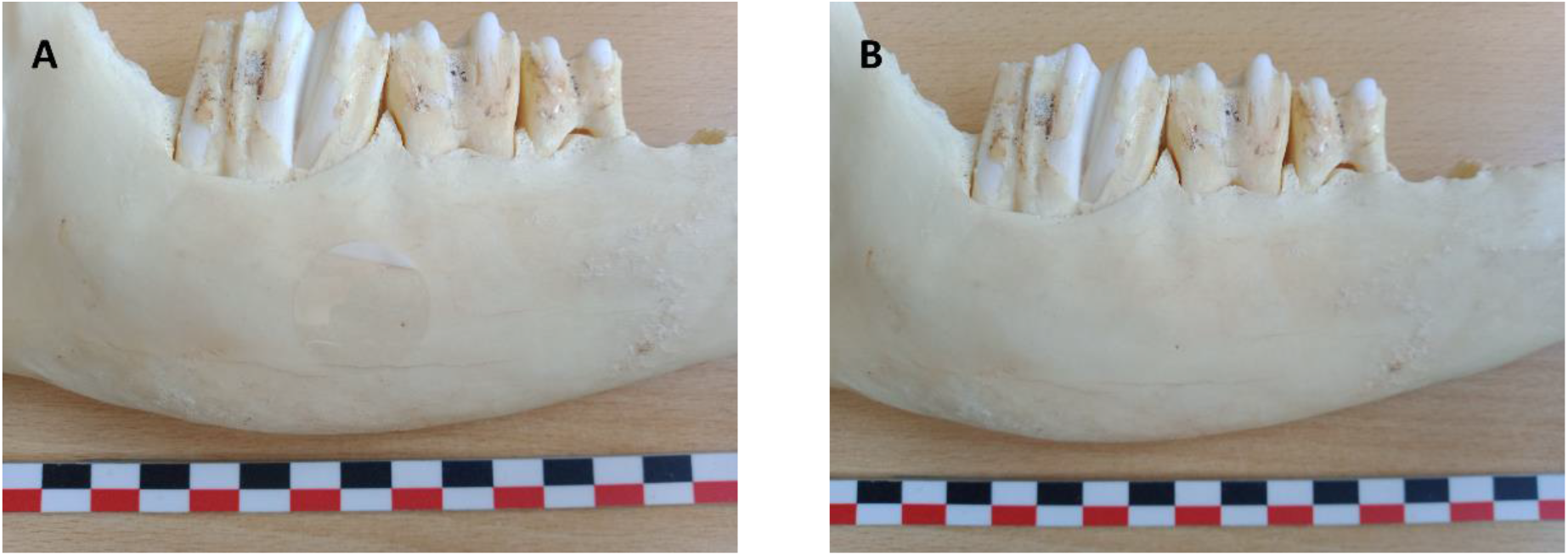
Image of modern *Bos taurus* mandible (Mod 1) with the dermatological skin tape-discs attached to the surface after pressing it (A) and after stripping (B). The black, white and red marks on the scale bar have a length of 1 cm.

The classical methods of sampling bones in archaeology and paleontology is performed conventionally by destructive sampling with a miniaturized grinder or a scalpel under conditions that avoid contaminations.^39^ However several low invasive samplings have been introduced recently (i) the triboelectric rubbing in bag sampling, (ii) the PVC eraser sampling by rubbing, (iii) the swab mopping sampling and (iv) the ammonium bicarbonate buffer etching sampling. We compared the effect on the surface of bones of the low invasive samplings except ammonium bicarbonate buffer etching which works on small fragments and of the conventional destructive method, with dermatological skin tape-disc stripping (Supplementary Information 1), There is no effect on the surface of the bone by methods triboelectric rubbing and PVC eraser sampling (Supplementary Information 1, Figure S4 A, B, C, D). A whitish deposit is observed after mopping by a swab (Supplementary Information 1, Figure S4 E, F). Figure S4, G, H shows the effect of sampling with scapel blade on the surface of the bone. The structure of the bone is not affected by stripping with 1 or 5 dermatological skin tape-disc, but after stripping with 10 and 20 dermatological skin tape-discs, the appearance of the bone surface is modified (Supplementary Information 1, Figure S4, I, J, K, L). The amount of material removed by our low sampling method is very small but it can leave a mark on the bone such as Virginie Sinet-Mathiot *at al.* showed in her study on the PVC eraser method, even if in our study, the PVC eraser method left no trace, but this depends on the state of conservation of the bone.^14^ It is important to take samples from areas where there is no trace of fracturing, cutting to avoid erasing traces of human activity. Supplementary Information 1, Figure S5, show the thin layer on the dermatological skin tape-disc after the first stripping and the powder obtained after scraping of the tape-disk which corresponding to 1 mg of bone powder. C. M. Oloesen *et al*. showed that the use of an adhesive strip tape removes 1.3 µm of epidermis by stripping.^40^

We tested next our methodology on a Neolithic bone. Supplementary 1, Figure S6, shows the effect of the sampling method on wild boar tibia from the Bouchain site (Bou 1). The image shows no effect on the surface of the archeological bone after triboelectric rubbing in forced bag sampling and eraser sampling by rubbing (Supplementary 1, Figure S6, A, B, C, D). With the swab methods, a slight abrasion was observed (Supplementary 1, Figure S6, E, F). With the Conventional destructive sampling, the sampling location is visible (Supplementary 1, Figure S6, G, H). The dermatological skin tape-discs sampling leaves no visible trace on the surface of the bone (Supplementary1, Figure S6, I, J).

### Optimization of the method based on dermatological skin tape-disc stripping

During sampling, each dermatological skin tape-disc cut in four part was deposited in a 1.5 mL Eppendorf® so that the extraction buffer covers the disc quarters to recover the bone powder. Each skin tape-discs was digested and analyzed separately, but also by combining them. When the stripping was performed with 5 dermatological skin tape-disc each disc was extracted separately and combined. The eFASP method was used to avoid the loss of bone powder present on the skin tape-discs and increase the numbers of identified peptides, proteins and coverage of proteins. The Figure 2, shows the spectrum of the digestion of the combined 5 dermatological skin tape-discs for modern *Bos taurus*. Specific ZooMS peptides markers are indicated. The resolution of the peptide P1^m^ at *m/z* 1105.574 was 315,000 identified in mammals samples.

**Figure 2.**
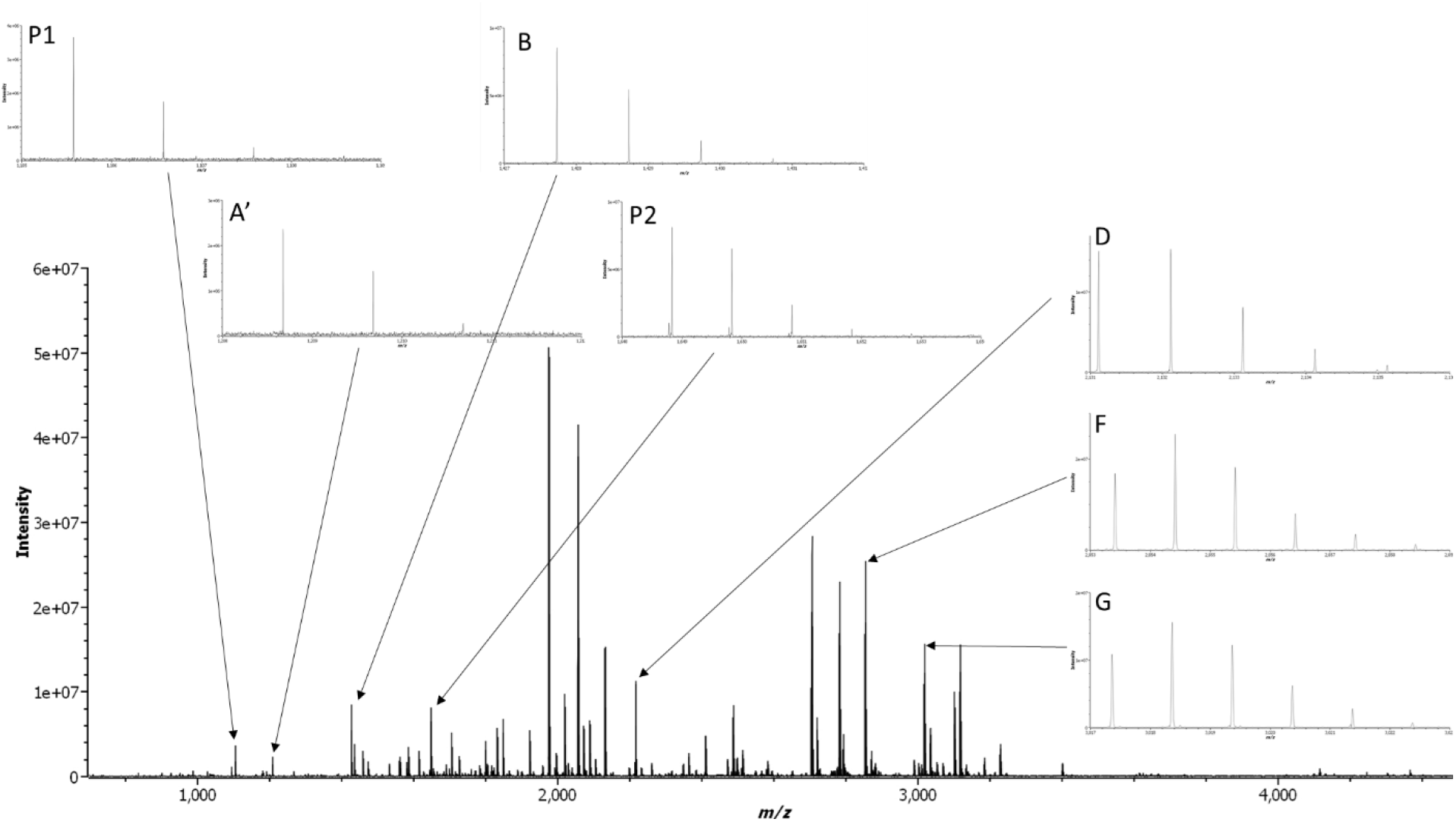
MALDI FTICR spectra of *Bos taurus* bone (Mod 1) tryptic digest using 5 dermatological skin tape-discs. Zoom on 6 ZooMS peptide markers (P1, A’, B, P2, D, F, G).

Each spectrum obtained from the separated and combined 5 dermatological skin tape-discs allowed to identify the genus Bos with the identification of the 12 ZooMS markers peptides (Supplementary Information 4). 1 and 5 combined dermatological skin tape-discs have been used for comparison with other destructive and non-destructive methods.

### Comparison with other destructive and non-destructive methods

We compared dermatological sampling with 1 and 5 skin tape-discs and the 5 other methods (i) triboelectric rubbing in forced bag, (ii) rubbing with a PVC eraser, (iii) mopping with swab, (iv) etching with ammonium bicarbonate buffer, and (v) conventional sampling by the destructive method taking 1 mg of bone on 4 modern samples and 3 archaeological samples. For the destructive method, 2 fractions were analyzed, the acid demineralization fraction and the bone powder one. The amount of peptide obtained after digestion evaluated by the absorbance at 215 nm is indicated in Supplementary Information 1, Table S2. Entries show that increasing the number of stripping with dermatological skin tape-discs increased the amount of peptide after digestion. In the literature But Dylan H. Multari *et al*. showed that no direct relationship exists between the number of strips used for sampling the skin on the forearm of a volunteer and the number of peptides identified.^20^ The authors indicate that the problem stems from overcrowding in the tubes that occurred during the extraction process. For avoiding this saturation during extraction we put each strip in a separate Eppendorf™ and extracted the bone powder and proteins from each skin tape-disc before bringing them together for digestion. The amount of peptides for the eraser method, acid fraction were similar to using 1 dermatological skin tape-discs. The amount of peptides was the lowest for the forced bag and ammonium bicarbonate etching methods. The mopping with swabs provided an amount equivalent to 5 dermatological skin tape-discs. Bone powder fraction of 1 milligram of bone yielded a greater amount of peptide. As stripping by 5 dermatological skin tape-discs method gave a better result than 1 dermatological skin tape-discs without modifying the appearance of the bone it was kept for further analyses.

The peptides obtained by digesting with trypsin the bone powder obtained by stripping with 5 skin tape-discs, the 4 low destructive samplings and the conventional destructive methods of 4 modern and 3 Neolithic bones were analyzed by MALDI FTICR. For each method, ZooMS markers peptide were identified and allowed to identify the taxonomic genus. The number of peptide markers for each sample and method is given in Supplementary Information 1, Figure S7. As expected, marker peptides were more easily detected in modern than Neolithic samples and their number was higher. The triboelectric charge methods (Eraser and forced bag) produced low quality spectra on archaeological samples but allowed taxa to be identified. Fewer marker peptides were identified compared to other methods. Several studies have shown that these methods can be used on ancient objects such as the St. Lawrence Iroquoian bone points (middle of the 14th to the late 16th centuries AD) analyzed by McGrath et al. or the Neanderthal bone artifacts analyzed by Martisius et al.^12, 16^ Coutu et al noted that the surface of the bones can influence the results. If the bones are smooth, the triboelectric charge methods have difficulty recovering bone particles, resulting in a low quantity of collagen and spectra with few marker peaks.^41^ However if the sample is crumbled, the forced bag method is most suitable.^19^ The type of plastic used have an impact on the results as the triboelectric charge density varied by more than three order of magnitudes from polytetrafluoroethylene (PTFE) or polyethylene(PE) to polyethyleneterephthalate(PET).^42^

For the swab mopping method, the number of marker peptides identified per sample was between 8 and 12. This method was never used to collect material but rather to clean objects. Surface analysis by microscopy showed a deposit of material on the surface of the bone.

The ammonium bicarbonate buffer etching method produced good results even though only 6 marker peptides were identified in the Trem2 sample. For the other samples, the number of peptides was 12 or 11. This method is less efficient when the bones are highly mineralized, which reduces extraction efficiency. Wang Naihui et al. showed that the addition of a demineralization step increased the number of identifiable samples.^43^

The results of the conventional destructive method for the acid fraction and bone powder fractions allowed better identification and detection of peptides markers than ammonium bicarbonate buffer etching method. In all the samples the 5 dermatological skin tape-disc stripping and the destructive sampling allowed identifying the most peptide markers. The 5 dermatological skin tape-disc stripping and conventional destructive method allowed the identification of the genus and causes little degradation of the samples.

The glue present on the patch was not found in the MALDI FTICR or the LC-MS/MS mass spectrometry analyses after digestion. The digestion with Amicon® filters should enable the glue polymer to be eliminated during the washing stages. So, we applied dermatological method on 19 paleontological and archaeological bones.

### Tape stripping method applied to palaeontological and archaeological bones: taxa identification

We applied our minimal invasive sampling approach based on dermatological skin tape-discs stripping on 19 bones from Iron Age, Neolithic and Upper Pleistocene bones dated from 130,000 to 100 BC. Firstly, after digestion, a MALDI FTICR analysis was achieved. Table S1 in Supplementary Information 4 contains the identified peptide biomarkers. The taxonomy of each bone has been identified by looking at the 12 ZooMS peptide biomarkers.^28^ MALDI FTICR MS analyses identified the taxonomy of all of the 19 samples. Only one sample Wa24 was identified as *Ursus* sp. while the identification made from osteomorphology by the paleontologists is *Cervus* sp. Although we used a MALDI FTICR MS, the more commonly used MALDI TOF MS can also be employed. It should be noted that the main difference between the two instruments is the 100 times higher resolution of the MALDI FTICR MS, which allows for the unambiguous detection of monoisotopic peak of native and deamidated peptides.

The same samples were also analyzed by LC-MS/MS to validate the MALDI FTICR MS identifications. Sample identification was performed using the NCBI mammalian and aves database which contains protein sequences from several sources, including annotated coding region translations in GenBank, RefSeq and TPA, as well as SwissProt records. The mammalian and aves databases were chosen because MALDI identifications only identified mammal and bird species in the same way as morphological identifications by archaeologists and palaeontologists. In this database, there are few protein sequences of extinct species such as cave bears and aurochs which complicates the identification of extinct species. To allow their identification, it is necessary to use the closest species at the phylogenetic level. For example, for the aurochs *Bos primigenius*, it is necessary to use the sequences of its closest ancestor which is *Bos taurus*.

Proteomic analysis of Iron age, Neolithic or Upper Pleistocene bones indicates that the major proteins identified are type I cytoskeleton keratin, type II cytoskeleton keratin, and type I collagen as in the control bone of modern *Bos taurus*. This information shows that the use of the patch allows the extraction of bone proteins despite the fossilization of the bone. The average number of identified peptides for the best identified proteins (collagen 1 alpha 1) on modern bones is 645, 458 on the Iron age, Neolithic samples and 190 on the Upper Pleistocene samples (Table 1). The preservation of the Iron age, Neolithic samples in the Tremblay-en-France and Bouchain, Pertus sites is exceptional, which explains the number of significant peptides identified in the samples. Sequence coverage varies between 59, 50 and 40% for modern, Iron age, Neolithic and Upper Pleistocene samples respectively based on the best identified proteins. The number of peptides and sequence coverage are lower in older bones due to collagen degradation over time. Supplementary Information 2 shows the identification of all proteins in the analyzed samples. The number of peptides for modern *Struthio camelus* samples is lower than others modern samples due to the processing of the bone. Indeed, this bone has been boiled to remove the flesh elements then cleaned with hydrogen peroxide. Despite this treatment, the analysis with the dermatological skin tape-discs and the preparation allows the identification of many peptides. This shows that the sampling method and proteomic analysis could be used to study bones contained in museums. This method would make it possible to obtain the global proteome and MALDI spectra of reference species. With these reference spectra, new marker peptides could be discovered even if the type I COL sequences are not in the databases. In this way, it will be possible to increase species identifications on archaeological sites.

**Table 1.** LC-MS/MS results for modern, Holocene and Upper Pleistocene bones sampled with dermatological skin tape-discs stripping. The table contains name of sample, taxonomic identification by archaeozoologists and paleontologists, the best identified proteins COL1A1, the species from NCBI database, the accession number, the Mascot score, the number of peptides and the coverage for the best identified protein.

| Name | ID<br>archaeologist | Protein | Species form<br>NCBI<br>database | Accession number | Mascot<br>score | Peptides | Coverage<br>(%) |
| --- | --- | --- | --- | --- | --- | --- | --- |
| Mod 1 | *Bos taurus* | Collagen<br>alpha-1(I)<br>chain | *Bos taurus* | AAI05185.1 | 9655 | 624 | 66 |
| Mod 2 | *Struthio*<br>*camelus* | Collagen<br>alpha-1(I)<br>chain | *Struthio*<br>*camelus* | XP_009685373.1 | 5192 | 254 | 56 |
| Mod 3 | *Capra ibex* | Collagen<br>alpha-1(I)<br>chain | *Capra hircus* | XP_017920382.1 | 11187 | 646 | 62 |
| Mod 4 | *Cervus*<br>*elaphus* | Collagen<br>alpha-1(I)<br>chain | *Cervus*<br>*canadensis* | XP_043327093.1 | 22864 | 1044 | 59 |
| PE-F01 | *Bos taurus* | Collagen<br>alpha-1(I)<br>chain | *Bos taurus* | AAI05185.1 | 12118 | 831 | 60 |
| PE-F21 | *Capra hircus* | Collagen<br>alpha-1(I)<br>chain | *Capra hircus* | XP_005678993.1 | 6701 | 396 | 58 |
| PE-F46 | *Sus* sp. | Collagen<br>alpha-1(I)<br>chain | *Sus scrofa* | XP_020922812.1 | 6613 | 488 | 52 |
| PE-F55 | *Ovis aries* | Hypothetical<br>protein<br>JEQ12_002924 | *Ovis aries* | KAG5203341.1 | 7575 | 512 | 49 |
| PE-F72 | *Cervus*<br>*elaphus* | Collagen<br>alpha-1(I)<br>chain | *Cervus*<br>*canadensis* | XP_043327093.1 | 5175 | 365 | 45 |
| Bou1 | *Sus scrofa* | Collagen<br>alpha-1(I)<br>chain | *Sus scrofa* | XP_020922812.1 | 6272 | 422 | 49 |
| Bou2 | *Bos taurus* | Collagen<br>alpha-1(I)<br>chain | *Bos taurus* | AAI05185.1 | 6637 | 399 | 57 |
| Bou3 | *Cervus*<br>*elaphus* | Collagen<br>alpha-1(I)<br>chain | *Cervus*<br>*canadensis* | XP_043327093.1 | 6456 | 359 | 63 |
| Bou4 | *Castor fiber* | Collagen<br>alpha-1(I)<br>chain | *Castor*<br>*canadensis* | JAV39636.1 | 4707 | 350 | 51 |
| Bou5 | *Capreolus*<br>*capreolus* | Hypothetical<br>protein<br>FD754_00932<br>5 | *Muntiacus*<br>*muntjac* | KAB0365169.1 | 8645 | 567 | 53 |
| Bou6 | *Sus scrofa* | Collagen<br>alpha-1(I)<br>chain | *Sus scrofa*<br>*domesticus* | BAX02568.1 | 5357 | 350 | 62 |
| Bou7 | *Bos taurus* | Collagen<br>alpha-1(I)<br>chain | *Bos taurus* | AAI49096.1 | 3639 | 250 | 44 |
| Trem 1 | *Equus sp.* | Collagen<br>alpha-1(I)<br>chain | *Equus asinus* | ACM24774.1 | 6219 | 400 | 49 |
| Trem 2 | *Bos taurus* | Collagen<br>alpha-1(I)<br>chain | *Bos taurus* | AAI05185.1 | 7388 | 410 | 62 |
| Lov11 | *Ursus*<br>*spelaeus* | Collagen<br>alpha-1(I)<br>chain | *Ursus*<br>*maritimus* | XP_040496745.1 | 4157 | 293 | 47 |
| Wa21 | *Equus sp.* | Collagen<br>alpha-1(I)<br>chain | *Equus asinus* | ACM24774.1 | 2349 | 178 | 45 |
| Wa22 | *Bos*<br>*primigenius* | Collagen<br>alpha-1(I)<br>chain | *Bos taurus* | AAI49096.1 | 1654 | 130 | 33 |
| Wa23 | *Cervus*<br>*elaphus* | Collagen<br>alpha-1(I)<br>chain | *Cervus*<br>*canadensis* | XP_043327093.1 | 1348 | 81 | 35 |
| Wa24 | *Cervus*<br>*elaphus* | Collagen<br>alpha-1(I)<br>chain | *Ursus*<br>*maritimus* | XP_040496745.1 | 4163 | 274 | 38 |

The *N-* and *C-*terminal parts are not identified because they are removed during collagen maturation.^44^ High number of the identified peptides correspond to the type I collagen sequence. Proteomic identification correlates with taxonomic identification by archaeologists and paleontologists using comparative anatomy. Analyses are 100% consistent with the family level and 95% consistent with the genus level of taxonomic identifications (Table 1). The LC-MS/MS analysis showed that the Wa24 sample corresponded to an Ursidae, as did the MALDI FTICR. This difference between the identification by mass spectrometry and that achieved by anatomy is due to the size of the bone, the preservation of the bone which did not allow for formal identification of the bone. Unique peptides identified by LC-MS/MS on collagen 1 alpha 1, 1 alpha 2, and 1 alpha 3 (major bone proteins), with Mascot software were selected and further analyzed with BlastP to validate family, genus and species. Table 2 lists the specific peptides for the species studied and the Supplementary Information 3 gathers the MS/MS of the unique peptides (Supplementary Information 3 Figure S5 to S16). This analysis was facilitated by the preliminary MALDI analysis that allowed targeting of the genus. The LC-MS/MS results show that dermatological skin tape-discs stripping allows identifying taxonomy of the bones using the collagen, but identification of the taxonomy is influenced by the collagen (I) reference database, and some taxa may be poorly resolved due to a lack of available sequences. Currently, the NCBI database contains 271,985,047 million sequences for 138,491 organisms but for example the COL1A1 and COL1A2 collagens of species of archaeological interest *Castor fiber* and *Capreolus capreolus* are not present. A second problem is the similarity of type I collagen sequences which does not allow discrimination between species as for *Ursus arctos horibilis, Ursus maritimus, Ursus americanus*. This indicates that discrimination between species should be done using other proteins present in the bones or teeth.^45^

**Table 2.**
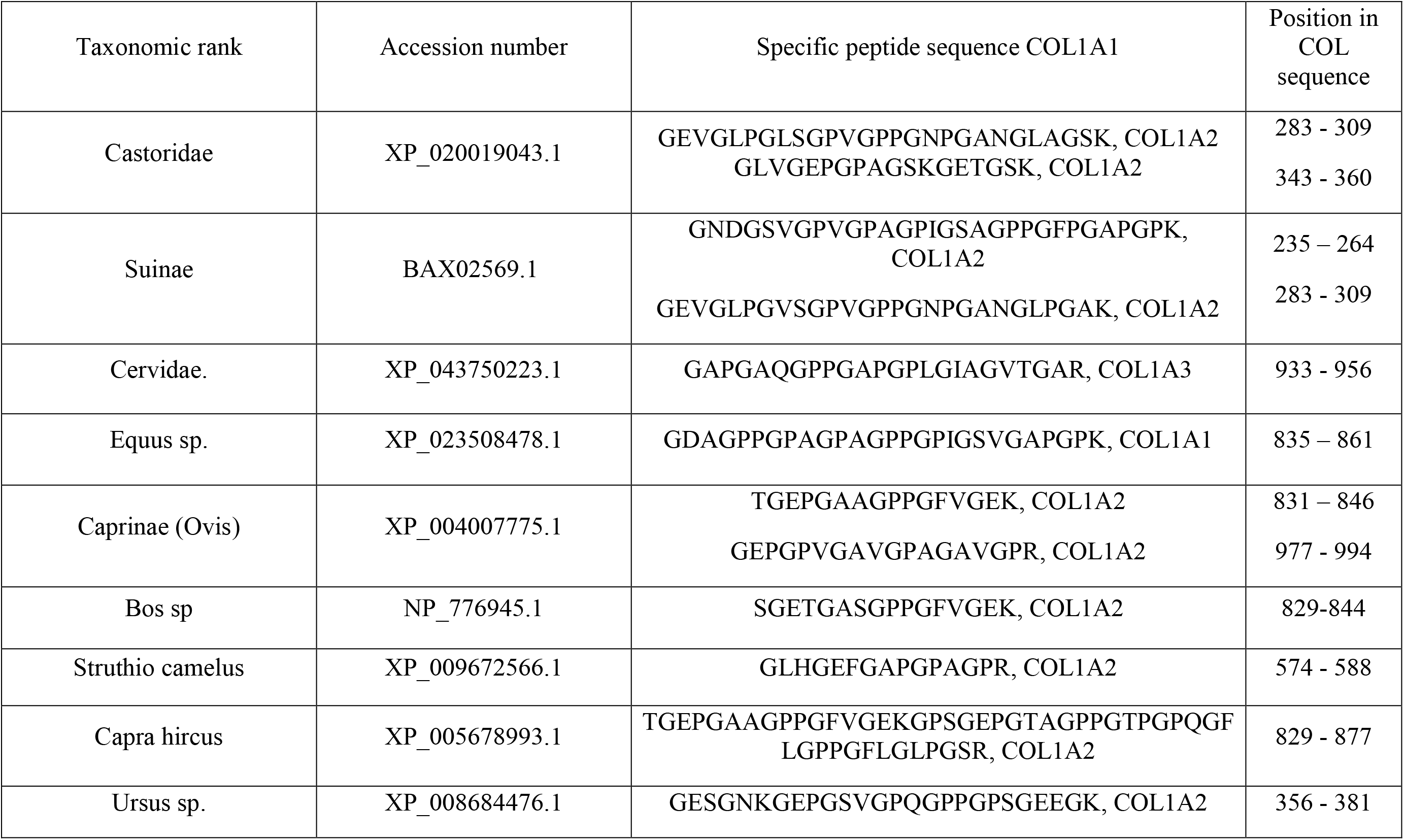
List of specific peptides of genus taxa for archaeological and paleontological sample identified by LC-MS/MS from bones sampled with dermatological skin tape-discs stripping and validated by BlastP with 100% identity and 100% cover query. The table includes the taxonomic rank, the collagen accession number, the peptide sequence and the position of the peptide on the sequence.

The database mining of the LC-MS/MS proteomics data led also to the identification of non-collagenous proteins (NCP) such as biglycan, chondroadherin, osteomodulin, calmodulin.^46^ This type of protein has been identified in bones from different periods and for different species such as humans, mammoths, moas, cattle, horses, turkeys, rabbits, squirrels, extinct rhinoceros. ^3, 28, 45–53^ Because many of the NCPs and some collagens have higher mutation rates than COL type 1, they are better targets for phylogenetic reconstructions, especially between closely related species.^8, 28^ The number of NCP proteins identified is higher in the Holocene samples than in the Upper Pleistocene. Although the stripping with dermatological skin tape-discs is a low-invasive method, it does identify minor proteins such as NCPs. The decrease in the number of these proteins is correlated with the smaller number of peptides identified in the Upper Pleistocene samples due to faster degradation of these proteins compared to collagen.^52^Both biglycan and lumican which exhibits collagen-binding properties were identified in modern, Iron Age and Neolithic samples but not in Pleistocene sample. This indicates that these proteins have been degraded in Pleistocene samples. Studies show that they can be found in samples as old as 650 Ka but using 50 mg of samples that is 50 times more at least than in this study.^51–52^

We detected also in the LC-MS/MS proteomics data a variety of *in vivo* and diagenetic post-translational modifications (PTMs) in the samples. Analysis of post-translational modifications such as glutamine deamidation provides information on the conservation status of bone. The degree of glutamine deamidation, has been previously reported to hold promising as an indicator of degradation in ancient materials.^54–57^ Older or degradated samples have a higher level of deamidation.^54–61^

Deamidation is used in many publications to validate that samples and proteins are ancient. Comparing deamidation data between publications is complex because (i) preparation methods are different and can influence the result. (ii) the environment can have an impact on the deamidation value even if the data from this publication does not show it. (iii) the calculation methods and software are different and do not give the same type of values.^36, 56–57, 59, 61–62^ The rate of deamidation between publications for bones from the same period may be different because the procedures for demineralization of bones, extraction of proteins are variable and may induce deamidation.^54–55, 59^ With regard to this last point, it is possible to obtain the percentage of glutamine deamidation from the peptide P1, the percentage of glutamine and asparagine deamidation of the peptides identified on type I collagen and other proteins. In 2012, Van Doorm *et al* showed that the deamidation rate of glutamine position on the collagen 1 alpha 1 are not identical.^55^

Figure 3 shows the percentages of deamidation in the samples. The percentage of deamidation on the glutamine based on the peptide P1^m^ for mammals and P1^b^ for birds increases significantly (Student’s *t test*, p<0.001) between modern vs Neolithic or Neolithic vs Pleistocene for the MALDI FTICR and LC-MS/MS analyses. Figure 3, the percentage of deamidation on the glutamine follows the same trend for both COL1A1 and COL1A2 based on Maxquant and the deamidation software package. A significant increase is observed between modern and pleistocene with this calculation method (Student’s *t test*, p<0.001, Figure 3). The percentage of deamidation on the glutamine based on the peptide P1 for both methods are correlated (*R*² = 0.91, Supplementary data S1, Figure S8). Although the two analytical methods MALDI FTICR and nanoLC-MS/MS Orbitrap have different ionization processes, the quantification data obtained on deamidation are similar. This deamidation on glutamine percentage for the peptide P1 determined by MALDI FTICR and LC-MS/MS correlates well with the global deamidation percentage for all glutamines calculated with Maxquant and the deamidation software package (*R*² = 0.94 and *R*² = 0.88, Supplementary data S1, Figure S9; Figure S10). This indicates that the calculation of the deamidation on glutamine using the peptide P1 or all potentially deamidated positions on COL1A1 or COL1A2 is correct. Even if some positions have a faster or slower deamidation rate, this has little impact on the result compared with the peptide P1 analysis alone.

**Figure 3.**
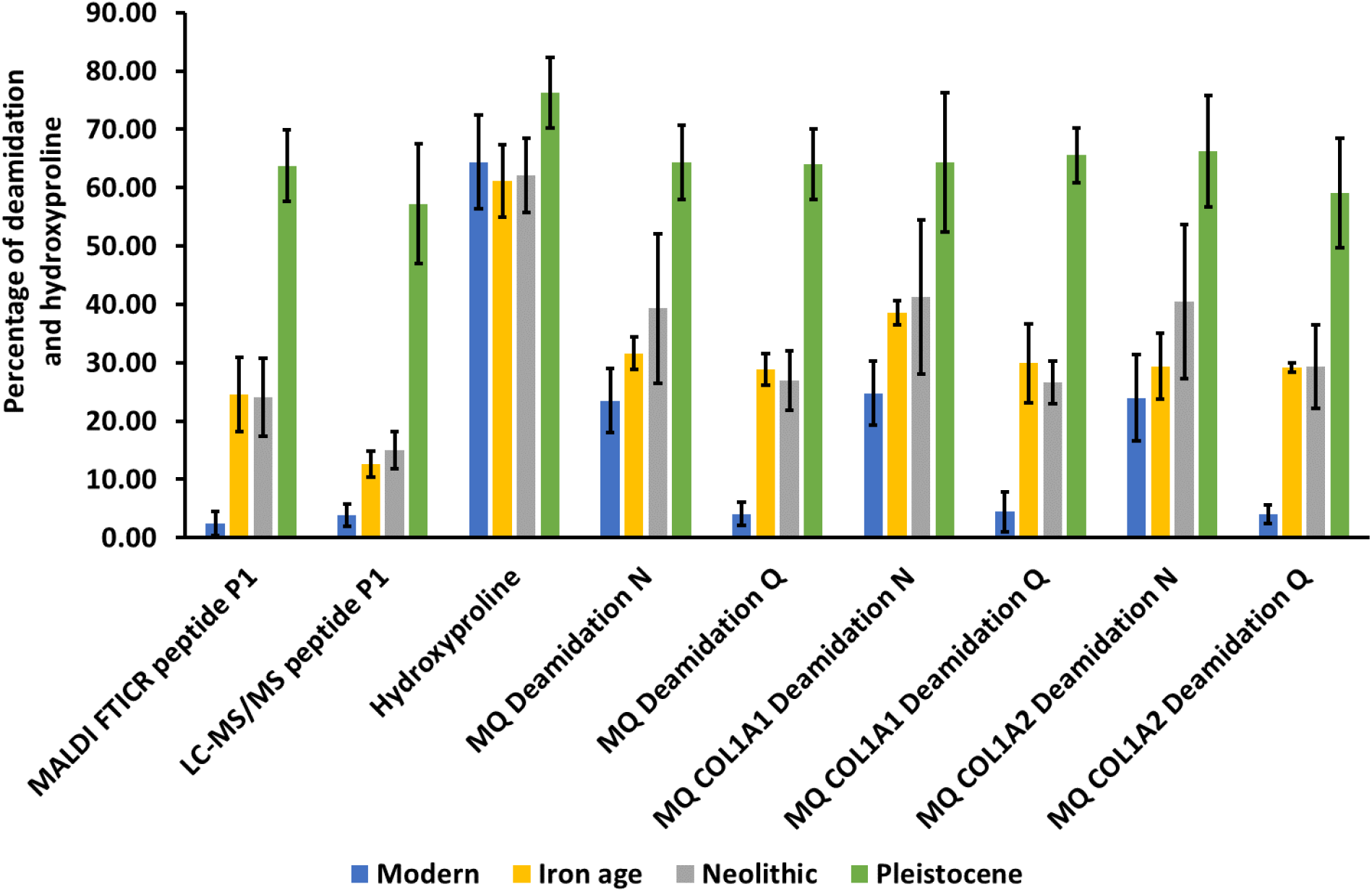
Percentage of deamidation and hydroxyproline for archeological samples by MALDI FTICR and LC-MS/MS. Bleu: modern sample, orange: Iron Age, grey: Neolithic and green Pleistocene. The deamidation were calculated with MALDI FTICR and LC-MS/MS based on the peptide P1^m^ and P1^b^, the hydroxylation value based on LC-MS/MS with PEAKS X+ PTM and deamidation N and Q were calculated with LC-MS/MS with Maxquant and deamidation package.

The percentage of the deamidation on the asparagine increase between modern and plesitocene and iron age, Neolithic and pleistocene (Student’s *t test*, p<0.001, Figure 3). There is no significant difference between modern and Iron Age or Neolithic sample for the deamidation on the asparagine. The percentage of deamidation on asparagine is sligthy higher than glutamine but without significant value. Pal Chowdhury *at al*, showed that the glutamine being much more stable towards deamidation than asparagine.^57^ The correlation of the global deamidation percentage of asparagine (based on Maxquant) with the deamidation on glutamine based on the peptide P1 obtained for MALDI and LC-MS/MS methods is slightly lower (*R*² = 0.70 and *R*² = 0.61, Supplementary data S1, Figure S11; S12) when compared to the correlation of glutamine deamidation (Supplementary data S1, Figure S9; Figure S10). Finally, the correlation between the global percentages of glutamine and asparagine deamidation determined by LC-MS/MS and calculated with Maxquant and the deamidation software package is a bit better (*R*² = 0.71, Supplementary data S1, Figure S13).

It is worth to notice that is no significant difference between the glutamine deamidation percentages between the bones from Temblay-en-France and Bouchain (both open air sites, Iron Age 2nd-1st century BC, Neolithic 3200-2900 BC) and the bones from Pertus II (cave site, Late Neolithic 3800 – 3300 BC). Comparing the deamidation rates between the three sites, it seems counter-intuitive that the rate for the Iron-Age samples is not the lowest. However, it has been described that storage conditions such as humidity and oxygen levels have an influence on the rate of deamidation.^56^ Ashley N. Coutu *et al* studied the domestication of sheep in southern Africa around 2000 BP through paleoproteomics.^41^ Using the deamidation measurement method with Maxquant and deamidation package, they obtained 70% of deamidation for asparagine and 40% for glutamine deamidation for ancient samples. The Bouchain and Pertus samples, which are closest in terms of dates, have slightly different values for glutamine deamidation and asparagine. The asparagine deamidation values are 29.8% +/− 6.1% and 52.5% +/− 4.5%, the glutamine deamidation values are 24.2% +/− 2.8% and 30.6% +/− 5.3% for Bouchain and Pertus site respectively. Concerning the modern sample, the deamidation for asparagine is 50% and 10% for glutamine deamidation for Ashley N. Coutu *et al*. The deamidation values for the modern sample in our study are slightly different (23.5% +/− 5.5% of deamidation for asparagine and 4.1% +/− 2.0% for glutamine deamidation). Comparing data is complicated by the fact that values can fluctuate due to sample preparation methods and environmental conditions.

The de novo analysis of the PTMs present in the samples using the PEAKS X+ software shows the presence of hydroxyproline. Hydroxyproline is naturally present in collagen to stabilize the structure.^63–64^ The average percentage of hydroxylation is 65%, with no significant difference between periods but a slight increase in Pleistocene sample (Figure 3). The percentage of hydroxylation is 12-13% in fresh collagen.^65^ There is an increase in hydroxyproline in archaeological bones, but this change is not correlated with sample age.

## CONCLUSION

Our work shows that dermatological skin tape-disc stripping, leaves very little trace on the bones, Proteomic method and high-resolution spectrometer allow identifying the proteins contained in bones giving access to the taxonomy identification of bones from today up to Pleistocene. The comparison of the different sampling methods shows that using dermatological skin tape-disc gives similar results to destructive acid demineralization, suggesting that dermatological skin tape-disc is a minimally invasive alternative to destructive sampling. The study of the deamidation shows that it is possible to differentiate modern, Neolithic and Plesitocene period from bones. The study of this modification would make it possible to identify bones from different periods on a site. It has also been shown that hydroxyproline is affected by bone ageing, but does not correlate with bone age. Elucidating the origin and significance of a bone fragment or artefact, while preserving its integrity, is a crucial issue for archaeologists, and the use of dermatological skin tape-disc is an alternative for these studies. Our study shows a potential application for museum analysis of reference and archaeological bones or artefact analysis. Our methods will allow to create a database of reference sample form museum for paleoproteomics community. This database will allow the identification of species for which no MALDI or LC-MS/MS data are available.

## ASSOCIATED CONTENT

**Supporting Information 1** contains the list of bone samples, the pictures of the bones before and after sampling, the comparison of sampling methods with peptide quantity and number of ZooMS markers, the correlation of the percentages of deamidation between MALDI FTICR MS and LC-MS/MS (file type, Adobe PDF).

**Supporting Information 2** contains the identification of proteins by LC-MS/MS (file type, Excel xlsx).

**Supporting Information 3** contains MS/MS spectra of peptides obtained by LC-MS/MS (file type, PDF).

**Supporting Information 4** contains the identification using ZooMS markers (file type, xlsx).

## AUTHOR INFORMATION

### Corresponding Author

Fabrice Bray – Univ. Lille, CNRS, USR 3290 – MSAP – Miniaturisation pour la Synthèse, l’Analyse et la Protéomique, F-59650 Lille, France; ORCID: https://orcid.org/0000-0002-4723-8206; Phone: +33 (0)3 20 33 71 12;

### Authors

Isabelle Fabrizi – Univ. Lille, CNRS, USR 3290 – MSAP – Miniaturisation pour la Synthèse, l’Analyse et la Protéomique, F-59650 Lille, France; ORCID: https://orcid.org/0000-0001-8934-4683; Phone: +33 (0)3 20 33 71 12;

Stéphanie Flament – Univ. Lille, CNRS, USR 3290 – MSAP – Miniaturisation pour la Synthèse, l’Analyse et la Protéomique, F-59650 Lille, France; ORCID: https://orcid.org/0000-0002-0135-675X; Phone: +33 (0)3 20 33 71 12;

Christian Rolando – Univ. Lille, CNRS, USR 3290 – MSAP – Miniaturisation pour la Synthèse, l’Analyse et la Protéomique, F-59000 Lille, France; Shrieking Sixties, F-59650 Villeneuve-d’Ascq, France. ORCID: https://orcid.org/0000-0002-3266-8860; Phone: + +33 (0)3 20 43 49 77;

Claire Delhon – Univ Côte d’Azur, CNRS, CEPAM – Culture Environnement Préhistoire Antiquité Moyen Age, F-06300 Nice, France; ORCID: https://orcid.org/0000-0002-7216-0148; Phone: + +33 (0)4 89 15 24 02;

Lionel Gourichon – Univ Côte d’Azur, CNRS, CEPAM – Culture Environnement Préhistoire Antiquité Moyen Age, F-06300 Nice, France; ORCID: https://orcid.org/0000-0002-5160-5902;

Manon Vuillien – Univ Côte d’Azur, CNRS, CEPAM – Culture Environnement Préhistoire Antiquité Moyen Age, F-06300 Nice, France; ORCID: https://orcid.org/0000-0001-7657-7613;

Tarek Oueslati – Univ. Lille, CNRS, UMR8164 – HALMA – Histoire Archéologie Littérature des Mondes Anciens, F-59650 Lille, France; ORCID: https://orcid.org/0000-0002-2886-085X; Phone: +33 (0)3 20 41 66 13;

Patrick Auguste – Univ. Lille, CNRS, UMR 8198 – EEP – Evolution, Ecology and Paleontology, F-59650, Lille, France ORCID: https://orcid.org/0000-0002-8302-6307;

### Author Contributions

I.F. methodology, writing-original draft. F.B. Conceptualization, methodology, writing-original draft, project administration, funding acquisition. S.F methodology. C. R: writing-review-editing, funding acquisition. C.D, L. G, M. V: review and access of the bones. T.O, P. A: review, access of the bones and funding acquisition. All authors have given approval to the final version of the manuscript

### Funding Sources

The authors acknowledge the IBiSA network for financial support of the UAR 3290 (MSAP) proteomics facility. The mass spectrometers were funded by University of Lille, CNRS, Région Hauts-de-France and the European Regional Development Fund. The authors deeply thank CNRS – Mission pour l’Interdisciplinarité for the funding of the Prot_HR_DAT project (2021-2022) and the AgroPastCN project (2020-2021). The authors are grateful to the Région Hauts-de-France for the funding of the ProtéOsHdF project (2021-2022) and the financial support from the IR INFRANALYTICS FR2054 for conducting the research is gratefully. The EU_FTICR _MS project Grant Agreement 731077, and the IPERION HS project Grant Agreement 871034) funded by EU Horizon 2020 Research and Innovation Program are profoundly thanked.

### Notes

Any additional relevant notes should be placed here.

## Supporting information

Supporting Information 1 contains the list of bone samples, the pictures of the bones before and after sampling, the comparison of sampling methods wi

Supporting Information 2 contains the identification of proteins by LC-MS/M

Supporting Information 3 contains MS/MS spectra of peptides obtained by LC-MS/MS

Supporting Information 4 contains the identification using ZooMS markers

## ACKNOWLEDGMENT

The authors thank warmly Cédric Lepère, (ÉVEHA –Études et valorisations archéologiques, Limoges, France and Univ. Côte d’Azur, CNRS, UMR 7264 – CEPAM – Cultures et Environnements Préhistoire, Antiquité, Moyen Âge, Nice, France) for allowing the authors to sample material from the cave Pertus II (Méailles, Alpes-de-Haute-Provence, France).

## Notes

### Competing Interest Statement

The authors have declared no competing interest.

