## Supporting Information 1 contains the list of bone samples, the pictures of the bones before and after sampling, the comparison of sampling methods wi for "Low-invasive sampling method for taxonomic for the identification of archaeological and paleontological bones by proteomics of their collagens"

**Table of content**

[**Supplementary Figure S4.** Optic microscopy (magnification ×4, Optika B-383PLi, Italy) sampling method on Bos taurus bone (Mod 1). A, C, E, G, I: before sampling; B, D, F, H, J, K, L: after sampling. A, B: Triboelectric rubbing in forced bag sampling; C, D: Eraser sampling by rubbing, E, F: Swab mopping sampling; G, H: Conventional destructive sampling, , I, J, K, L: 1 D-Squame® skin tape-disc method. J: 5 D-Squame® skin tape-disc; K: 10 D-Squame® skin tape-disc; L: 20 D-Squame® skin tape-disc. The black circle indicates a modification of the surface after sampling. The black bar represents 1 cm.	13](#_Toc145939526)

[**Supplementary Table S3.** Mass to charge ratio (m/z) of the native and deamidated universal peptides. The table contains the peptide sequence, the post-translational modifications, the formula, the [M+H]^+^ of peptides using for MALDI FTICR deamidation values and the [M+2H]^2+^ of peptides using for LC-MS/MS deamidation values.	17](#_Toc145939536)

**Sites information**

**Waziers “Le Bas-Terroir” (Nord, France)**

This site is located in France in the Nord department. Waziers “Le Bas-Terroir” site is dated to the end of the Chibanian (new name for the Middle Pleistocene) and the beginning of the Upper Pleistocene (130,000-120,000 BC).{Hérisson, 2022 #232} The remains are divided into 10 stratigraphic units: US2, US3, US4, US5a to US5e, US6 and US7. The meso and macrofauna discovered in the Waziers sequence were analysed exhaustively and a list of the taxa present was drawn up. The aurochs (*Bos primigenius*) is the best represented species, followed by the roe deer (*Capreolus capreolus*), the beaver (*Castor fiber*), the Achenheim horse (*Equus achenheimensis*) and the prairie rhinoceros (*Stephanorhinus hemitoechus*).{Auguste, 2022 #233}

**The Loverval Deposit (Hainault Province, Belgium)**

This site is located west of the municipality of Gerpinnes near Charleroi in Belgium. It is a karst complex located in the sub-basin of the Sambre. Situated in a Carboniferous limestone massif, its total underground extent is estimated to be about 120 meters. An age of 33,050 BC was first estimated by the speleologist team, following the size determination of the flint, attributed to the Upper Paleolithic, and the presence of Open Arctic taxa; but since sedimentological and stratigraphic information is lacking, the chronological attribution of this site is rather imprecise. Regardless, it is estimated that the age of this site is between the beginning of the MIS 2 and 4 stages, i.e. between c. 25,050 and 63,050 years BC.{Auguste, 2012 #173} The faunal remains, which are scanty but excellently preserved, belong mainly to carnivores, in particular the cave bear *Ursus spelaeus* and the cave hyena *Crocuta crocuta spelaea*. They are associated to a lesser extent with remains of wolf *Canis lupus*, cave lion *Panthera spelaea*, woolly mammoth *Mammuthus primigenius*, woolly rhinoceros *Coelodonta antiquitatis* and horse *Equus* sp.

**Bouchain (Nord, France)**

The site of Bouchain is the first recent Neolithic site (3200-2900 cal. BC) excavated in Northern France. The excavation campaigns have revealed an activity zone on the bank of a paleochannel of the Scheldt river. The fauna comprising 1892 bone remains buried in a waterlogged sediment. These archaeological bones are very well-preserved. Traces of cutting, breaking and cooking were observed on the surface of the bones. On the Bouchain site, wild mammals and birds are important elements of the subsistence strategy followed by poultry farming. This is a unique Final Neolithic or Middle Neolithic site in the regional Nord of France context.{Oueslati, 2022 #174}

**Grotte de Pertus II (Alpes-de-Haute-Provence, France)**

The Pertus II cave is located in France in the Alpes-de-Haute-Provence at 1000 m a.s.l. between alpine and mediterranean ecologic influences. It was occupied mainly during the Middle and Late Neolithic between 4600-2500 BC, but the major occupation sequence corresponds to the Chassean D & E (3800–3300 BC). This cave presents well-preserved sheepfold levels, alternating with pottery workshop and dwelling.{Battentier, 2016 #176} The study of faunal remains is still ongoing, but the preliminary results indicate a predominance of livestock, particularly sheep and goat, and a high diversity of wild taxa such as wild pig, red deer, roe deer, chamois and hare.{Vuillien, 2020 #227}

**Tremblay-en-France (Seine-Saint-Denis, France)**

The excavation of an Iron Age rural settlement at Tremblay-en-France (Seine-Saint-Denis, France) dating back to the 2^nd^-1^st^ century BC provides evidence of a modest agricultural establishment. The settlement was primarily focused on the production of cereals and the breeding of cattle and sheep. Initially, the site was located within a wooded environment, but over time, it became increasingly deforested.{Broutin, 2021 #234}

**Supplementary Table S1.** Bone Specimens used in the study. The table contains sample name, location, geological sequence, morphology and taxonomy

| **Sample name** | **Site** | **Location** | **Country** | **Period** | **Sequence** | **Datation (BC)** | **Description** | **Taxonomy** |
| --- | --- | --- | --- | --- | --- | --- | --- | --- |
| Mod 1 | Lille | Hauts-de-France | France | Modern |  |  | Mandible | *Bos taurus* |
| Mod 2 | Lille | Hauts-de-France | France | Modern |  |  | Femur | *Struthio camelus* |
| Mod 3 | Lille | Hauts-de-France | France | Modern |  |  | Pelvis | *Capra ibex* |
| Mod 4 | Lille | Hauts-de-France | France | Modern |  |  | Mandible | *Cervus elaphus* |
| Bou 1 | Bouchain | Hauts-de-France | France | Neolithic | 19 L10 US12 | 3200-2900 | Mandible | *Sus domesticus* |
| Bou 2 | Bouchain | Hauts-de-France | France | Neolithic | 19 L10 US12 | 3200-2900 | Phalanx | *Bos taurus* |
| Bou 3 | Bouchain | Hauts-de-France | France | Neolithic | 19 M17 US12 | 3200-2900 | Radius | *Cervus elaphus* |
| Bou 4 | Bouchain | Hauts-de-France | France | Neolithic | 19 D3 US11 | 3200-2900 | Mandible | *Castor canadensis* |
| Bou 5 | Bouchain | Hauts-de-France | France | Neolithic | 19 D3 US11 | 3200-2900 | Metatarsal | *Capreolus capreolus* |
| Bou 6 | Bouchain | Hauts-de-France | France | Neolithic | 19 N16 US12 | 3200-2900 | Tibia | *Sus scrofa* |
| Bou 7 | Bouchain | Hauts-de-France | France | Neolithic | 18 Q26 US12 | 3200-2900 | Femur | *Bos primigenius* |
| Trem 1 | Tremblay-en-France | Seine-Saint-Denis | France | Iron Age | US 2982 | 260 - 150 | Scapula | *Equus sp.* |
| Trem 2 | Tremblay-en-France | Seine-Saint-Denis | France | Iron Age | US 2982 | 260 - 150 | Tibia | *Bos taurus* |
| PE-F01 | Pertus II | Alpes-de-Haute-Provence | France | Chasséen | 2016-2762-2974 | 3800-3300 | Radius | *Bos taurus* |
| PE-F21 | Pertus II | Alpes-de-Haute-Provence | France | Chasséen | 2017-5100PREL-5250 | 3800-3300 | Ulna | *Capra hircus* |
| PE-F46 | Pertus II | Alpes-de-Haute-Provence | France | Chasséen | 2017-4875-5091 | 3800-3300 | Mandible | *Sus sp.* |
| PE-F55 | Pertus II | Alpes-de-Haute-Provence | France | Chasséen | 2016-2762-2939 | 3800-3300 | Mandible | *Ovis aries* |
| PE-F72 | Pertus II | Alpes-de-Haute-Provence | France | Chasséen | 2018-5210-5240b | 3800-3300 | Occipital | *Cervus elaphus* |
| Lov11 | Loverval | Wallonia | Belgium |  |  | 63 ka | Mandible | *Ursus spelaeus* |
| Wa21 | Waziers | Hauts-de-France | France | Upper pleistocene | WBT 17, US II | 132 – 123 ka | Humerus | *Equus achenheimensis* |
| Wa22 | Waziers | Hauts-de-France | France | Upper pleistocene | WB 17, US IV, 615 | 132 – 123 ka | Humerus | *Bos primigenius* |
| Wa23 | Waziers | Hauts-de-France | France | Upper pleistocene | WB17 | 132 – 123 ka | Metapodial | *Cervus elaphus* |
| Wa24 | Waziers | Hauts-de-France | France | Upper pleistocene | WB 17 | 132 – 123 ka | Long bone | *Cervus elaphus ?* |

| 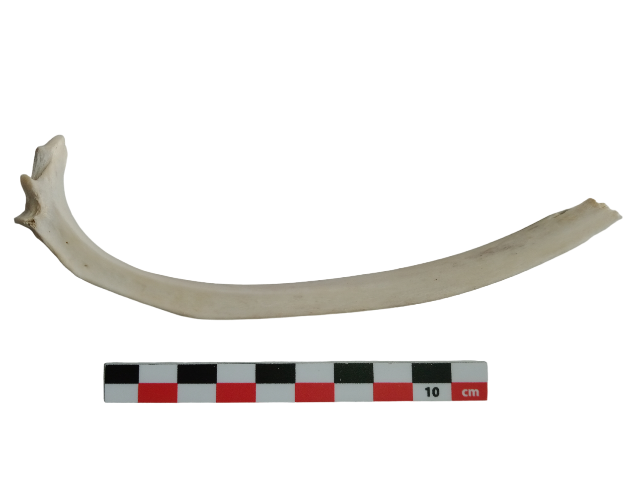A | 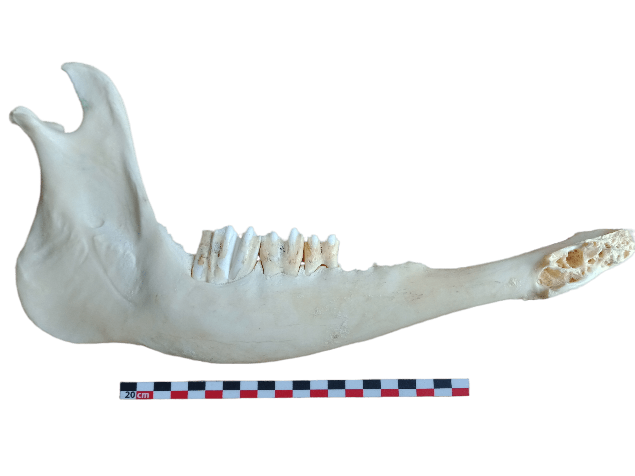 B |
| --- | --- |
| 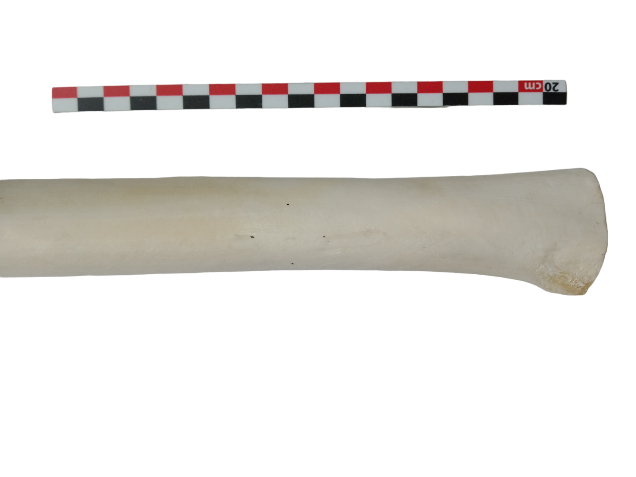C | 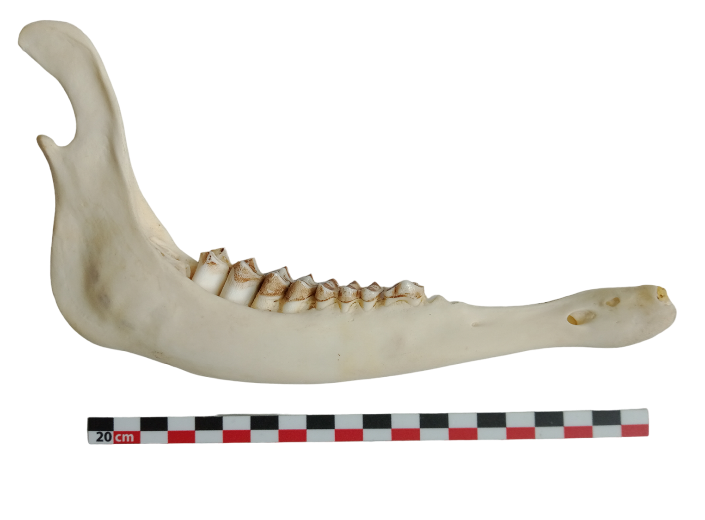D |

**Supplementary Figure S1*.*** Picture of modern bones. A- *Capra ibex* rib, B- *Bos taurus* mandible, C- *Struthio camelus* femur, D- *Cervus elaphus* mandible. One square of the scale bar represents 1 cm.

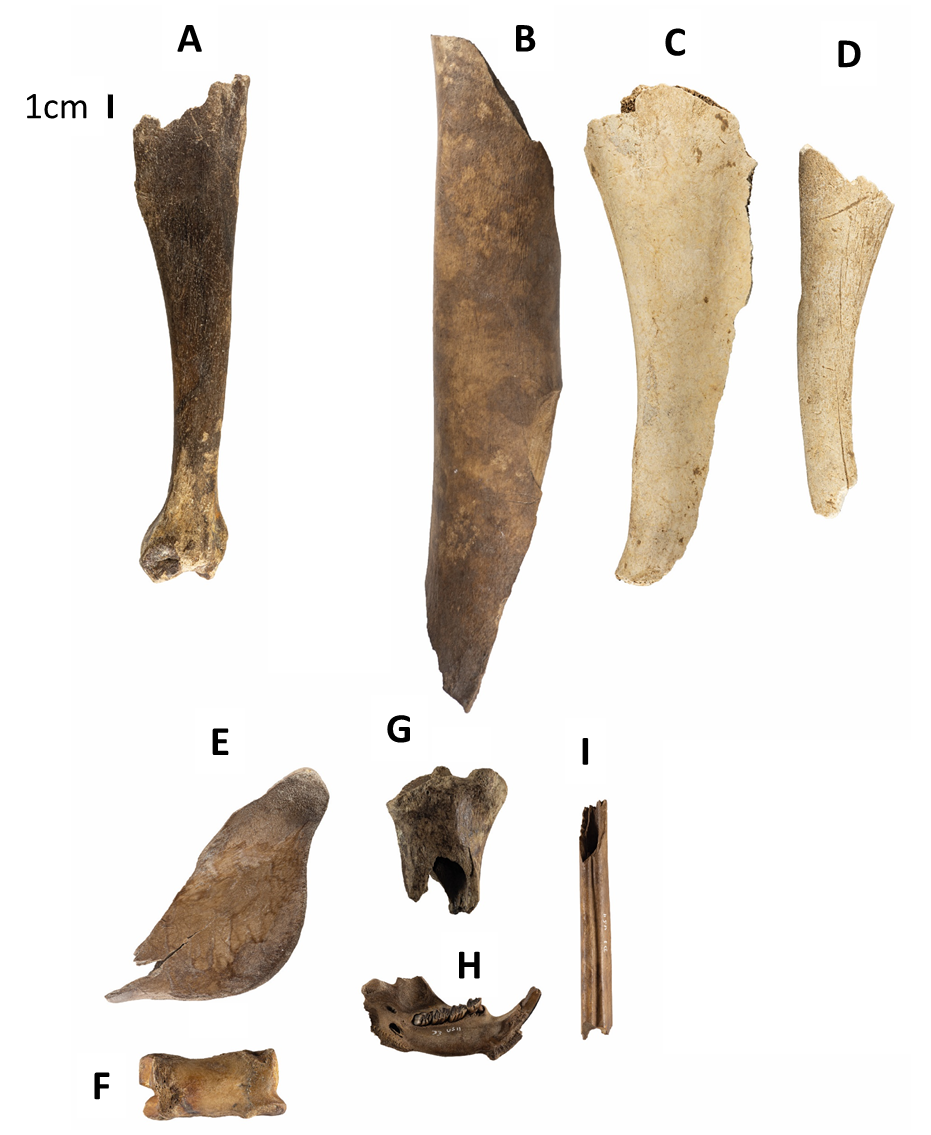

**Supplementary Figure S2*.*** Picture of archaeological bones from Bouchain, Tremblay-en-France. A: wild boar tibia (Bouchain); B: aurochs femur (Bouchain); C: horse scapula (Tremblay-en-France); D: cattle tibia (Tremblay-en-France); E: pig mandible (Bouchain); F: cattle first phalanx (Bouchain); G: red deer radius (Bouchain); H: beaver mandible (Bouchain); I: roe deer metatarsal (Bouchain). The scale bar represents 1 cm.

| 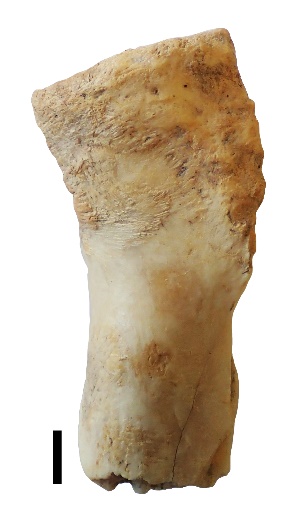A | 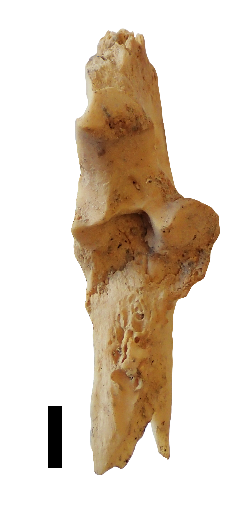B |
| --- | --- |
| 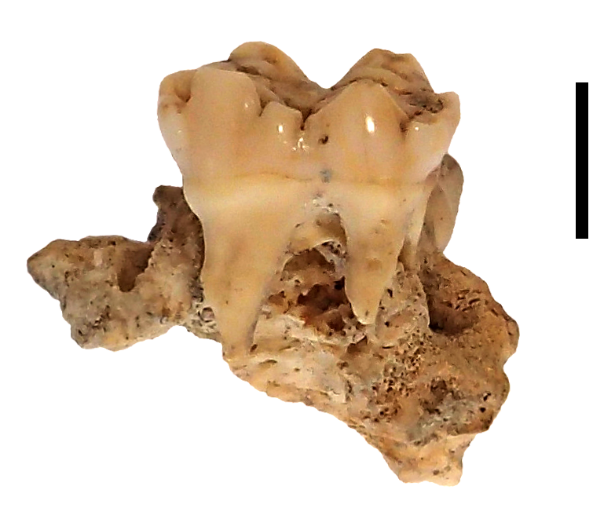C | 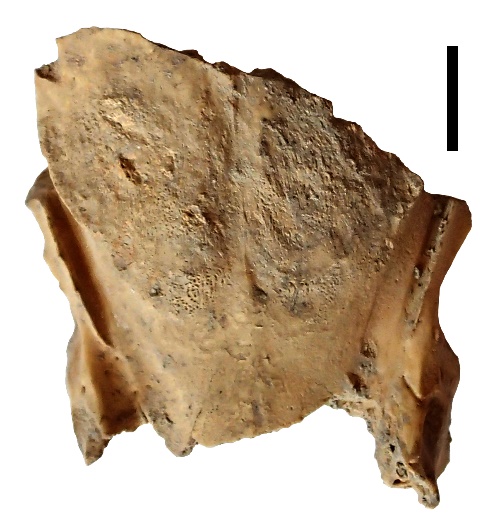D |
| 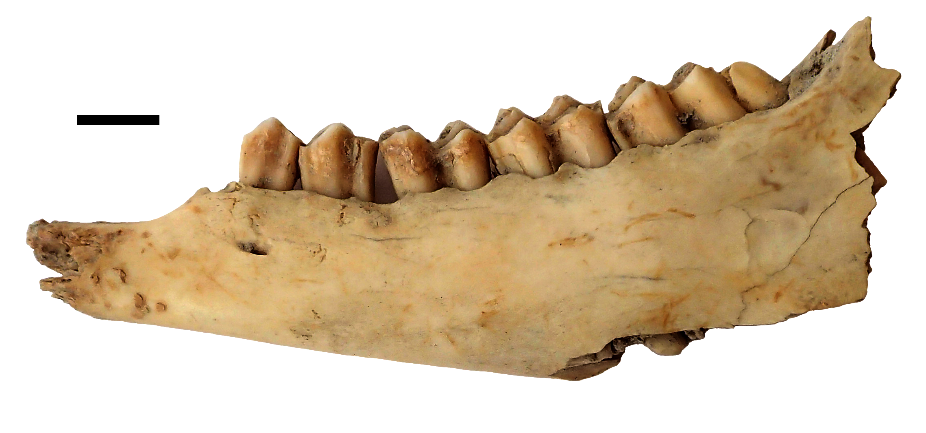E | |

**Supplementary Figure S3*.*** Picture of archaeological bones from Pertus. A- *Bos taurus* radius, B – *Capra hircus* ulna, C – *Sus sp* madibule, D – *Cervus elaphus* occipital, E – *Ovis sp* mandible. The black bar represents 1 cm.

| 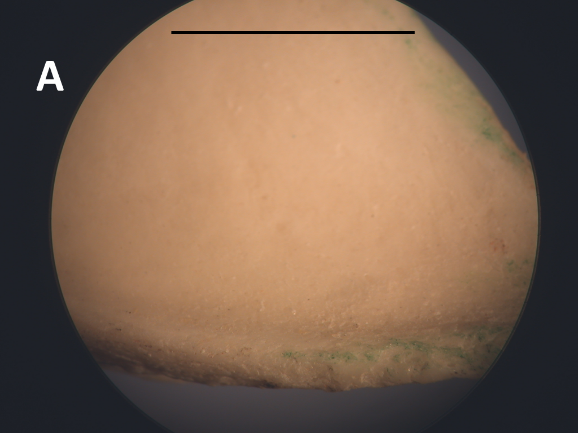 | 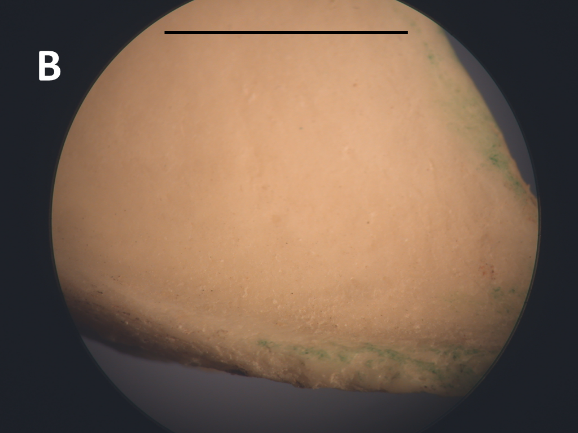 |
| --- | --- |
| 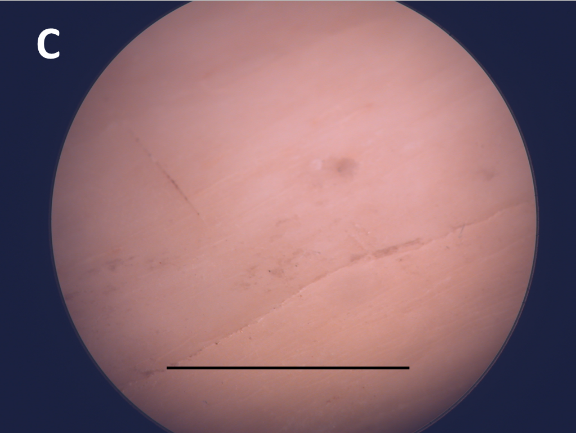 | 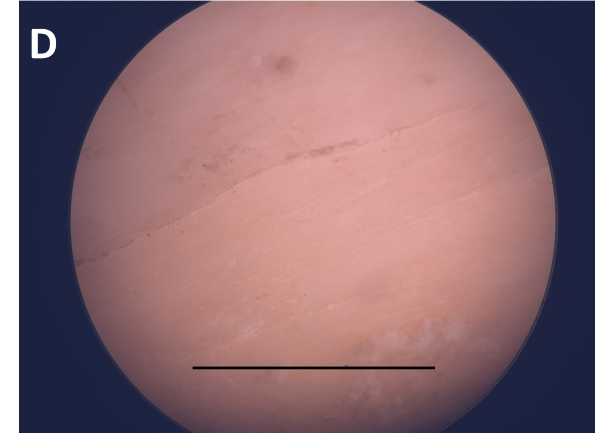 |
| 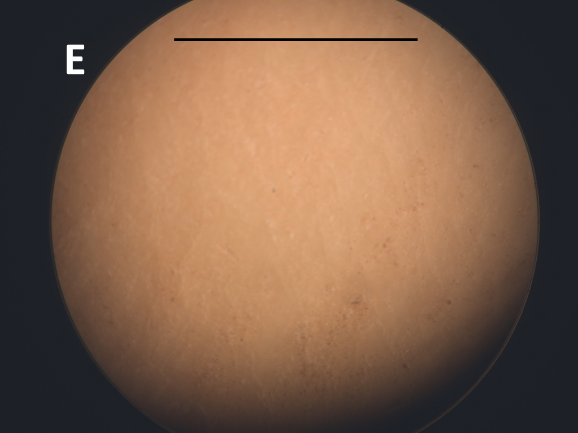 | 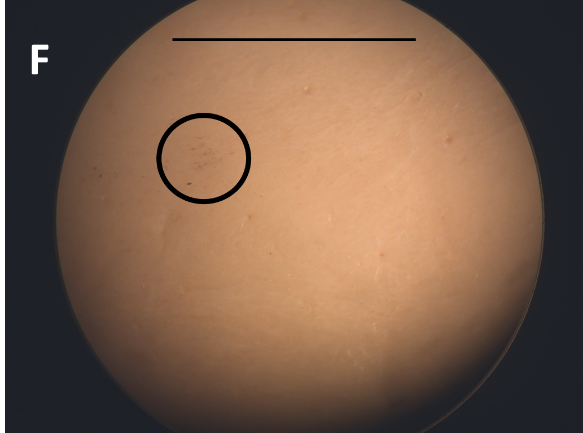 |
| 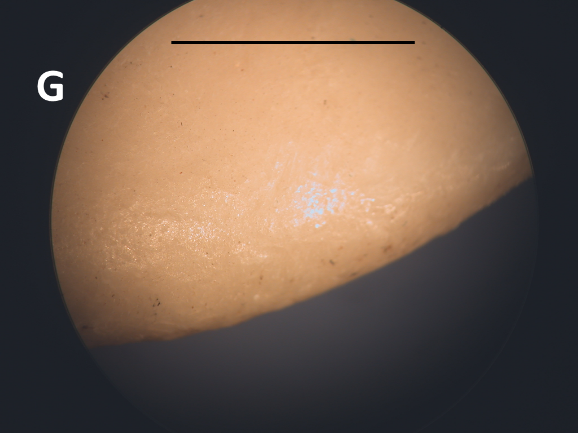 | 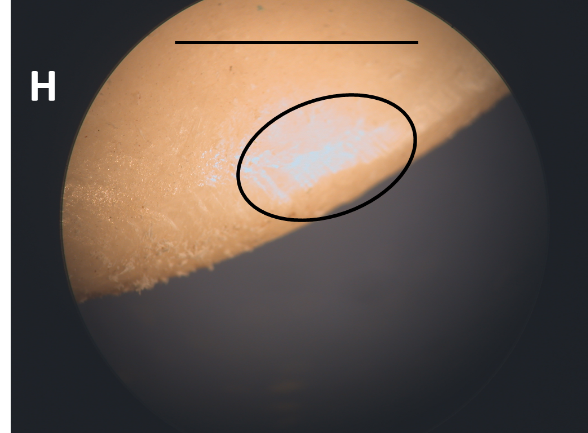 |
| 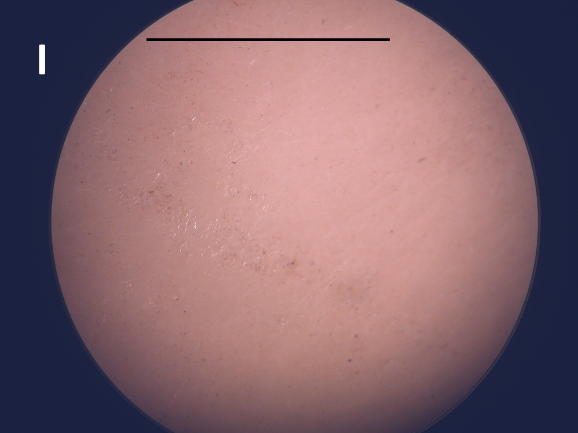 | 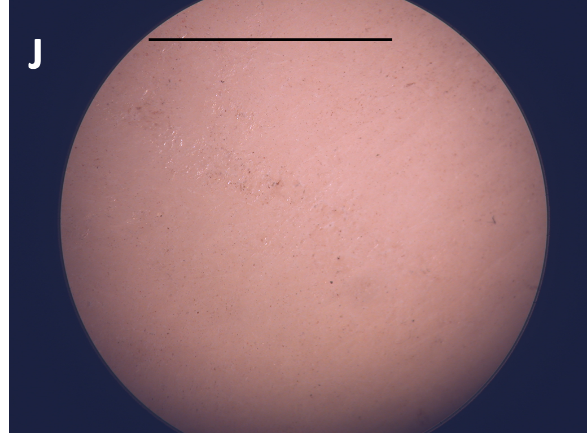 |
| 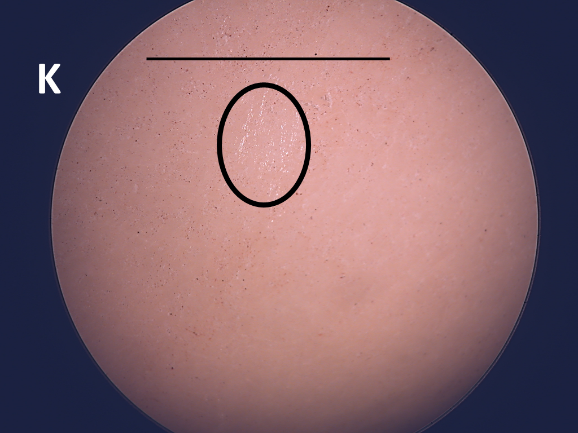 | 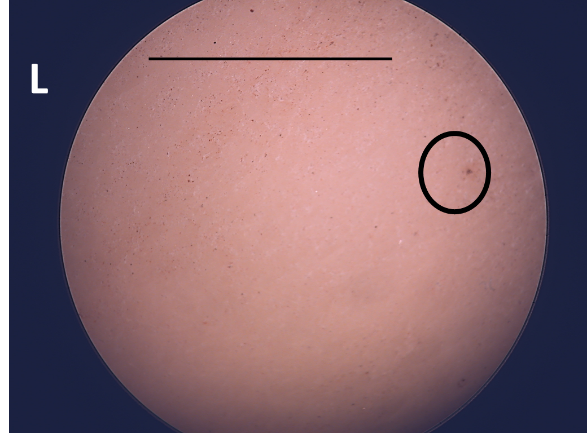 |

**Supplementary Figure S4.** Optic microscopy (magnification ×4, Optika B-383PLi, Italy) sampling method on *Bos taurus* bone (Mod 1). A, C, E, G, I: before sampling; B, D, F, H, J, K, L: after sampling. A, B: Triboelectric rubbing in forced bag sampling; C, D: Eraser sampling by rubbing, E, F: Swab mopping sampling; G, H: Conventional destructive sampling, I, J, K, L: 1 D-Squame® skin tape-disc method. J: 5 D-Squame® skin tape-disc; K: 10 D-Squame® skin tape-disc; L: 20 D-Squame® skin tape-disc. The black circle indicates a modification of the surface after sampling. The black bar represents 1 cm.

| 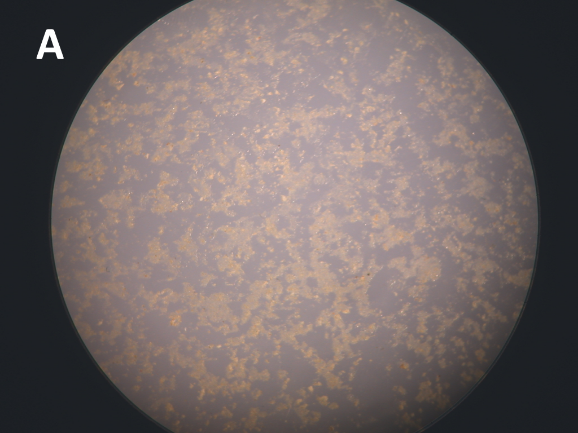 | 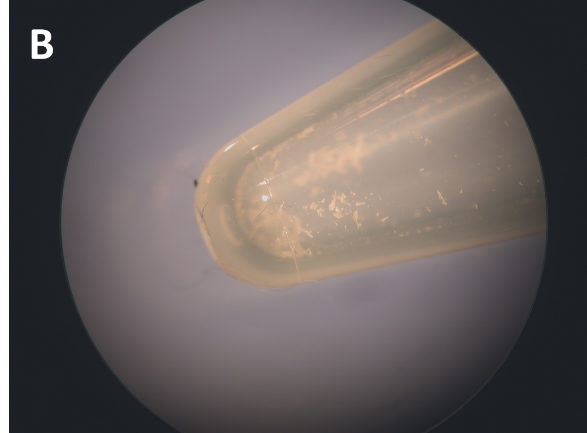 |
| --- | --- |

**Supplementary Figure S5.** Optic microscopy (magnification ×4, Optika B-383PLi, Italy) of A) the D-Squame® skin tape-disc and B) powder obtained after scraping. A thin layer of bone is visible.

| 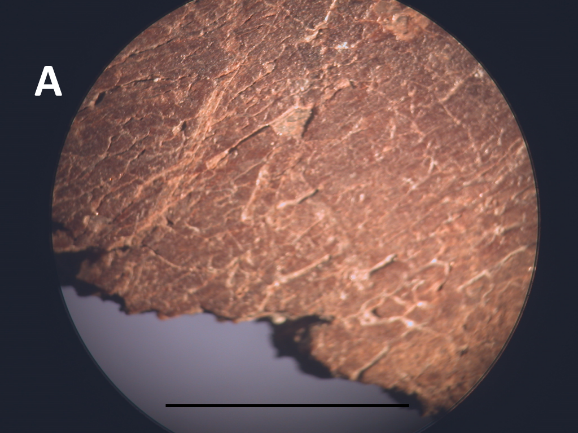 | 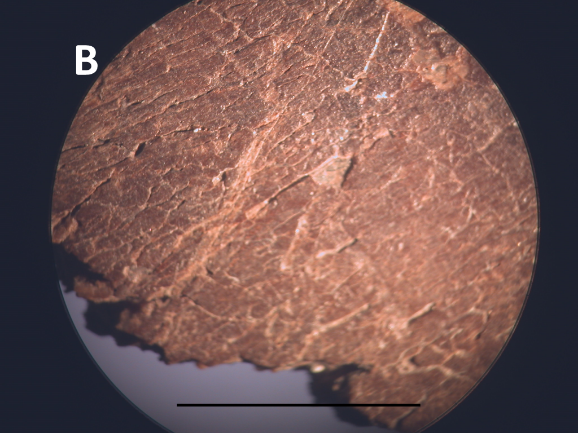 |
| --- | --- |
| 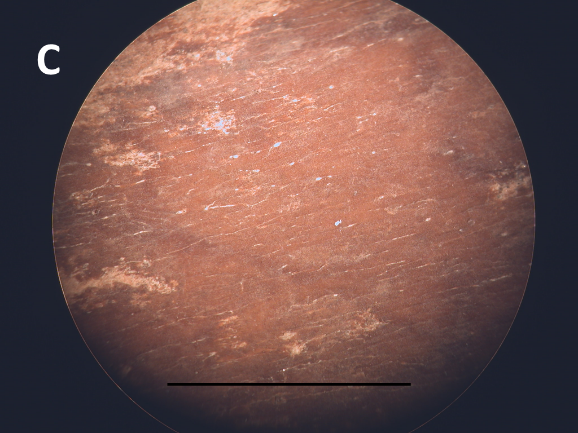 | 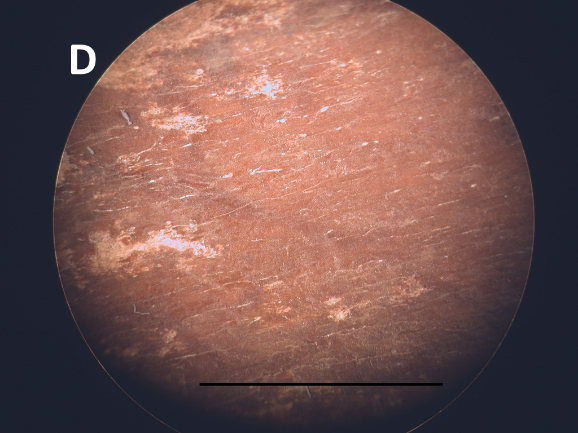 |
| 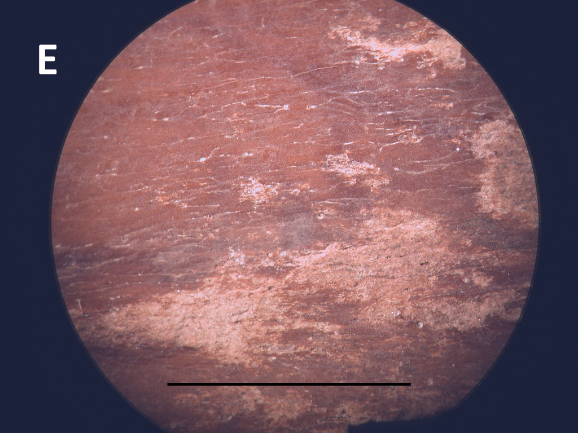 | 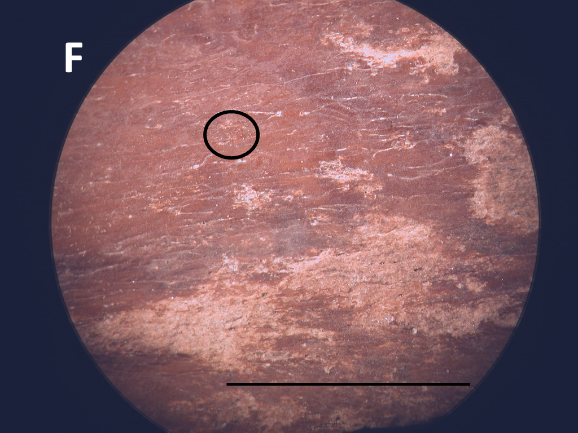 |
| 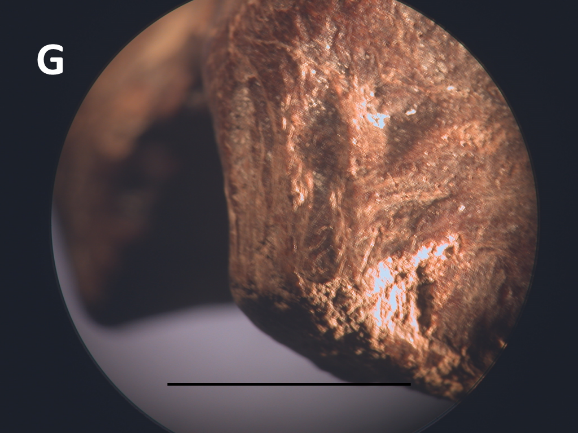 | 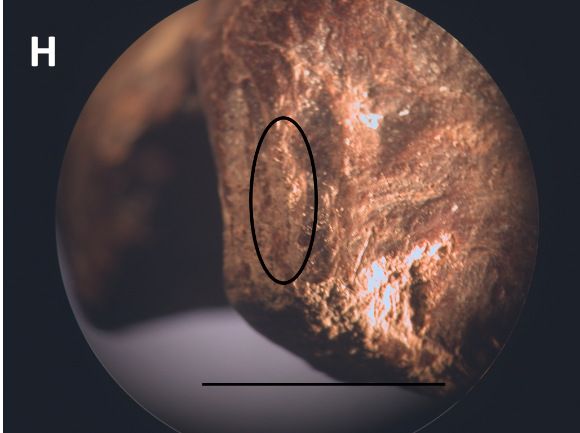 |
| 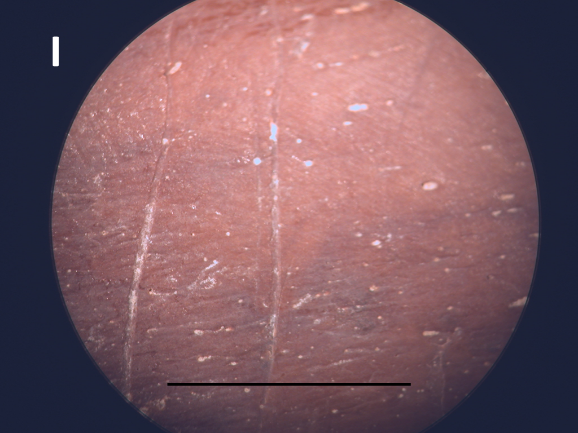 | 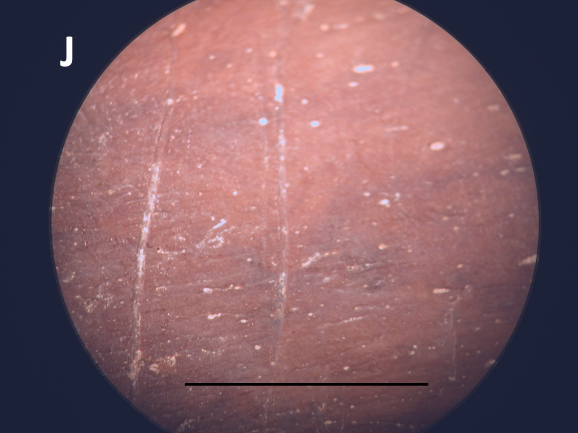 |

**Supplementary Figure S6.** Optical microscopy (magnification ×4, Optika B-383PLi, Italy) sampling method on wild boar, *Sus scrofa* tibia (Bou 1). A, C, E, G, I: before sampling; B, D, F, H, J: after sampling. A, B*:* Triboelectric rubbing in forced bag sampling; C, D: Eraser sampling by rubbing, E, F: Swab mopping sampling, G, H: Conventional destructive sampling, I, J: 5 D-Squame® skin tape-disc. The black circle indicates a modification of the surface after sampling. The black bar represents 1 cm.

**Supplementary Table S2.** Peptide quantity for each method on different bones measured by UV spectrometry at 215 nm with Denovix DS11+. The concentration is in µg/µL.

|  | **Modern** | | | | **Neolithic** | | |
| --- | --- | --- | --- | --- | --- | --- | --- |
|  | **Bos** | **Capra** | **Cervus** | **Struthio** | **Bou 6** | **Bou 2** | **Bou 7** |
| Triboelectric rubbing in forced bag sampling | 0.9 | 0.4 | 0.4 | 0.5 | 0.3 | 0.3 | 0.2 |
| Eraser sampling by rubbing | 1.2 | 1.5 | 1.6 | 1.8 | 1.2 | 1.2 | 1.3 |
| Swab mopping sampling | 2 | 1.8 | 1.9 | 2.6 | 2 | 1.5 | 1.5 |
| Ammonium bicarbonate buffer | 2 | 1.9 | 2.1 | 2.2 | 1.8 | 2.1 | 1.6 |
| Conventional destructive, acid fraction | 1.3 | 1.2 | 1.3 | 1.4 | 1.0 | 1.1 | 0.9 |
| Conventional destructive, bone powder | 6.1 | 4.5 | 2.9 | 3.5 | 3.1 | 2.8 | 3.8 |
| 1 Dermatological skin tape-discs sampling | 1.2 | 0.9 | 1.1 | 1 | 0.8 | 0.6 | 0.5 |
| 5 Dermatological skin tape-discs sampling | 2.3 | 3 | 2.5 | 3 | 2.3 | 2.8 | 2.1 |

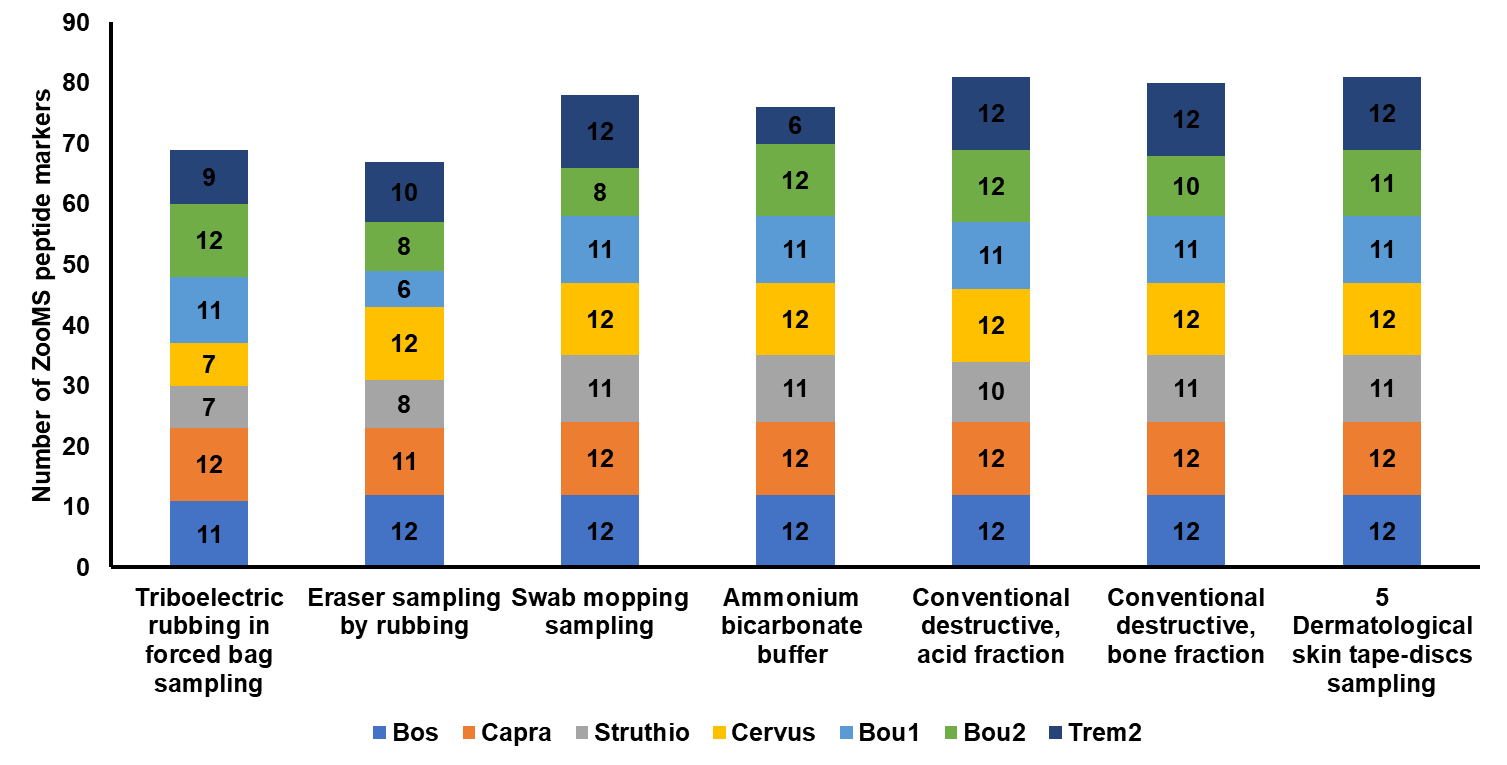

**Supplementary Figure S7.** Number of ZooMS peptide markers for modern (4 samples) and Neolithic (3 samples) for each tested method. For each sample the number of ZooMS peptides markers are indicated. The axe represents the cumulative numbers of peptides markers and the represent the six methods.

**Supplementary Table S3.** Mass to charge ratio (m/z) of the native and deamidated universal peptides. The table contains the peptide sequence, the post-translational modifications, the formula, the [M+H]^+^ of peptides using for MALDI FTICR deamidation values and the [M+2H]^2+^ of peptides using for LC-MS/MS deamidation values.

| Sequence | PTMs | Formula | [M+H]^+^ for MALDI FTICR | [M+2H]^2+^ for LC-MS/MS |
| --- | --- | --- | --- | --- |
| GVQGP^ox^PGPAGPR | 1 oxidation | C_47_H_76_N_16_O_15_ | 1,105.5748 | 553.2910 |
| GVQ^dem^GP^ox^PGPAGPR | 1 oxidation  1 deamidation | C_47_H_75_N_15_O_16_ | 1,106.5580 | 553.7830 |
| GVQGP^ox^PGPQGPR | 1 oxidation | C_49_H_79_N_17_O_16_ | 1162.5963 | 581.8018 |
| GVQ^dem^GP^ox^PGPQGPR | 1 oxidation  1 deamidation | C_49_H_78_N_16_O_17_ | 1163.5803 | 582.2938 |

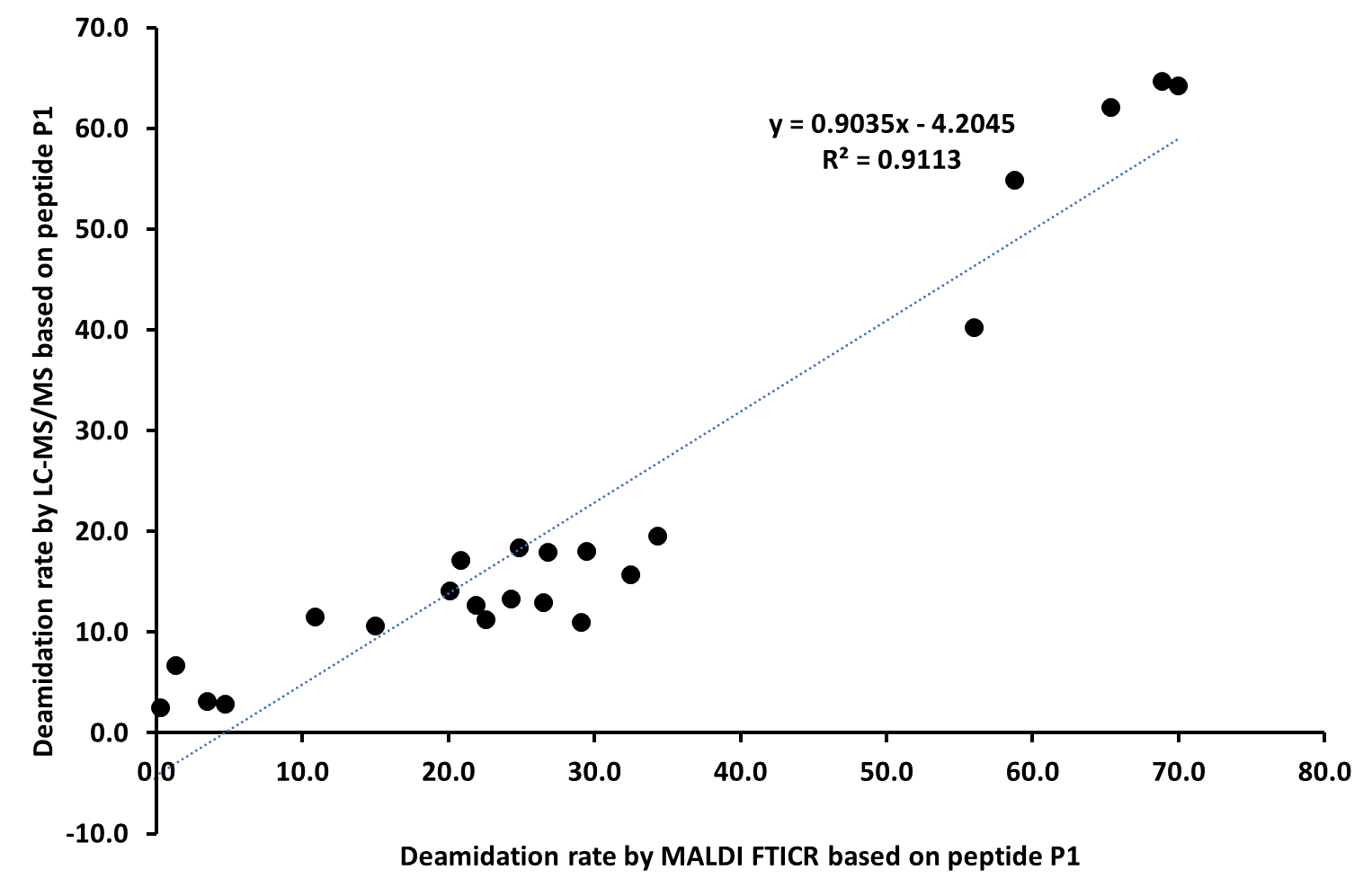

**Supplementary Figure S8.** Percentages of glutamine deamidation by MALDI FTICR and LC-MS/MS based on the peptide P1 sequence from COL1A1 GVQ^dem^GP^ox^PGPAGPR for mammals and GVQ^dem^GP^ox^PGPQGPR for birds. Each of the 23 points corresponds to a sample analysis (4 modern and 19 archeological samples).

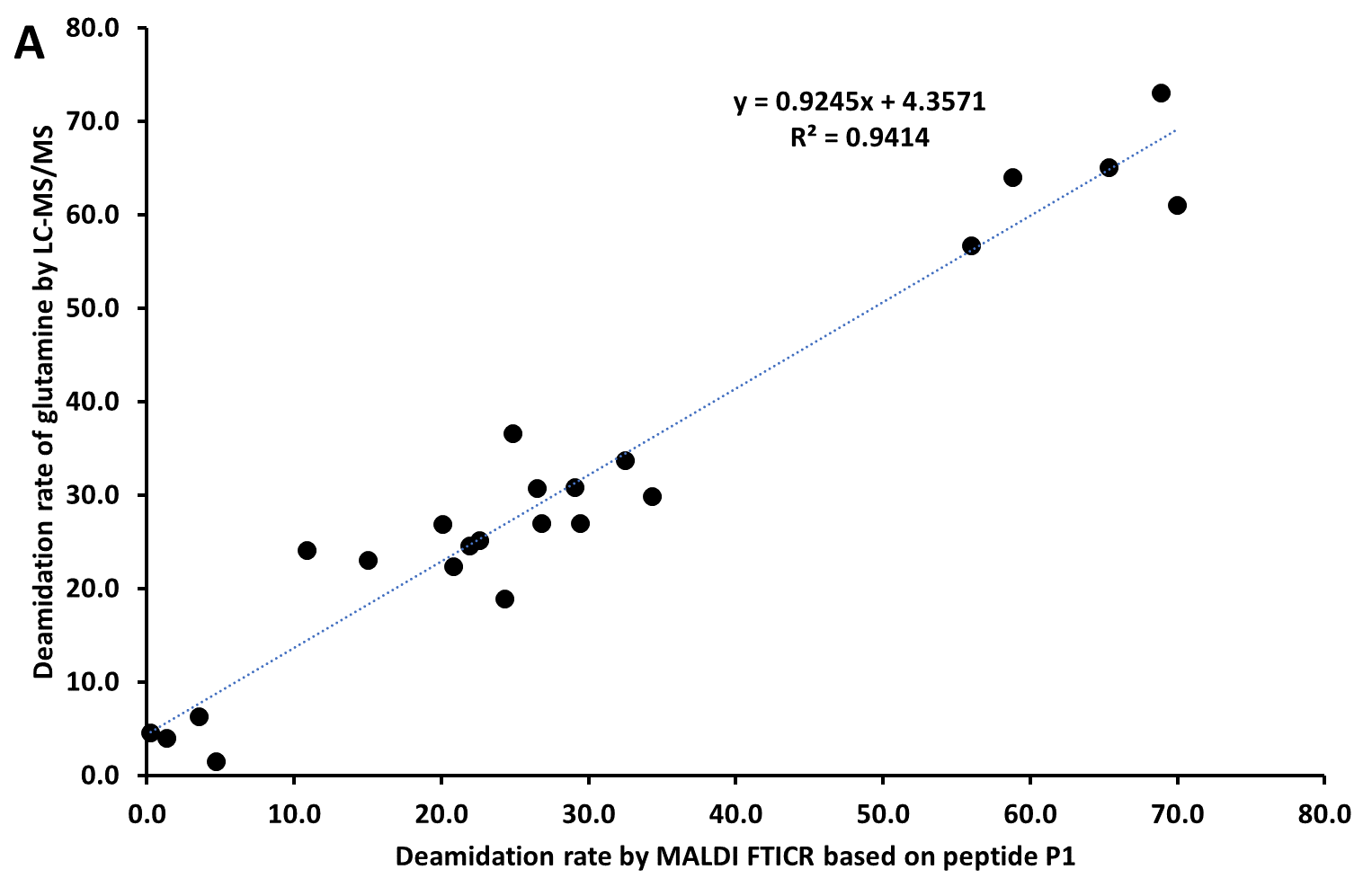

**Supplementary Figure S9.** Percentages of glutamine deamidation by MALDI FTICR based on the peptide P1 sequence from COL1A1 GVQ^dem^GP^ox^PGPAGPR for mammals and GVQ^dem^GP^ox^PGPQGPR for birds and the percentages of glutamine deamidation LC-MS/MS based on Maxquant and the deamidation package.

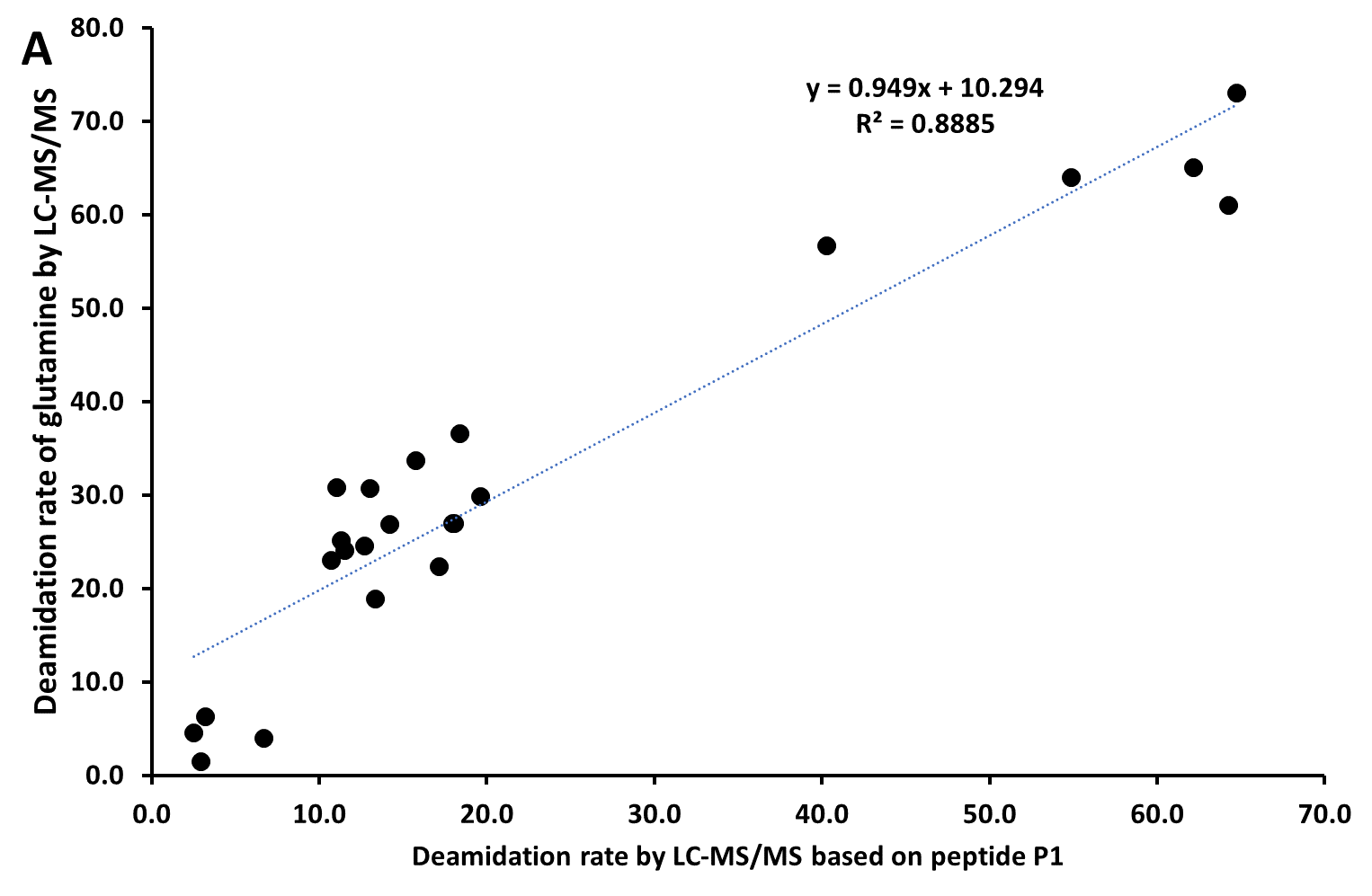

**Supplementary Figure S10.** Percentages of glutamine deamidation by LC-MS/MS based on the peptide P1 sequence from COL1A1 GVQ^dem^GP^ox^PGPAGPR for mammals and GVQ^dem^GP^ox^PGPQGPR for birds and the percentages of glutamine deamidation LC-MS/MS based on Maxquant and the deamidation package.

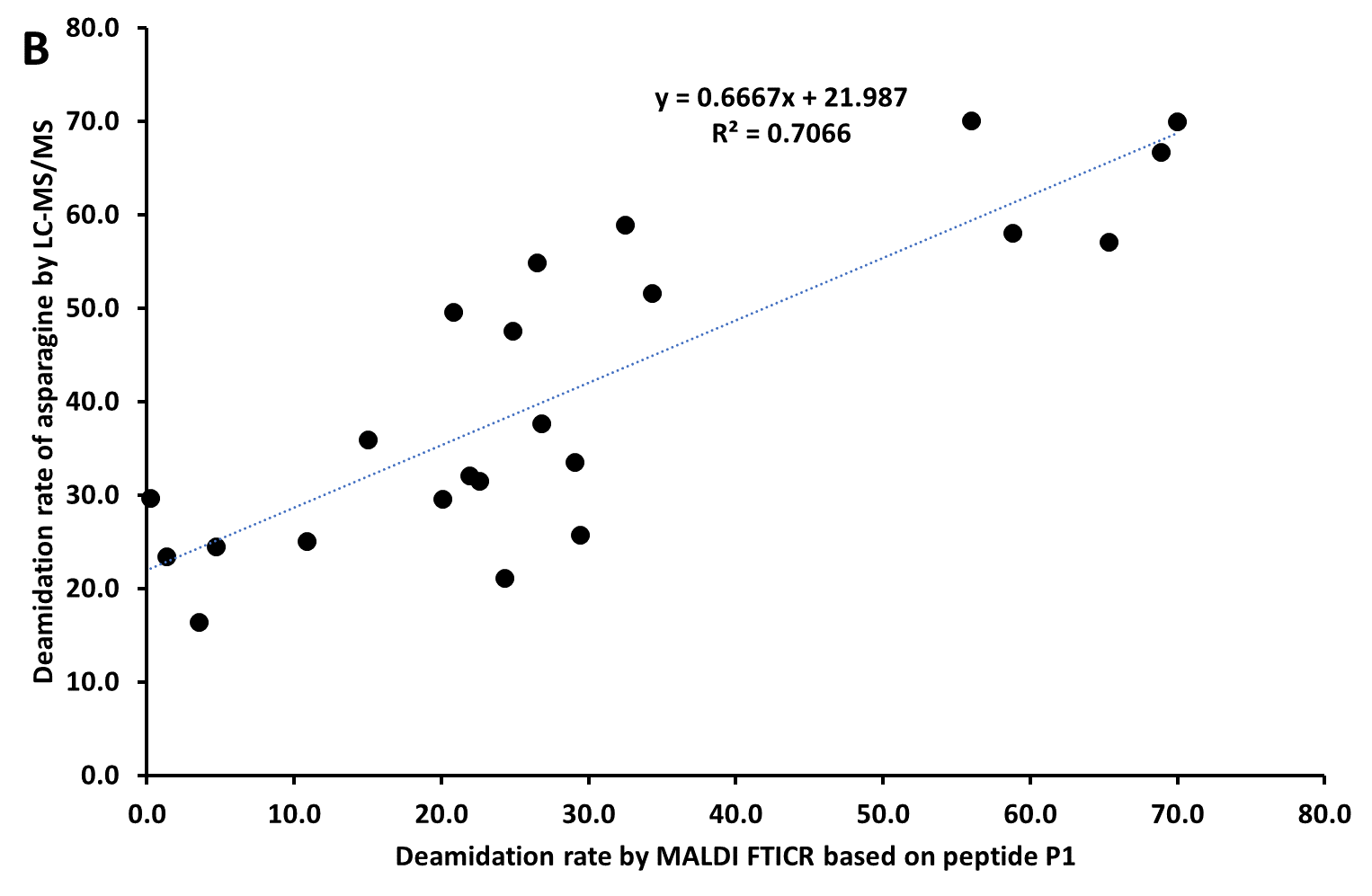

**Supplementary Figure S11.** Percentages of glutamine deamidation by MALDI FTICR based on the peptide P1 sequence from COL1A1 GVQ^dem^GP^ox^PGPAGPR for mammals and GVQ^dem^GP^ox^PGPQGPR for birds and the percentages of asparagine deamidation LC-MS/MS based on Maxquant and the deamidation package.

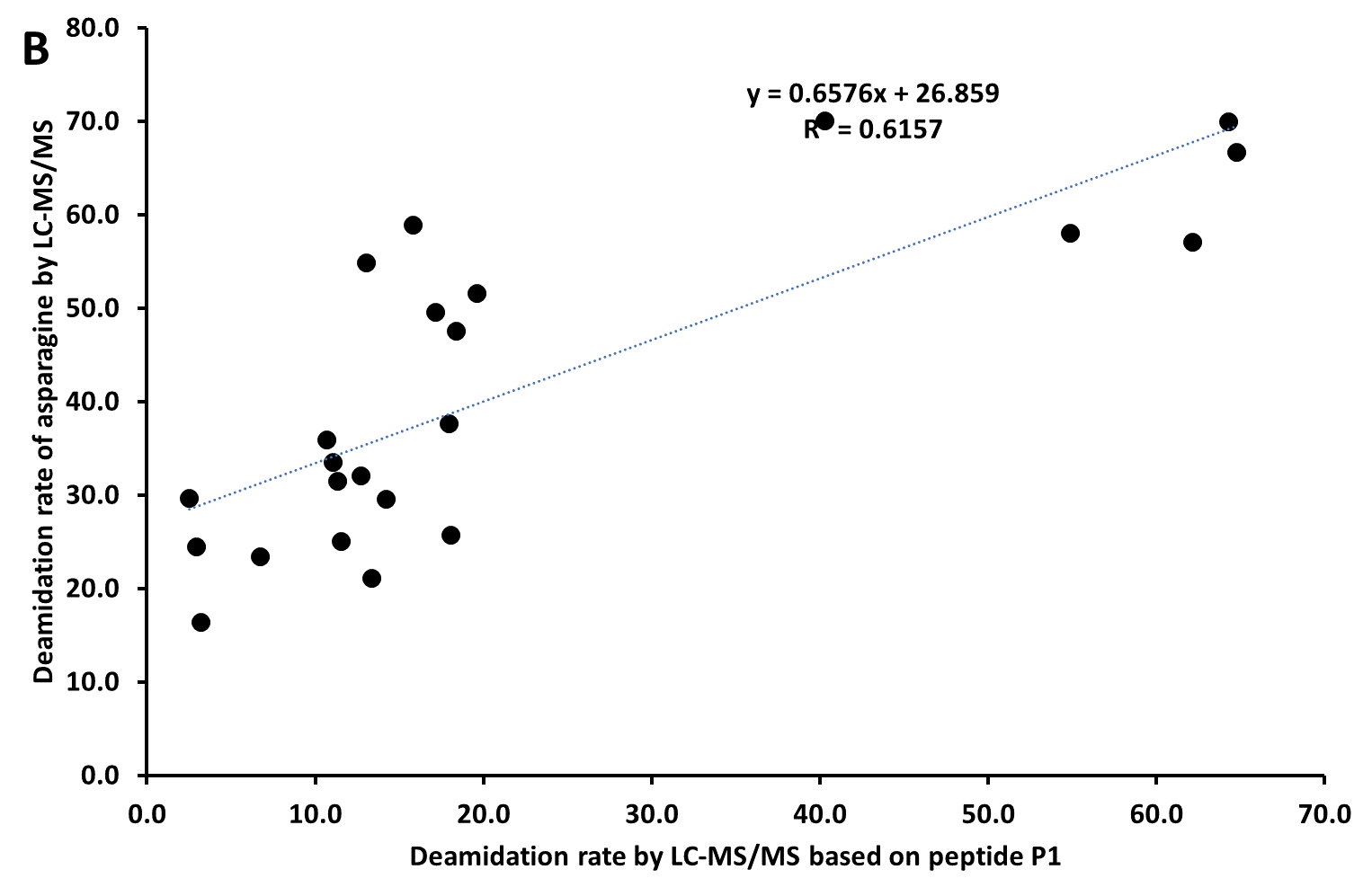

**Supplementary Figure S12.** Percentages of glutamine deamidation by LC-MS/MS based on the peptide P1 sequence from COL1A1 GVQ^dem^GP^ox^PGPAGPR for mammals and GVQ^dem^GP^ox^PGPQGPR for birds and the percentages of asparagine deamidation LC-MS/MS based on Maxquant and the deamidation package.

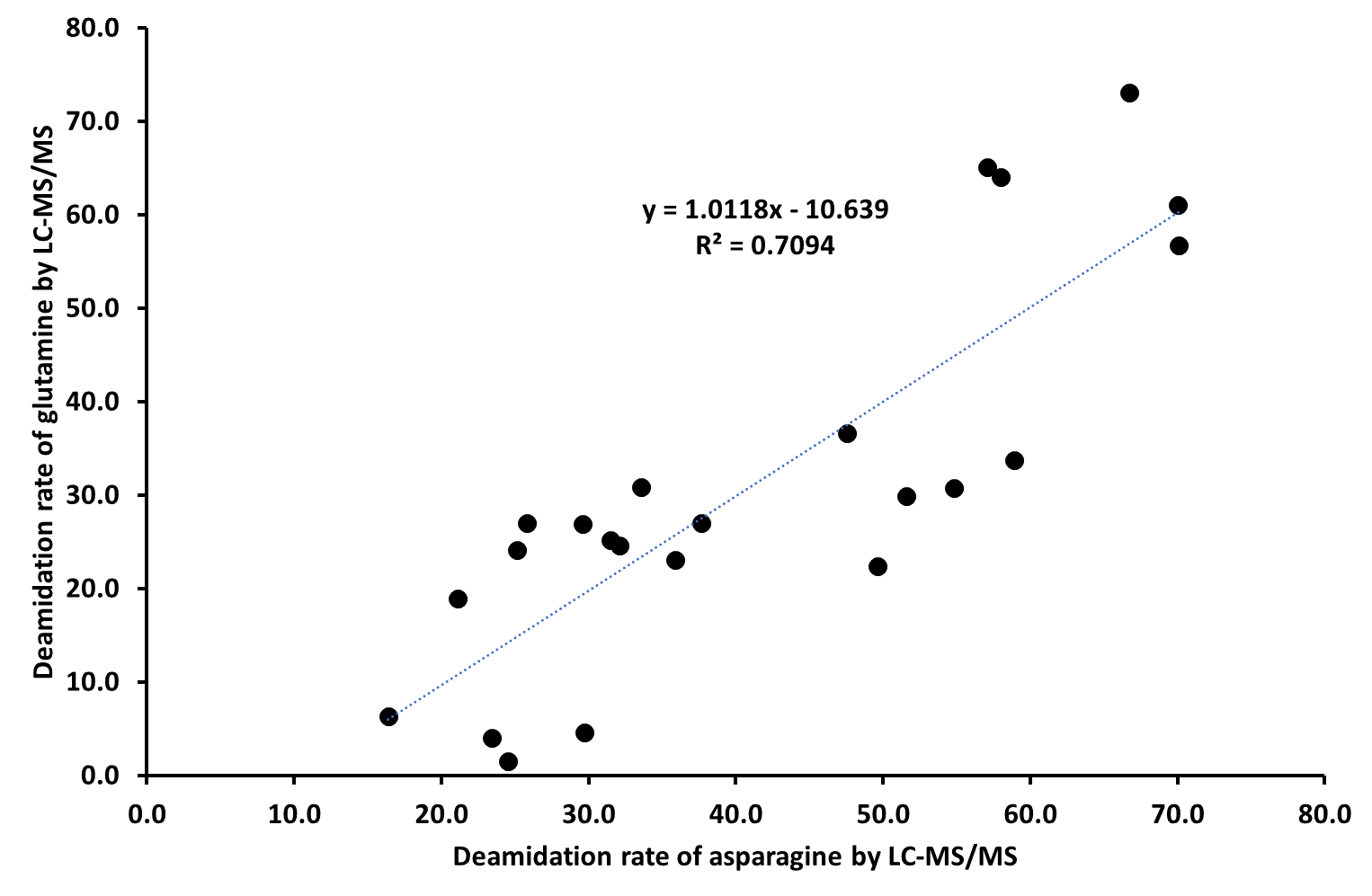

**Supplementary Figure S13.** Global percentages of asparagine and glutamine deamidation by LC-MS/MS based on on Maxquant and the “deamidation” package.
