## Supporting Information 3 contains MS/MS spectra of peptides obtained by LC-MS/MS for "Low-invasive sampling method for taxonomic for the identification of archaeological and paleontological bones by proteomics of their collagens"

**Table content**

[**Supplementary Figure S1**. MS/MS Fragmentation of GVQGPP^ox^GPAGPR found in AAI05185.1 in NCBi_Mamm, collagen alpha-1(I) chain precursor [Bos taurus]	5](#_Toc146114062)

[**Supplementary Figure S*2***. MS/MS Fragmentation of GVQ^dem^GPP*^ox^*GPAGPR found in AAI05185.1 in NCBi_Mamm, collagen alpha-1(I) chain precursor [*Bos taurus*]	5](#_Toc146114063)

[**Supplementary Figure S3**. MS/MS Fragmentation of GVQGPP^ox^GPQGPR from XP_009685373.1 in NCBi_Aves, collagen alpha-1(I) chain precursor [Struthio camelus]	6](#_Toc146114064)

[**Supplementary Figure S4**. MS/MS Fragmentation of GVQ^dem^GPP^ox^GPQGPR from XP_009685373.1 in NCBi_Aves, collagen alpha-1(I) chain precursor [Struthio camelus]	6](#_Toc146114065)

[**Supplementary Figure S5**. MS/MS Fragmentation of GEVGLP^ox^GLSGPVGPP^ox^GNP^ox^GANGLAGSK found in JAV39576.1 in NCBi_Mamm, collagen alpha-2(I) chain precursor [Castor canadensis]	7](#_Toc146114066)

[**Supplementary Figure S6**. MS/MS Fragmentation of GLVGEP^ox^GPAGSKGETGSK found in JAV39576.1 in NCBi_Mamm, collagen alpha-2(I) chain precursor [Castor canadensis]	7](#_Toc146114067)

[**Supplementary Figure S7**. MS/MS Fragmentation of GNDGSVGPVGPAGPIGSAGPP^ox^GFP^ox^GAP^ox^GPK found in BAX02569.1 in NCBi_Mamm, alpha2 chain of type I collagen [Sus scrofa domesticus]	8](#_Toc146114068)

[**Supplementary Figure S8**. MS/MS Fragmentation of GEVGLP^ox^GVSGPVGPP^ox^GNP^ox^GANGLPGAK found in BAX02569.1 in NCBi_Mamm, alpha2 chain of type I collagen [Sus scrofa domesticus]	8](#_Toc146114069)

[**Supplementary Figure S9.** MS/MS Fragmentation of GAP^ox^GAQ^dem^GPP^ox^GAP^ox^GPLGIAGVTGAR found in XP_043343785.1 in NCBi_Mamm, collagen alpha-1(III) chain [Cervus canadensis]	9](#_Toc146114070)

[**Supplementary Figure S10**. MS/MS Fragmentation of GDAGP^ox^PGPAGPAGPPGPIGSVGAPGP^ox^K found in sp|C0HJN9.1|CO1A1_EQUSP in NCBi_Mamm, RecName: Full=Collagen alpha-1(I) chain; AltName: Full=Alpha-1 type I collagen [Equus caballus]	9](#_Toc146114071)

[**Supplementary Figure S11.** MS/MS Fragmentation of TGEP^ox^GAAGPP^ox^GFVGEK found in XP_004007775.1 in NCBi_Mamm, collagen alpha-2(I) chain [Ovis aries]	10](#_Toc146114072)

[**Supplementary Figure S12.** MS/MS Fragmentation of GEP^ox^GPVGAVGPAGAVGPR found in XP_004007775.1 in NCBi_Mamm, collagen alpha-2(I) chain [Ovis aries]	10](#_Toc146114073)

[**Supplementary Figure S13**. MS/MS Fragmentation of SGETGASGPP^ox^GFVGEK found in NP_776945.1 in NCBi_Mamm, RecName: collagen alpha-2(I) chain precursor [Bos taurus]	11](#_Toc146114074)

[**Supplementary Figure S14.** MS/MS Fragmentation of GLHGEFGAP^ox^GPAGPR found in XP_009672566.1 in NCBI_Aves, PREDICTED: collagen alpha-2(I) chain isoform X1 [Struthio camelus australis]	11](#_Toc146114075)

[**Supplementary Figure S15.** MS/MS Fragmentation of TGEP^ox^GAAGPP^ox^GFVGEKGP^ox^SGEPGTAGPPGTPGPQGFLGPP^ox^GFLGLP^ox^GSR found in XP_005678993.1 in NCBi_Mamm, PREDICTED: collagen alpha-2(I) chain [Capra hircus]	12](#_Toc146114076)

[**Supplementary Figure S16.** MS/MS Fragmentation of GESGN^dem^KGEP^ox^GSVGPQ^dem^GPP^ox^GPSGEEGK found in XP_008684476.1 in NCBi_Mamm, PREDICTED: collagen alpha-2(I) chain [Ursus maritimus]	12](#_Toc146114077)

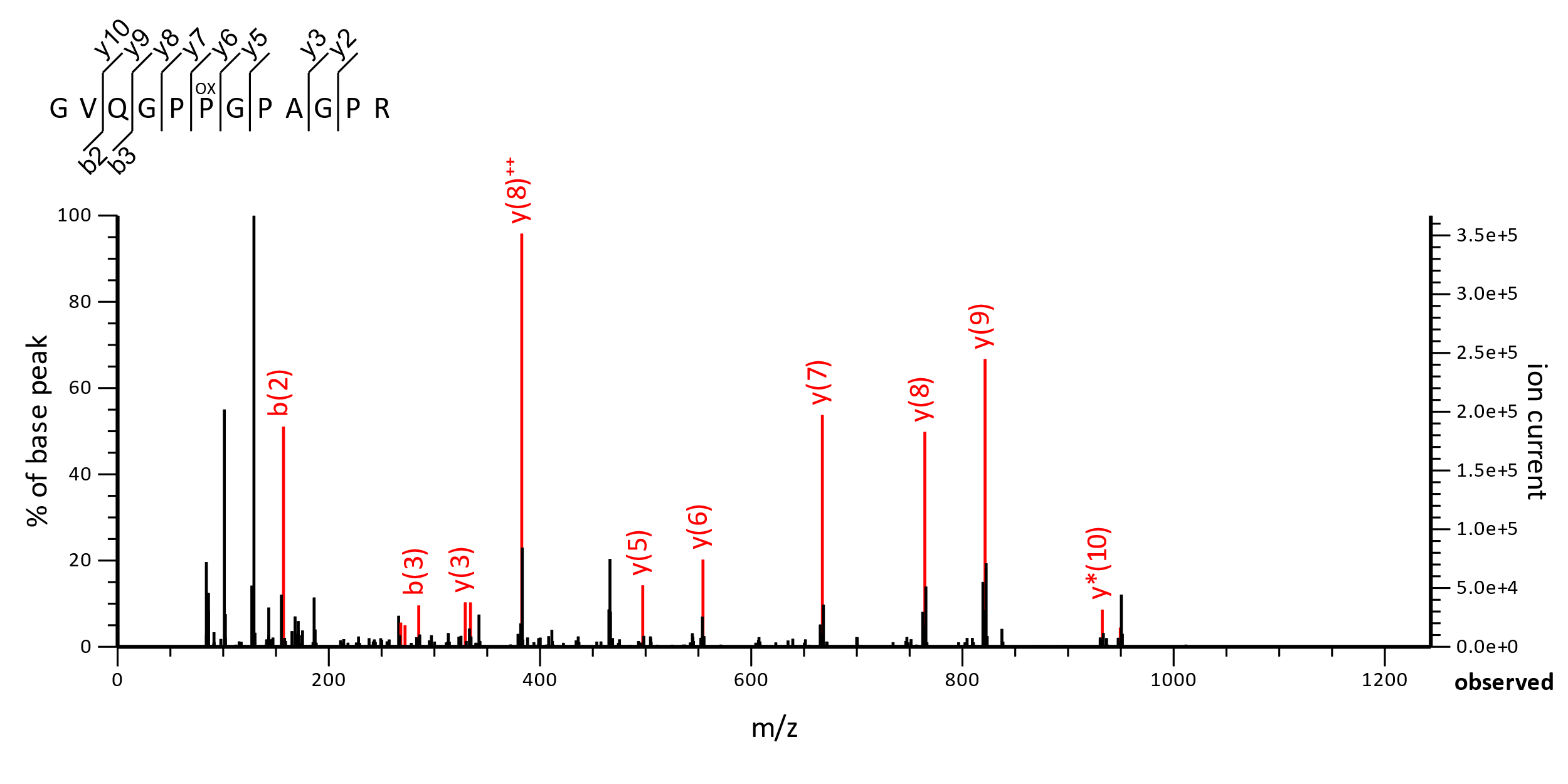

**Supplementary Figure S1**. MS/MS Fragmentation of GVQGPP^ox^GPAGPR found in AAI05185.1 in NCBi_Mamm, collagen alpha-1(I) chain precursor [Bos taurus]

Match to Query 3580: 1104.568456 from (553.291504,2+) intensity (3836956.9609) rtinseconds (2580.99978) index (6717)

Monoisotopic mass of neutral peptide Mr(calc): 1104.5676

Variable modifications: P6 : Oxidation (P), Ions Score: 39 Expect: 0.00075

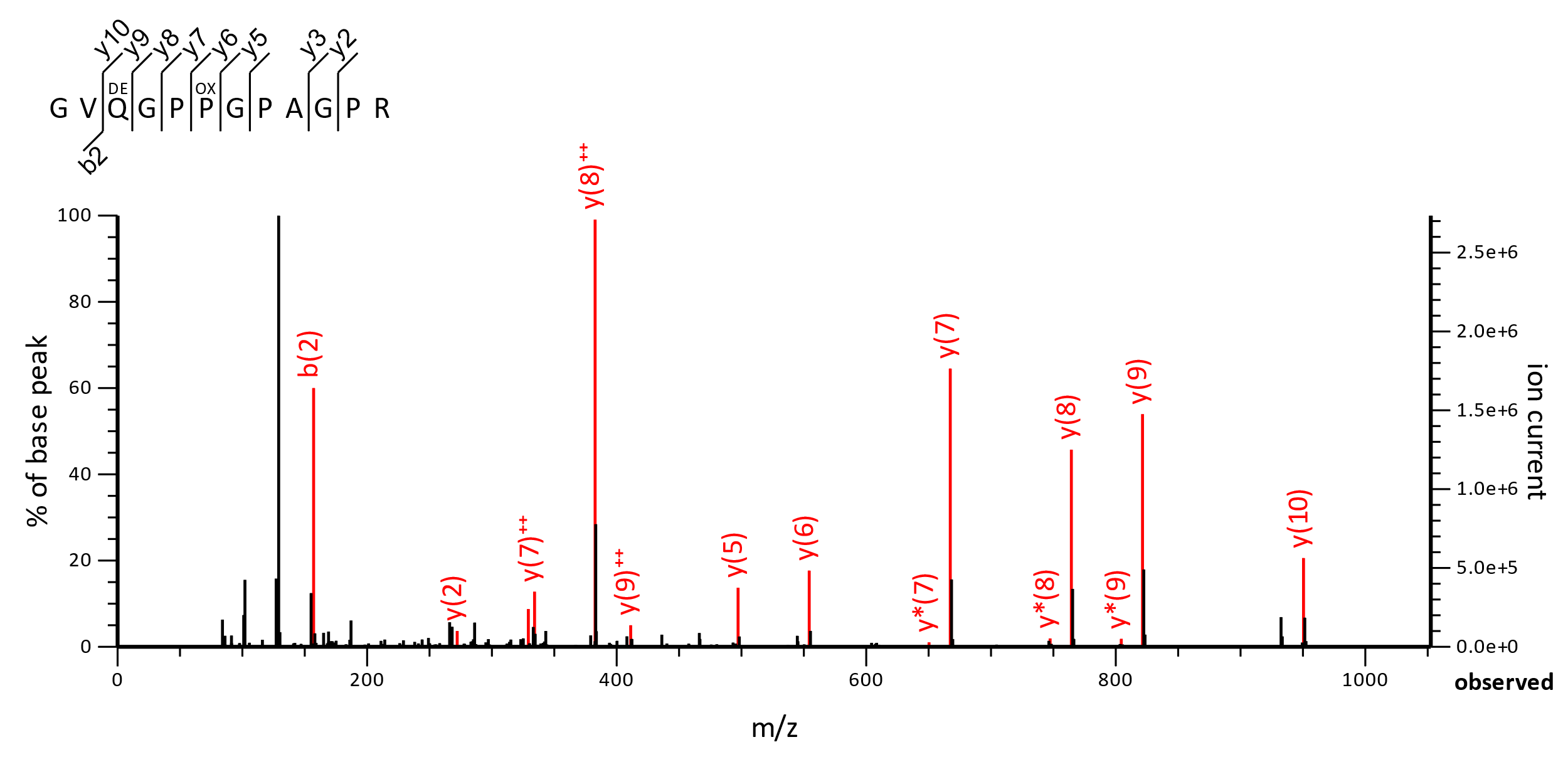
**Supplementary Figure S*2***. MS/MS Fragmentation of GVQ^dem^GPP*^ox^*GPAGPR found in AAI05185.1 in NCBi_Mamm, collagen alpha-1(I) chain precursor [*Bos taurus*]

Match to Query 3595: 1105.551718 from (553.783135,2+) intensity (30251253.3555) rtinseconds (2559.30732) index (6606)

Monoisotopic mass of neutral peptide Mr(calc): 1105.5516

Variable modifications: Q3 : Deamidated (NQ), P6 : Oxidation (P), Ions Score: 43 Expect: 0.0021

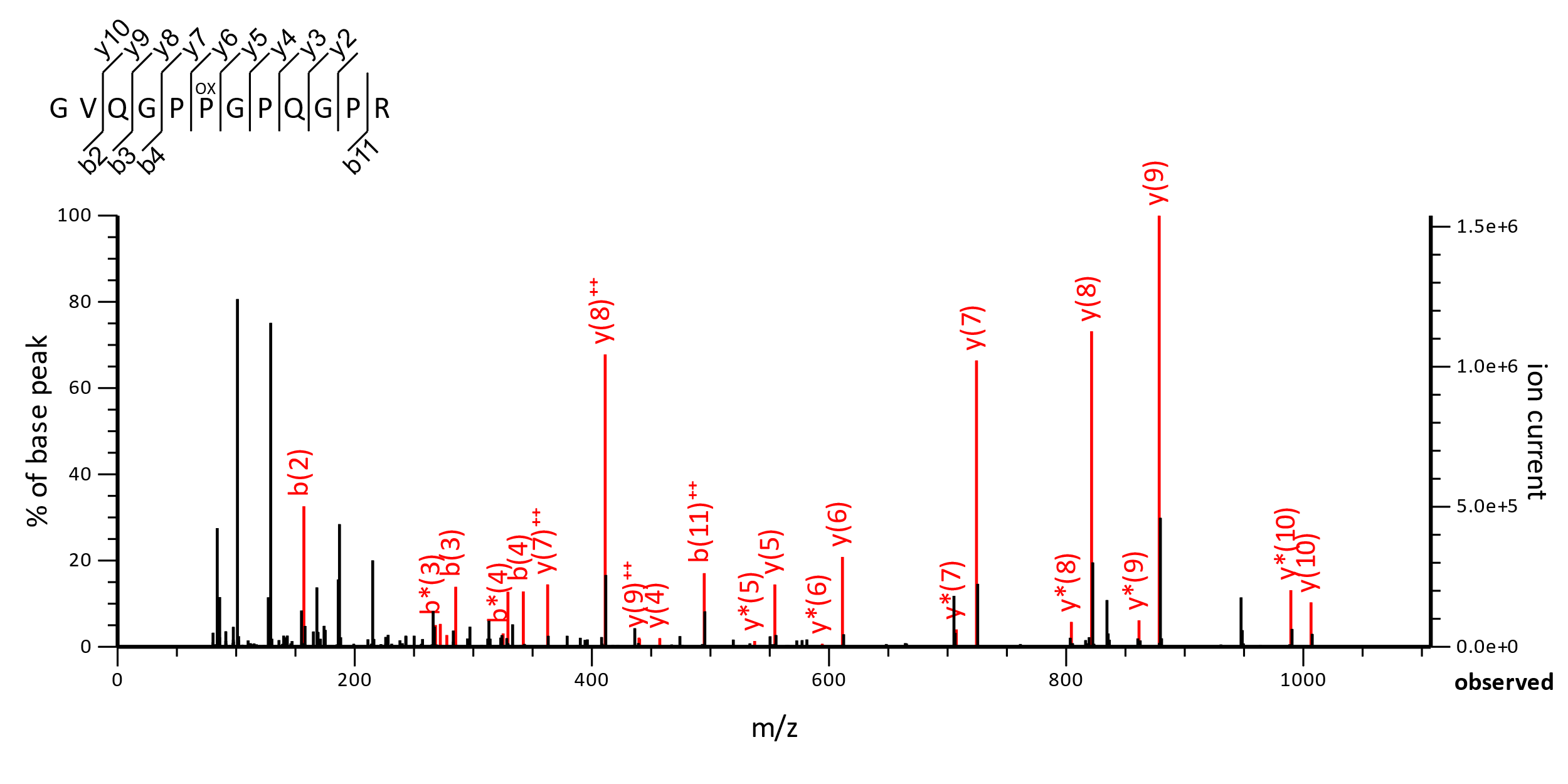
**Supplementary Figure S3**. MS/MS Fragmentation of GVQGPP^ox^GPQGPR from XP_009685373.1 in NCBi_Aves, collagen alpha-1(I) chain precursor [Struthio camelus]

Match to Query 7536: 1161.589458 from (581.802005,2+) intensity (49137275.2266) rtinseconds (1738.7082) index(4717)

Monoisotopic mass of neutral peptide Mr(calc): 1161.5891 Variable modifications: P6 : Oxidation (P), Ions Score: 37 Expect: 0.0036

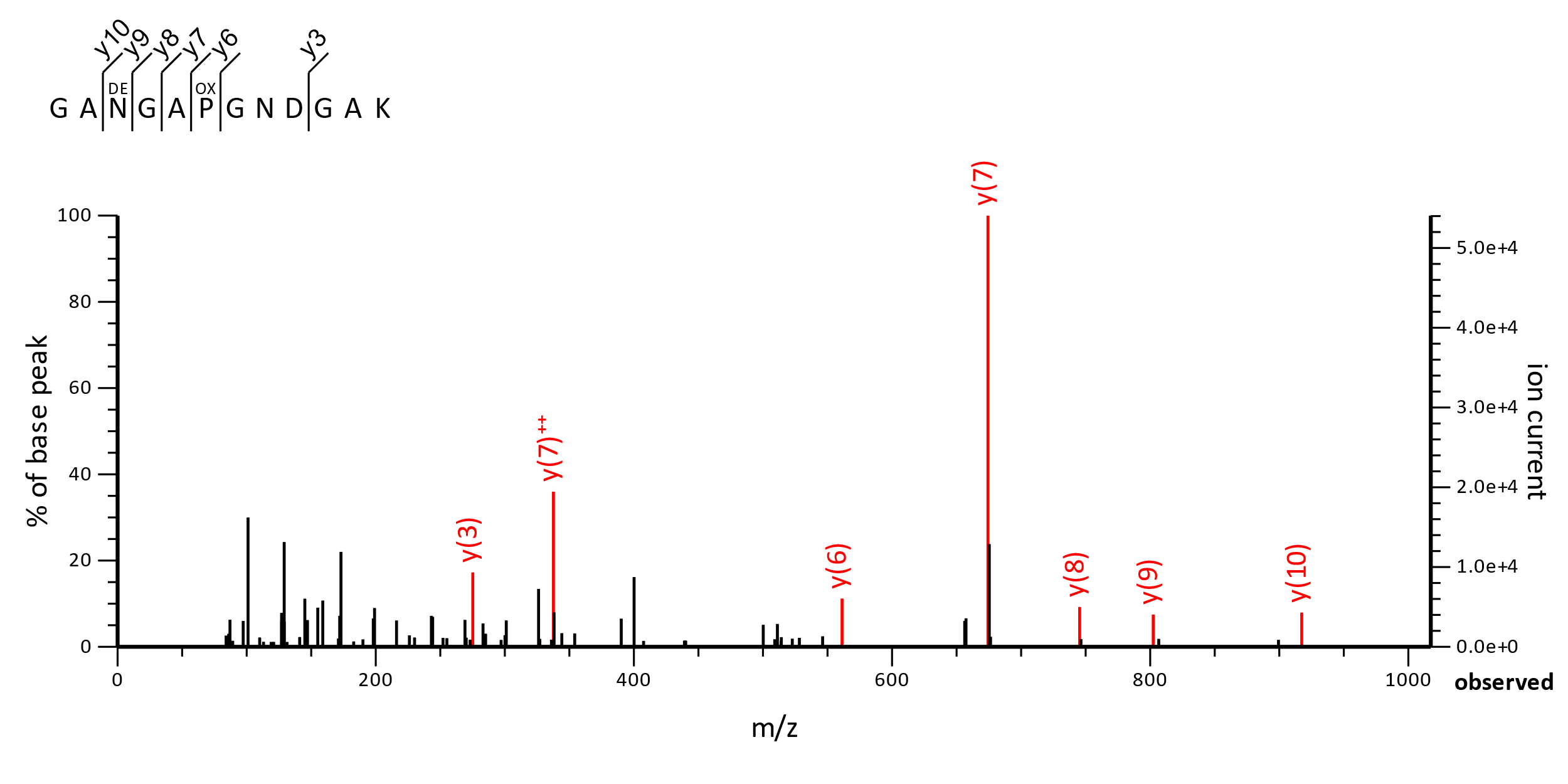
**Supplementary Figure S4**. MS/MS Fragmentation of GVQ^dem^GPP^ox^GPQGPR from XP_009685373.1 in NCBi_Aves, collagen alpha-1(I) chain precursor [Struthio camelus]

Match to Query 5086: 1044.445320 from (523.229936,2+) intensity (993099.7197) rtinseconds (1202.8476) index (1341)

Monoisotopic mass of neutral peptide Mr(calc): 1044.4472

Variable modifications: N3: Deamidated (NQ), P6: Oxidation (P), Ions Score: 49 Expect: 0.016

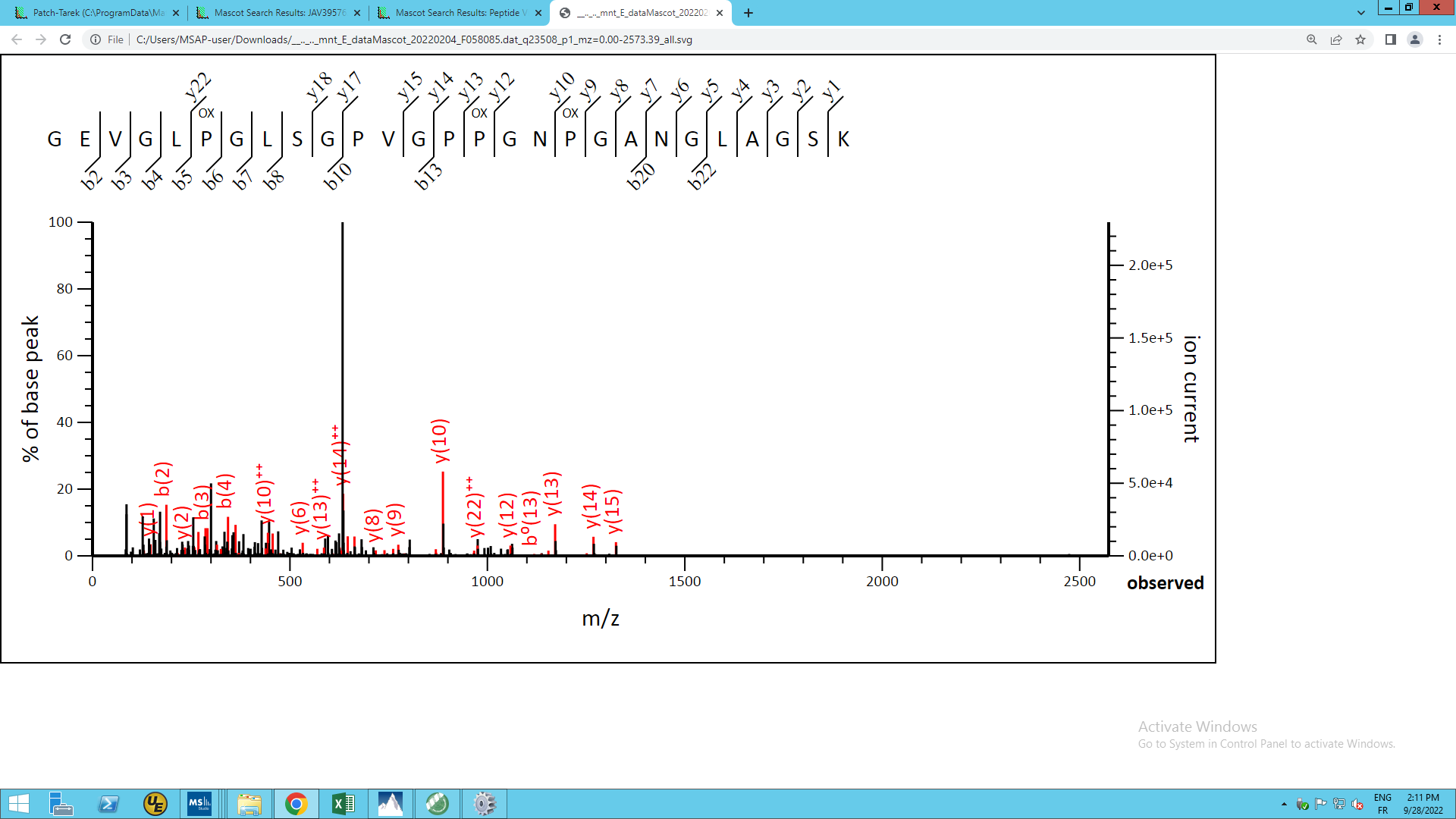

**Supplementary Figure S5**. MS/MS Fragmentation of GEVGLP^ox^GLSGPVGPP^ox^GNP^ox^GANGLAGSK found in JAV39576.1 in NCBi_Mamm, collagen alpha-2(I) chain precursor [Castor canadensis]

Match to Query 23508: 2403.213642 from (802.078490,3+) intensity (1371103.2500) rtinseconds (5108) scans (27653) index (22179)

Monoisotopic mass of neutral peptide Mr (calc): 2403.2030

Fixed modifications : Carbamidomethyl (C) (apply to specified residues or termini only)

Variable modifications : P6 : Oxidation (P) P15 : Oxidation (P) P18 : Oxidation (P) Ions Score : 42 Expect : 0.00049.

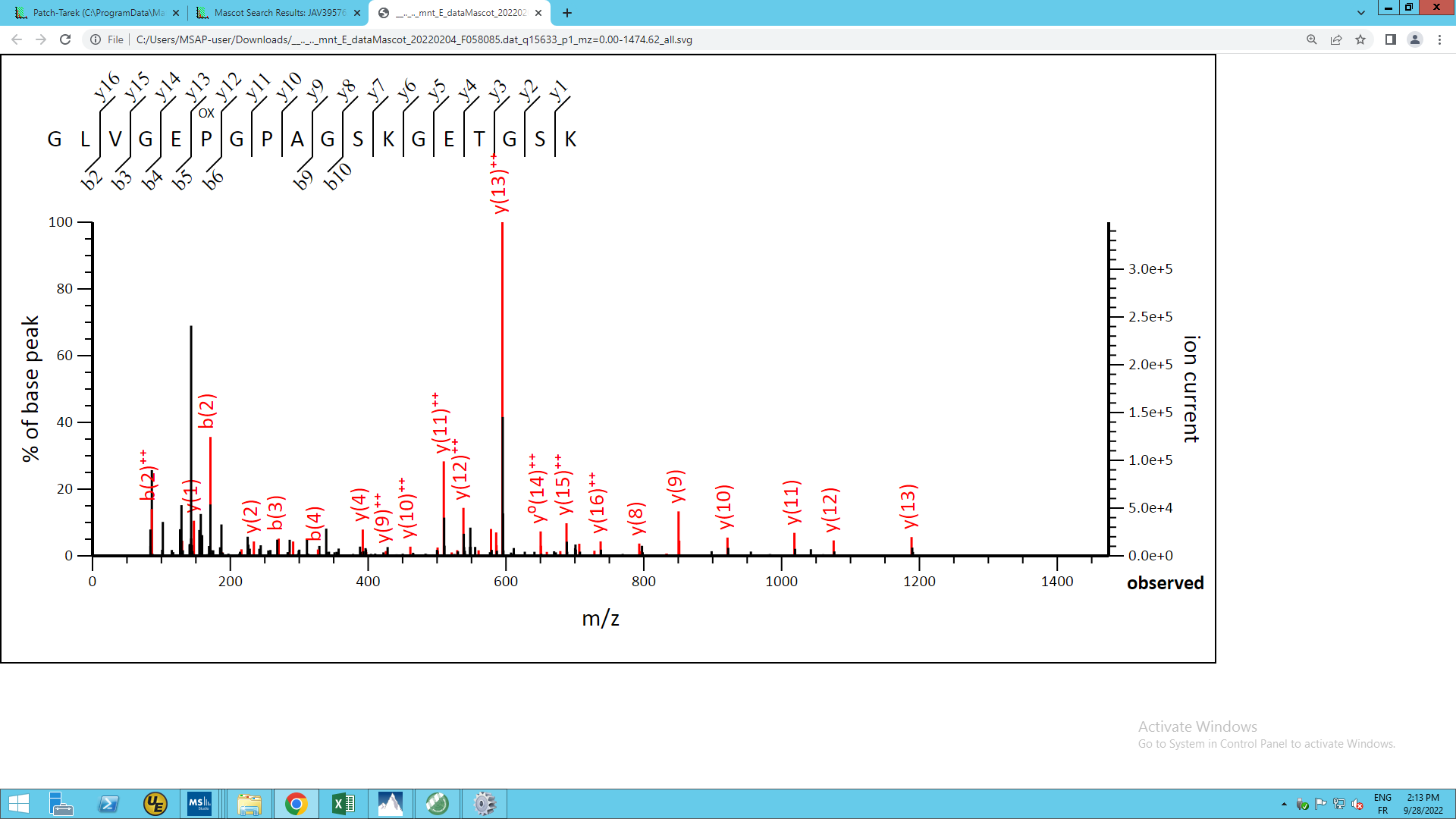

**Supplementary Figure S6**. MS/MS Fragmentation of GLVGEP^ox^GPAGSKGETGSK found in JAV39576.1 in NCBi_Mamm, collagen alpha-2(I) chain precursor [Castor canadensis]

Match to Query 15633: 1642.817652 from (548.613160,3+) intensity (2182462.0000) rtinseconds (2121) scans (8422) index (4468)

Monoisotopic mass of neutral peptide Mr (calc): 1642.8162

Fixed modifications : Carbamidomethyl (C) (apply to specified residues or termini only)

Variable modifications : P6 : Oxidation (P) Ions Score: 53 Expect : 7.1e-005.

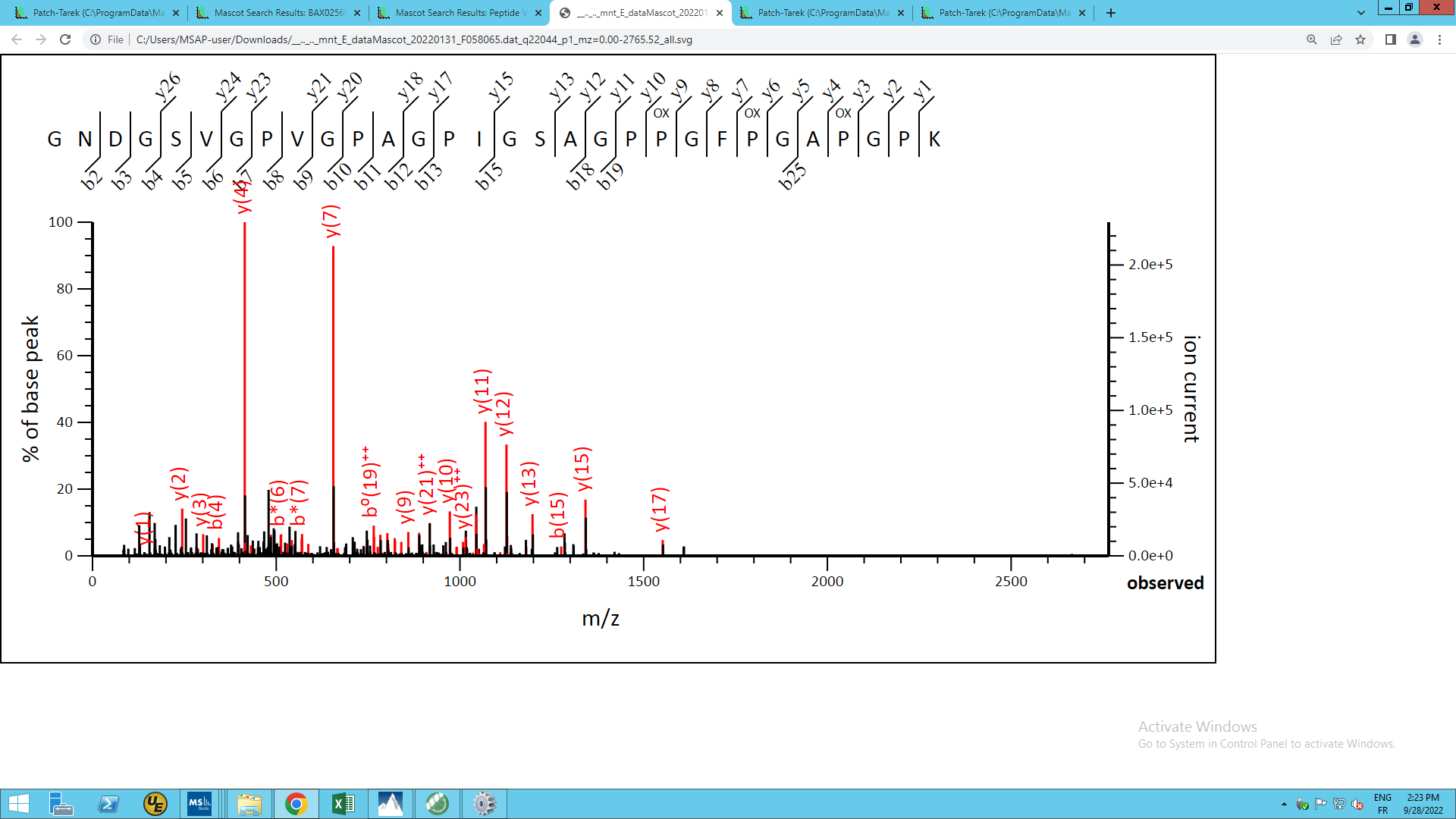

**Supplementary Figure S7**. MS/MS Fragmentation of GNDGSVGPVGPAGPIGSAGPP^ox^GFP^ox^GAP^ox^GPK found in BAX02569.1 in NCBi_Mamm, alpha2 chain of type I collagen [Sus scrofa domesticus]

Match to Query 22044: 2615.284528 from (1308.649540,2+) intensity (616256.3125) rtinseconds (4671) scans (24738) index (19359)

Monoisotopic mass of neutral peptide Mr (calc): 2615.2617

Fixed modifications : Carbamidomethyl (C) (apply to specified residues or termini only)

Variable modifications : P21 : Oxidation (P), P24 : Oxidation (P), P27 : Oxidation (P), Ions Score : 76 Expect : 1.9e-007

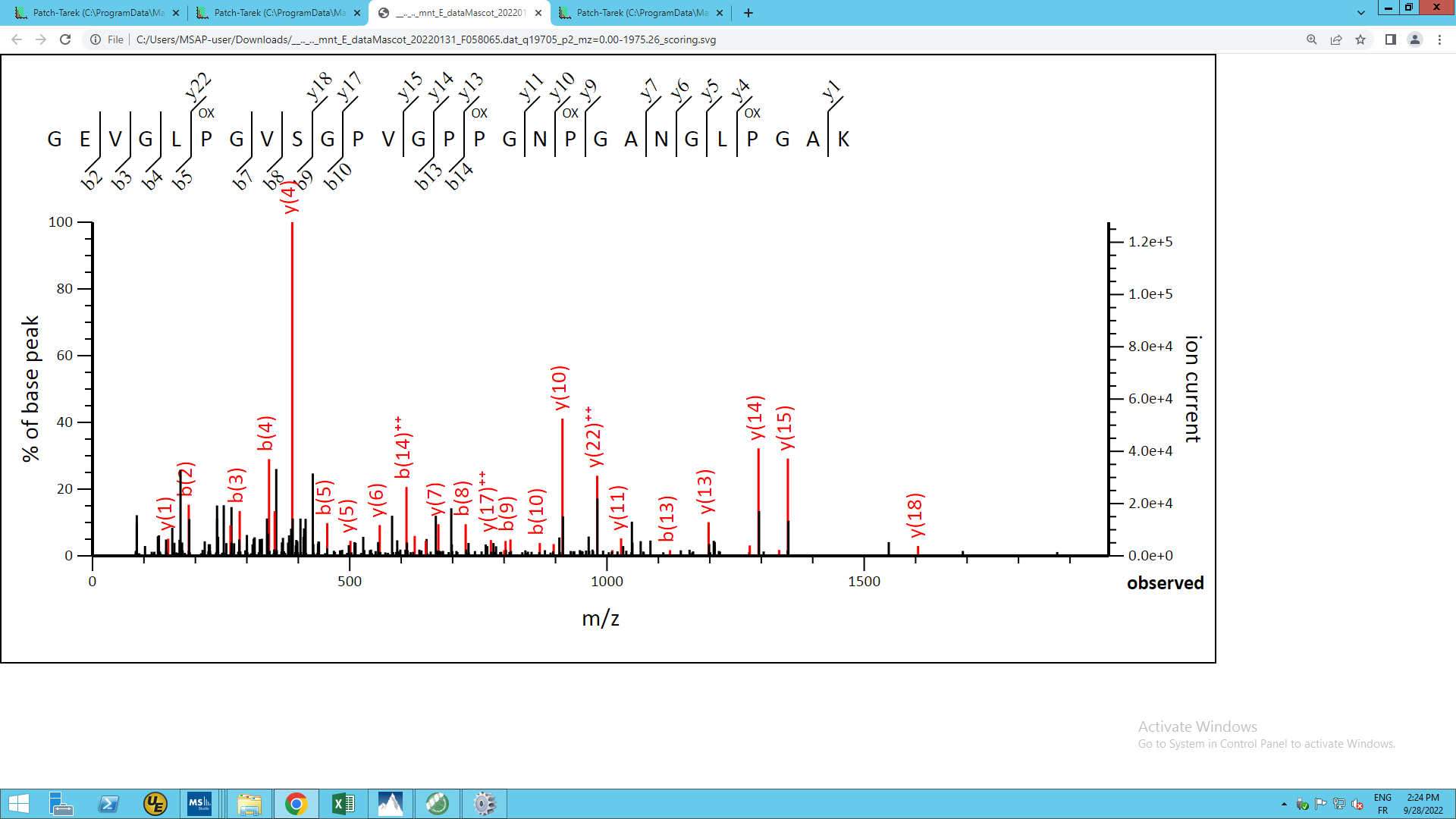

**Supplementary Figure S8**. MS/MS Fragmentation of GEVGLP^ox^GVSGPVGPP^ox^GNP^ox^GANGLPGAK found in BAX02569.1 in NCBi_Mamm, alpha2 chain of type I collagen [Sus scrofa domesticus]

Match to Query 19705: 2415.210048 from (1208.612300,2+) intensity (635233.9375) rtinseconds (4373) scans (22853) index (17734)

Monoisotopic mass of neutral peptide Mr (calc): 2415.2031

Fixed modifications: Carbamidomethyl (C) (apply to specified residues or termini only)

Variable modifications: P6 : Oxidation (P), P15 : Oxidation (P), P18 : Oxidation (P), P24 : Oxidation (P), Ions Scor e: 49 Expect : 2.8e-005

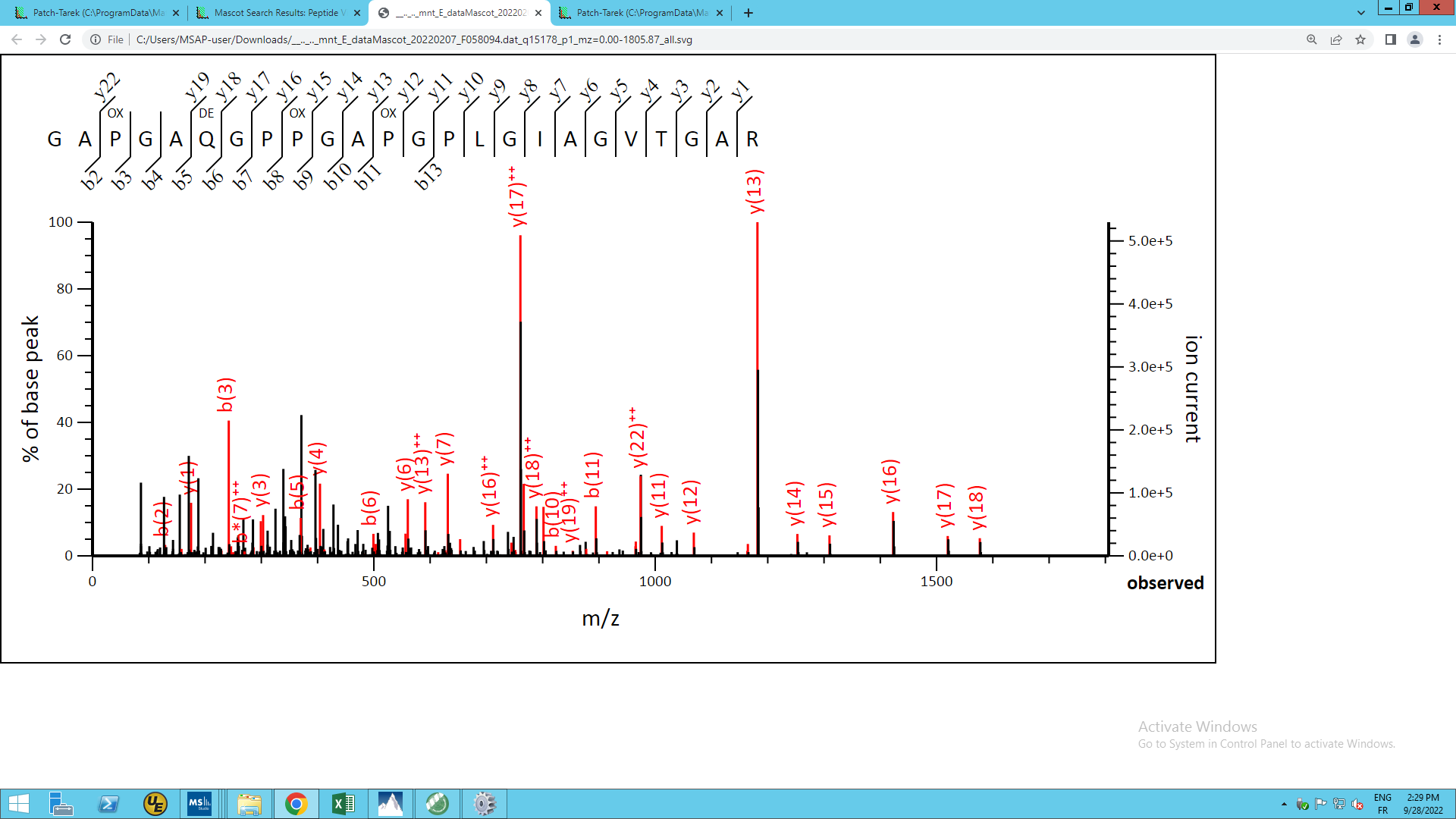

**Supplementary Figure S9.** MS/MS Fragmentation of GAP^ox^GAQ^dem^GPP^ox^GAP^ox^GPLGIAGVTGAR found in XP_043343785.1 in NCBi_Mamm, collagen alpha-1(III) chain [Cervus canadensis]

Match to Query 15178: 2074.051128 from (1038.032840,2+) intensity (3587856.2500) rtinseconds (4777) scans (25815) index (20162)

Monoisotopic mass of neutral peptide Mr(calc): 2074.0444

Fixed modifications : Carbamidomethyl (C) (apply to specified residues or termini only)

Variable modifications : P3 : Oxidation (P), Q6 : Deamidated (NQ), P9 : Oxidation (P)

P12 : Oxidation (P), Ions Score : 72 Expect : 1e-006

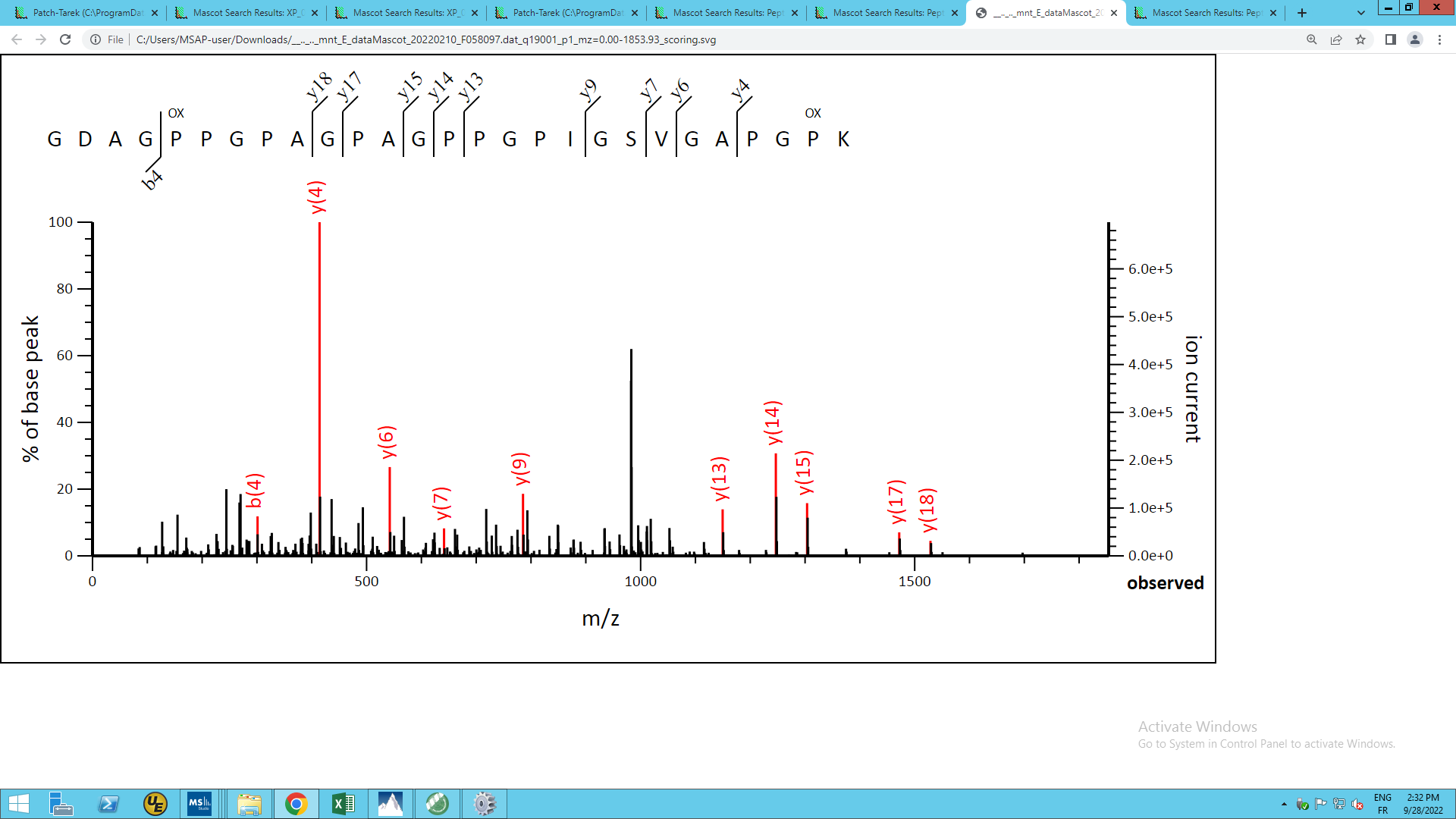

**Supplementary Figure S10**. MS/MS Fragmentation of GDAGP^ox^PGPAGPAGPPGPIGSVGAPGP^ox^K found in sp|C0HJN9.1|CO1A1_EQUSP in NCBi_Mamm, RecName: Full=Collagen alpha-1(I) chain; AltName: Full=Alpha-1 type I collagen [Equus caballus]

Match to Query 19001: 2263.124368 from (1132.569460,2+) intensity (7975591.5000) rtinseconds (4346) scans (22744) index (17398)

Monoisotopic mass of neutral peptide Mr (calc): 2263.1234

Fixed modifications : Carbamidomethyl (C) (apply to specified residues or termini only)

Variable modifications : P5 : Oxidation (P), P26 : Oxidation (P), Ions Score : 54 Expect : 1.8e-005

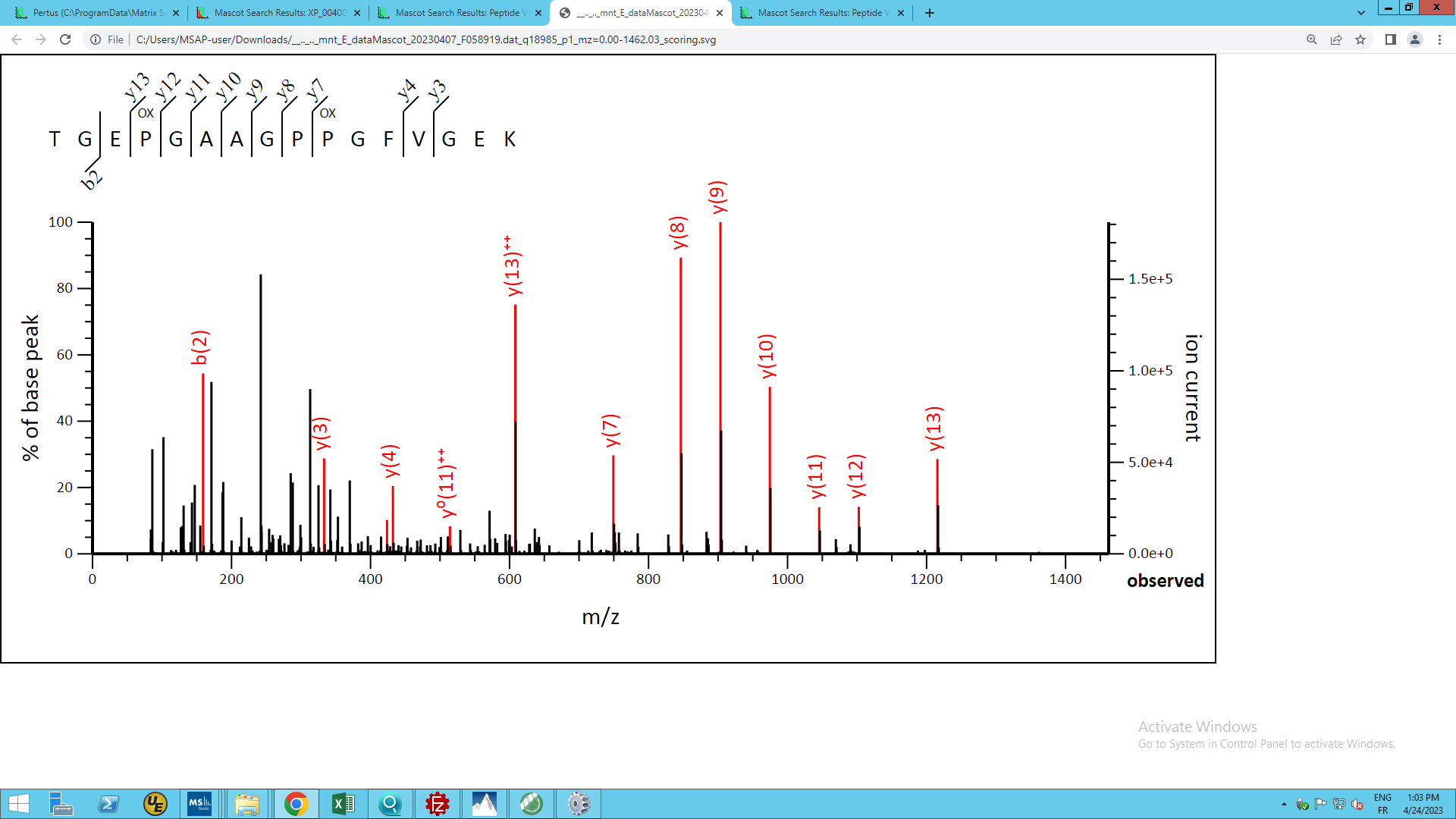

**Supplementary Figure S11.** MS/MS Fragmentation of TGEP^ox^GAAGPP^ox^GFVGEK found in XP_004007775.1 in NCBi_Mamm, collagen alpha-2(I) chain [Ovis aries]

Match to Query 18985: 1501.703648 from (751.859100,2+) scans (17233) rtinseconds (3478.58544) index (12775)

Monoisotopic mass of neutral peptide Mr(calc): 1501.7049

Fixed modifications: Carbamidomethyl (C) (apply to specified residues or termini only)

Variable modifications: P4 Oxidation (P), P10 Oxidation (P), Ions Score: 55 Expect: 8.8e-005

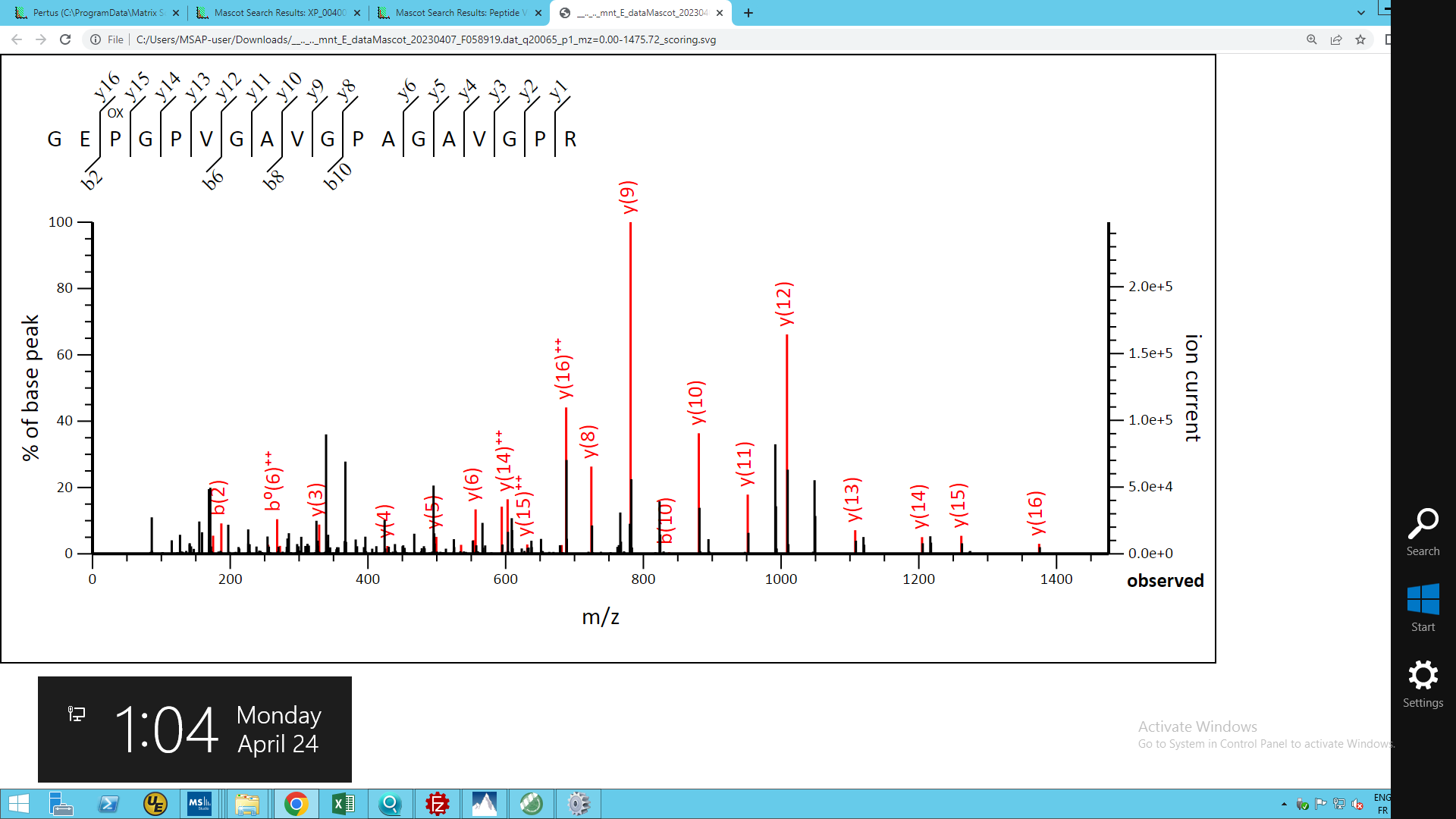

**Supplementary Figure S12.** MS/MS Fragmentation of GEP^ox^GPVGAVGPAGAVGPR found in XP_004007775.1 in NCBi_Mamm, collagen alpha-2(I) chain [Ovis aries]

Match to Query 20065: 1559.804248 from (780.909400,2+) scans (21719) rtinseconds (4177.03578) index (16977)

Monoisotopic mass of neutral peptide Mr(calc): 1559.8056

Fixed modifications: Carbamidomethyl (C) (apply to specified residues or termini only)

Variable modifications: P3 Oxidation (P), Ions Score: 56 Expect: 0.0086

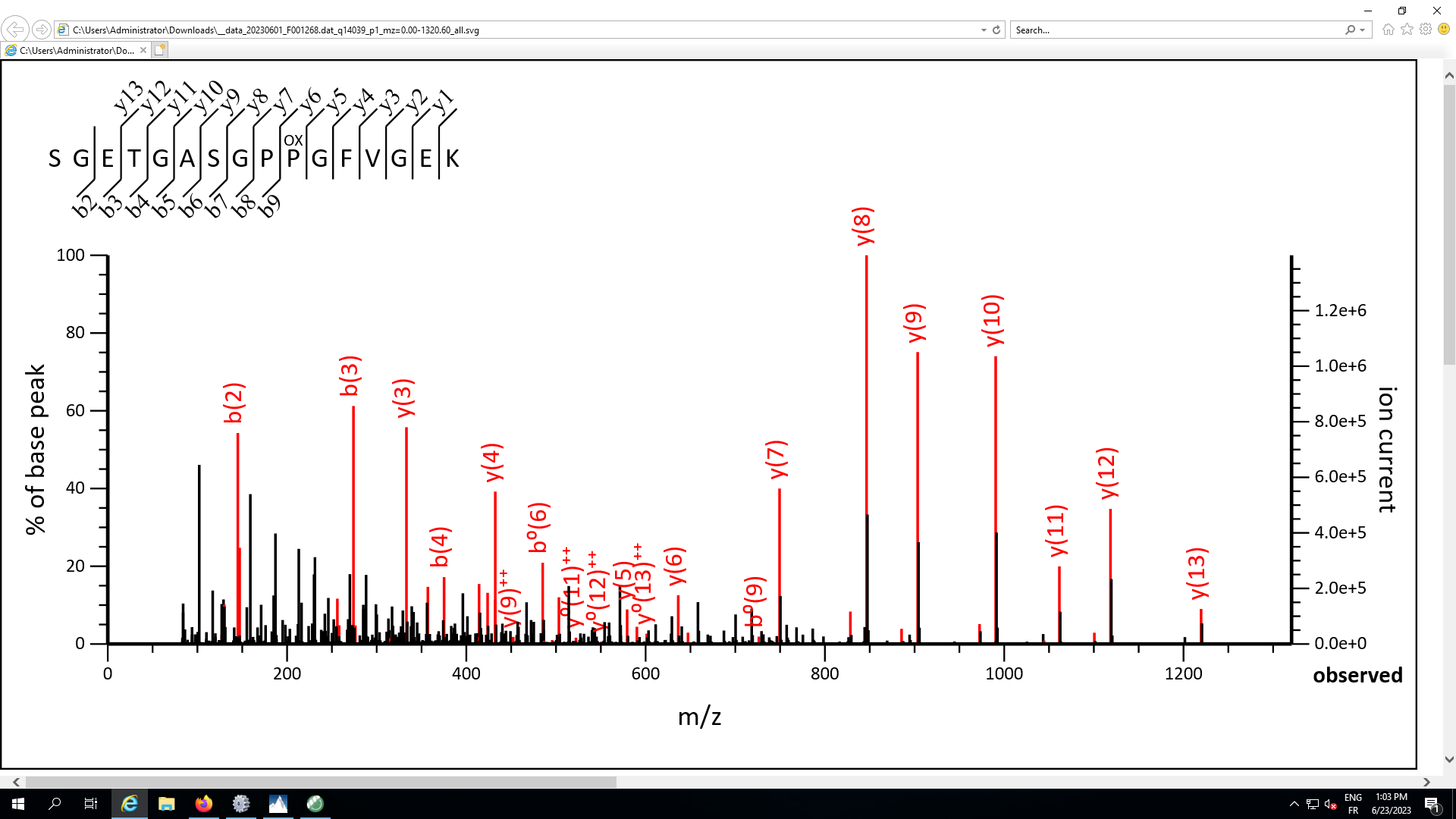

**Supplementary Figure S13**. MS/MS Fragmentation of SGETGASGPP^ox^GFVGEK found in NP_776945.1 in NCBi_Mamm, RecName: collagen alpha-2(I) chain precursor [Bos taurus]

Match to Query 14039: 1491.682620 from (746.848586,2+) intensity (121761818.6875) rtinseconds (3065.95896) index (9366)

Monoisotopic mass of neutral peptide Mr(calc): 1491.6842

Fixed modifications: Carboxymethyl (C) (apply to specified residues or termini only)

Variable modifications: P10 : Oxidation (P), Ions Score: 93 Expect: 8e-08

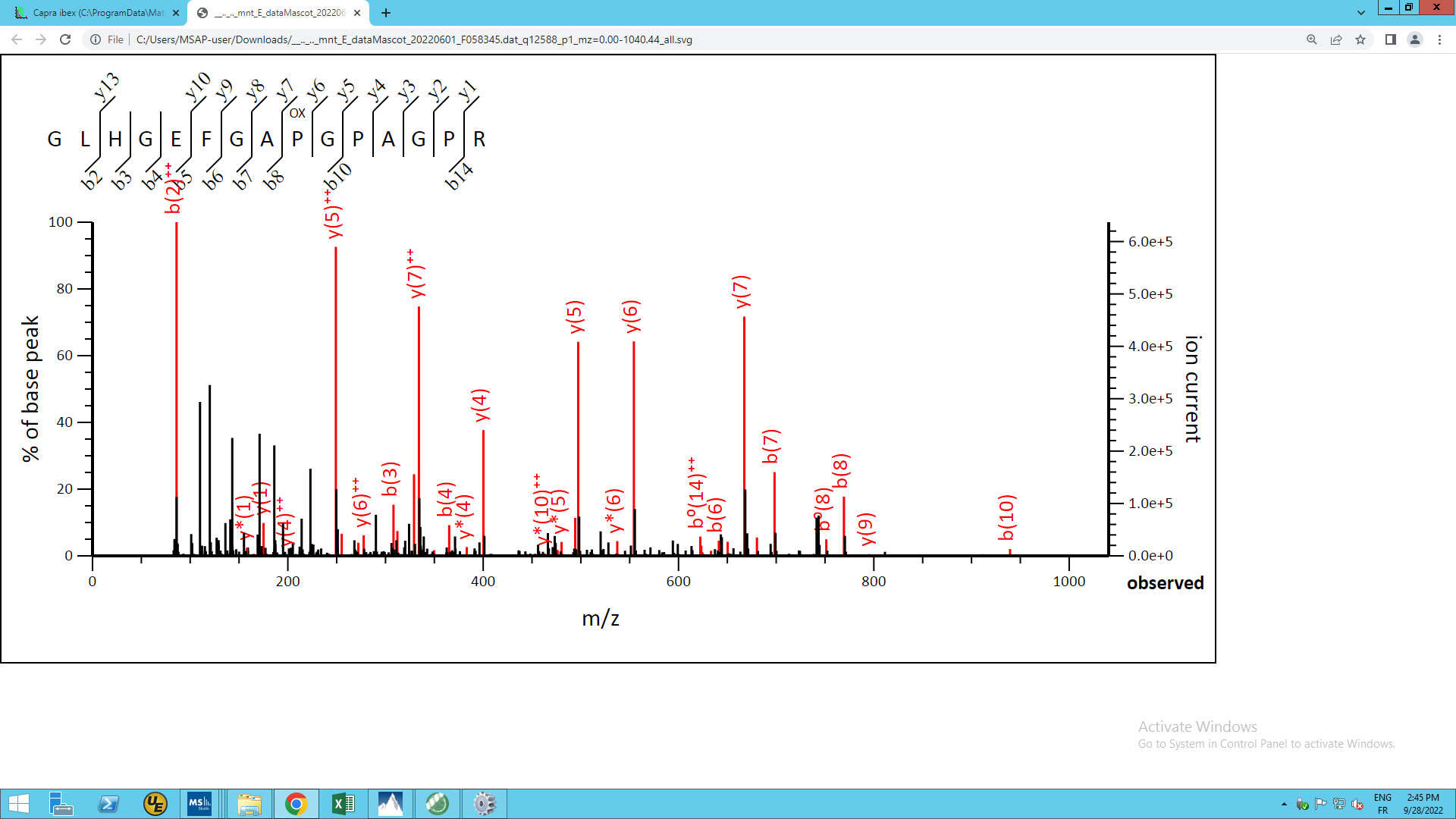

**Supplementary Figure S14.** MS/MS Fragmentation of GLHGEFGAP^ox^GPAGPR found in XP_009672566.1 in NCBI_Aves, PREDICTED: collagen alpha-2(I) chain isoform X1 [Struthio camelus australis]

Match to Query 12588: 1434.702732 from (479.241520,3+) intensity (9234155.0000) rtinseconds (2230) scans (11044) index (7752)

Monoisotopic mass of neutral peptide Mr (calc): 1434.7004

Fixed modifications : Carbamidomethyl (C) (apply to specified residues or termini only)

Variable modifications : P9 : Oxidation (P), Ions Score : 38 Expect : 0.00048

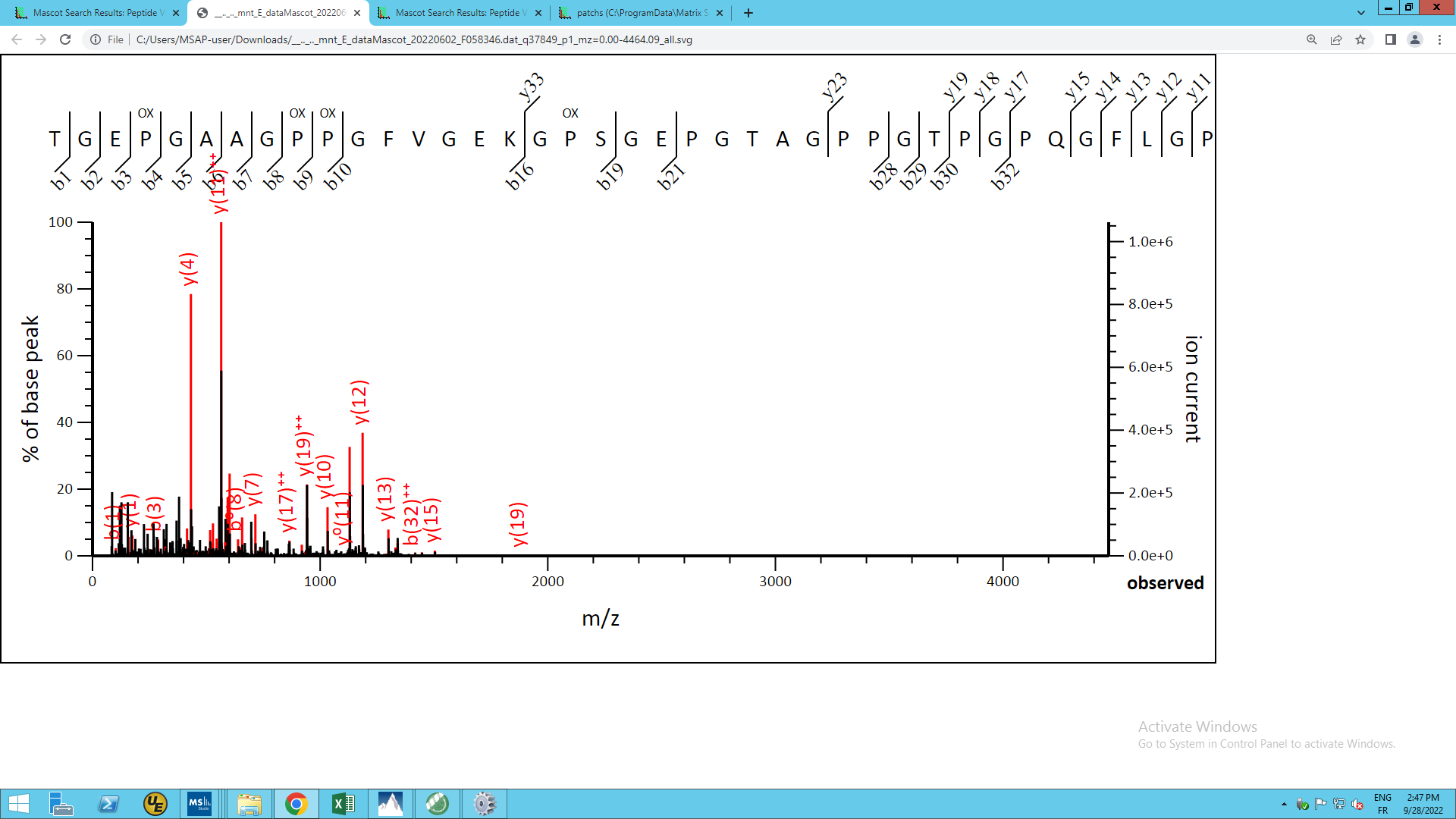

**Supplementary Figure S15.** MS/MS Fragmentation of TGEP^ox^GAAGPP^ox^GFVGEKGP^ox^SGEPGTAGPPGTPGPQGFLGPP^ox^GFLGLP^ox^GSR found in XP_005678993.1 in NCBi_Mamm, PREDICTED: collagen alpha-2(I) chain [Capra hircus]

Match to Query 37849: 4560.212576 from (1141.060420,4+) intensity (2038266.3750) rtinseconds (6806) scans (34309) index (26141)

Monoisotopic mass of neutral peptide Mr(calc): 4560.1835

Fixed modifications : Carbamidomethyl (C) (apply to specified residues or termini only)

Variable modifications : P4 : Oxidation (P), P9 : Oxidation (P), P10 : Oxidation (P), P18 : Oxidation (P), P40 : Oxidation (P), P46 : Oxidation (P), Ions Score : 55 Expect : 1.3e-005

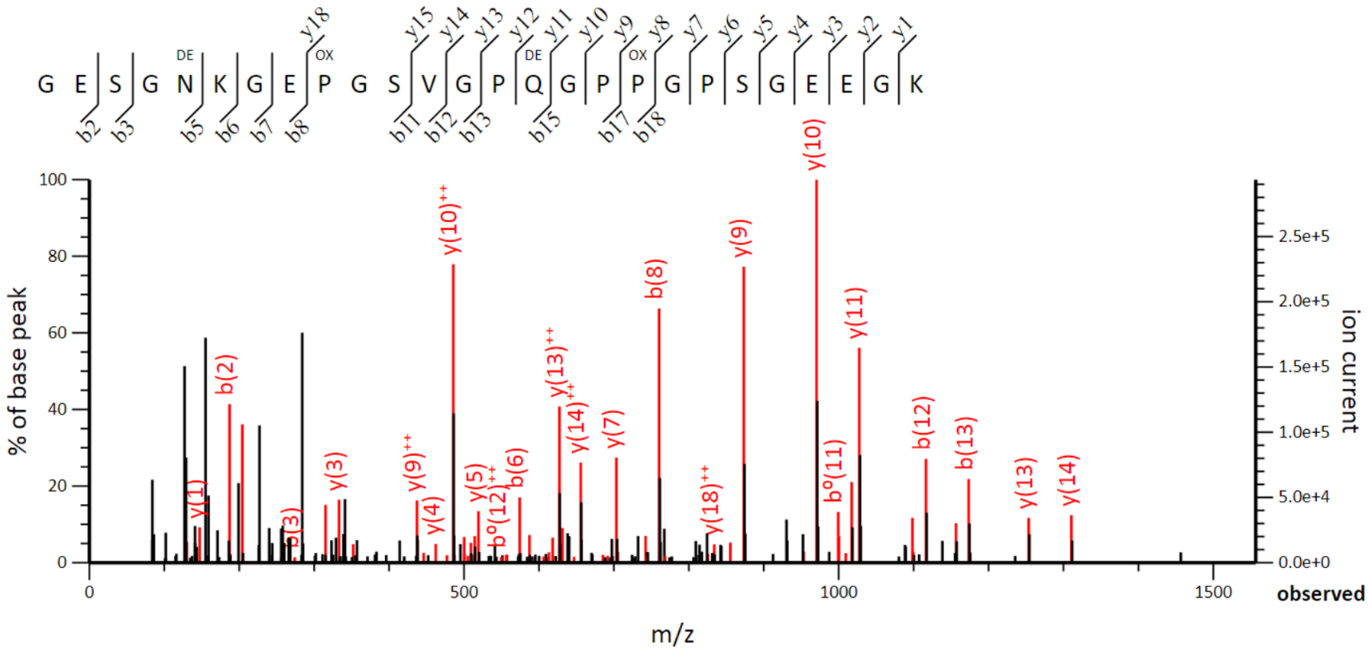

**Supplementary Figure S16.** MS/MS Fragmentation of GESGN^dem^KGEP^ox^GSVGPQ^dem^GPP^ox^GPSGEEGK found in XP_008684476.1 in NCBi_Mamm, PREDICTED: collagen alpha-2(I) chain [Ursus maritimus]

Match to Query 22640: 2425.050612 from (809.357480,3+) intensity (24988886.0000) rtinseconds (2327) scans (9383) index (5089)

Monoisotopic mass of neutral peptide Mr(calc): 2425.0517

Fixed modifications: Carbamidomethyl (C) (apply to specified residues or termini only)

Variable modifications: N5 Deamidated (NQ), P9 Oxidation (P), Q15 Deamidated (NQ), P18 Oxidation (P), Ions Score: 56 Expect: 0.00072
